# Development of covalent probes to capture *Legionella pneumophila* effector enzymes

**DOI:** 10.1101/2024.03.19.585531

**Authors:** Max S. Kloet, Rishov Mukhopadhyay, Rukmini Mukherjee, Mohit Misra, Minwoo Jeong, Cami M. P. Talavera Ormeño, Angeliki Moutsiopoulou, Rayman T. N. Tjokrodirijo, Peter A. van Veelen, Donghyuk Shin, Ivan Ðikić, Aysegul Sapmaz, Robbert Q. Kim, Gerbrand J. van der Heden van Noort

**Affiliations:** Department of Cell and Chemical Biology, Leiden University Medical Centre, Leiden, The Netherlands; Buchmann Institute for Molecular Life Sciences, Goethe University, Frankfurt, Germany; Department of Systems Biology, College of Life Science and Biotechnology, Yonsei University, Seoul, Republic of Korea; Center for Proteomics and Metabolomics, Leiden University Medical Center, Leiden, The Netherlands

## Abstract

Upon infection of host cells, *Legionella pneumophila* releases a multitude of effector enzymes into the cells cytoplasm that hijack a plethora of cellular activities, including the hosts ubiquitination pathways. Effectors belonging to the SidE-family are involved in non-canonical serine phosphoribosyl ubiquitination of host substrate proteins contributing to the formation of a Legionella-containing vacuole that is crucial in the onset of Legionnaires’ disease. This dynamic process is reversed by effectors called Dups that hydrolyse the phosphodiester in the phosphoribosyl ubiquitinated protein. We installed reactive warheads on chemically prepared ribosylated ubiquitin to generate a set of probes targeting these Legionella enzymes. In vitro tests on recombinant DupA revealed that a vinyl sulfonate warhead was most efficient in covalent complex formation. Mutagenesis and x-ray crystallography approaches were used to identify the site of covalent crosslinking to be an allosteric cysteine residue. The subsequent application of this probe highlights the potential to selectively enrich the Dup enzymes from Legionella-infected cell lysates.

## Introduction

Ubiquitination, a crucial post-translational modification (PTM) in eukaryotes, involves attachment of a 76 amino acid-long protein to substrate proteins. This modification plays a vital role in many cellular processes, including proteasomal degradation of proteins and the maintenance of genome stability. ^1–3^ Adenosine diphosphate ribosylation (ADPr) is another PTM that is mainly known for its role in regulating DNA repair processes^4–6^, but that also is widely recognized as signal for ubiquitination.^7^ As such, in addition to the individual impact of both these isolated PTMs on protein signaling, there has been a growing interest in the regulation of ubiquitination by ADPribosylation and vice-versa.^8^ This interest has been particularly fuelled by the discovery of a class of multi-domain *Legionella pneumophila* effector enzymes that physically link these two post-translational modifications together en route to effectively replicate within the host cell.^9,10^ The eukaryotic ubiquitination process begins with the activation of the C-terminus of ubiquitin by an E1 enzyme, which requires energy in the form of ATP. Following the formation of a reactive thioester linkage between E1 and Ub, ubiquitin is subsequently transferred to the E2 and finally, E3 enzymes ligate Ub onto mostly the lysine of target proteins via formation of an iso-peptide bond. In contrast, the ubiquitination mechanism used by Legionella SidE effectors in non-canonical phosphoribosyl (PR)-ubiquitination, starts with ADP-ribosylation of Arg42 of ubiquitin in an NAD^+^ dependent manner^9,10^ mediated by the mART-domain of SidE enzymes. The formed Ub^ADPr^ is subsequently transferred to the phosphodiesterase (PDE) domain of SidEs, which mediate the conjugation to a serine amino acid residue in target proteins resulting in a phosphoribosyl (PR) linkage between ubiquitin and the modified protein substrate (**Supplementary Fig. 1**).^10–13^ Proteome analysis of Legionella infected cells identified proteins predominantly involved in Endoplasmatic Reticulum (ER) fragmentation and membrane recruitment to the Legionella-containing vacuole (LCV) to be PR-ubiquitination targets.^14,15^ Legionella hence gains local control over part of the host-ubiquitinome and is able to orchestrate efficient LCV formation in order to create a favourable environment for bacterial replication. Legionella depends on the SidE effector activities to proliferate inside the host cell, as bacterial replication is significantly hindered without these effectors.^9,16,17^ This process is highly dynamic and regulation of SidE PR-Ub ligase activity by glutamylase SidJ as well as deconjugation of PR-ubiquitination from the host cells proteins by deubiquitinases for phosphoribosyl ubiquitination (Dups) have been identified to control PR-ubiquitination levels. ^14,17–20^ These Dups (DupA and DupB, also named LaiE and LaiF) consist of a single PDE-like domain and perform their action by directly cleaving the phosphomonoester linkage restoring the native host protein and concomitantly releasing phosphoribosyl ubiquitin, a molecule that has been speculated to locally inactivate the conventional host ubiquitination process and acts as an autophagic blockade.^14,18^

Although recent advances in the development of chemical tools, such as small molecule inhibitors^21^, Ub^ADPr^ substrates^22^, stabilised Ub^ADPr^-analogues^23^ and fluorescence-polarization assay-reagents^24^ have demonstrated their value in the study of the PR-ubiquitination pathway, covalently binding probes frequently used to target conventional (de)ubiquitinating enzymes, have not been developed so far to target the PR-ubiquitination machinery.

In this work, we use an adaptable platform based on the copper catalysed click reaction between an azide modified synthetic Ub protein and a variety of propargyl modified ribose molecules that are equipped with reactive groups to generate a small library of probes to target the Legionella Dup family. This method not only gives easy access to a variety of covalent probes but also results in a hydrolytically stable triazole bond between the Ub protein and ribose group, facilitating application of the probes in cellular environments. The hence-created set of probes is subjected to recombinantly expressed DupA to verify covalent binding of the probes and the site of reactivity is investigated using mutagenesis and structural approaches. Subsequent application of the most reactive probe in pull-down experiments from Legionella infected HEK293T cell lysate shows excellent enrichment of the targeted Legionella enzymes highlighting the applicability of our probes to study Legionella effector enzymes in complex biological settings.

## Results

### Chemical synthesis allows introduction of diverse warheads onto Ub

Both the catalytic domains of SidE ligases (PDE-domain) and the deconjugating Dups rely on a triad of Glu-His-His in contrast to the cysteine-based catalysis present in most conventional Ub-ligases and -proteases.^25^ It has been proposed that during cleavage of the phosphodiester bond, the Dups form a covalent phosphoramidate intermediate initiated by nucleophilic attack of His67 on the PR-ubiquitinated substrate thereby liberating the native protein (**Fig. 1 – upper panel**). The subsequent step in the hydrolysis reaction is facilitated by the second reactive histidine of the Dup (His189) that activates a water molecule to attack and hydrolyse the formed Dup(His67)-PR-Ub intermediate (**Supplementary Fig. 2**). The formation of the initial covalent intermediate between the Dup and its substrate opens the opportunity to design covalent probes in analogy to probes carrying Michael acceptor type warheads such as Ub-VME or Ub-PA, used to capture cysteine DUBs.^26,27^ To covalently trap the two members of the Legionella Dup family, we prepared a set of probes that carry a reactive group substituting the phosphodiester group that in a bona-fide substrate links the serine substrate protein to PR-Ub. Upon nucleophilic attack of the Dups’ active site His67 to the probe, a stabile intermediate that cannot be further hydrolysed by the enzyme will result in a permanent covalent bond between probe and enzyme (**Fig. 1A – lower panel**). However, targeting catalytic histidine residues is challenging as the nucleophilicity of this residue is moderate compared to lysine or cysteine. Amongst others, warheads such as fluorosulfonates^28,29^, vinyl sulfonates^30^, vinyl phosphonates^31^ and thiophosphorochloridates^32^ were shown capable to covalently bind surface exposed histidines, although examples of active site histidine reactivity are scarce.^33,34^ Five warheads tailored to react with the mild nucleophilic histidine residue were selected and installed at the 5’-hydroxyl of the ribosyl group; vinyl phosphonate **1**, vinyl ethoxy phosphonate **2**, fluorosulfonates **3** and **4** and vinyl sulfonate **5**. After synthesis of the ribosides (**Supplementary Scheme 1** and **2**) equipped with the diverse warheads at the 5’-hydroxyl and a propargyl moiety installed at the 1’-hydroxyl we used copper catalysed click chemistry to conjugate full length ubiquitin carrying a rhodamine fluorophore on the N-terminus and an Arg42 to azido-homoalanine mutation to each of the ribosides, resulting in five fluorescently labelled covalent probes (**Fig. 1A** – lower panel).^35^

**Fig. 1.**
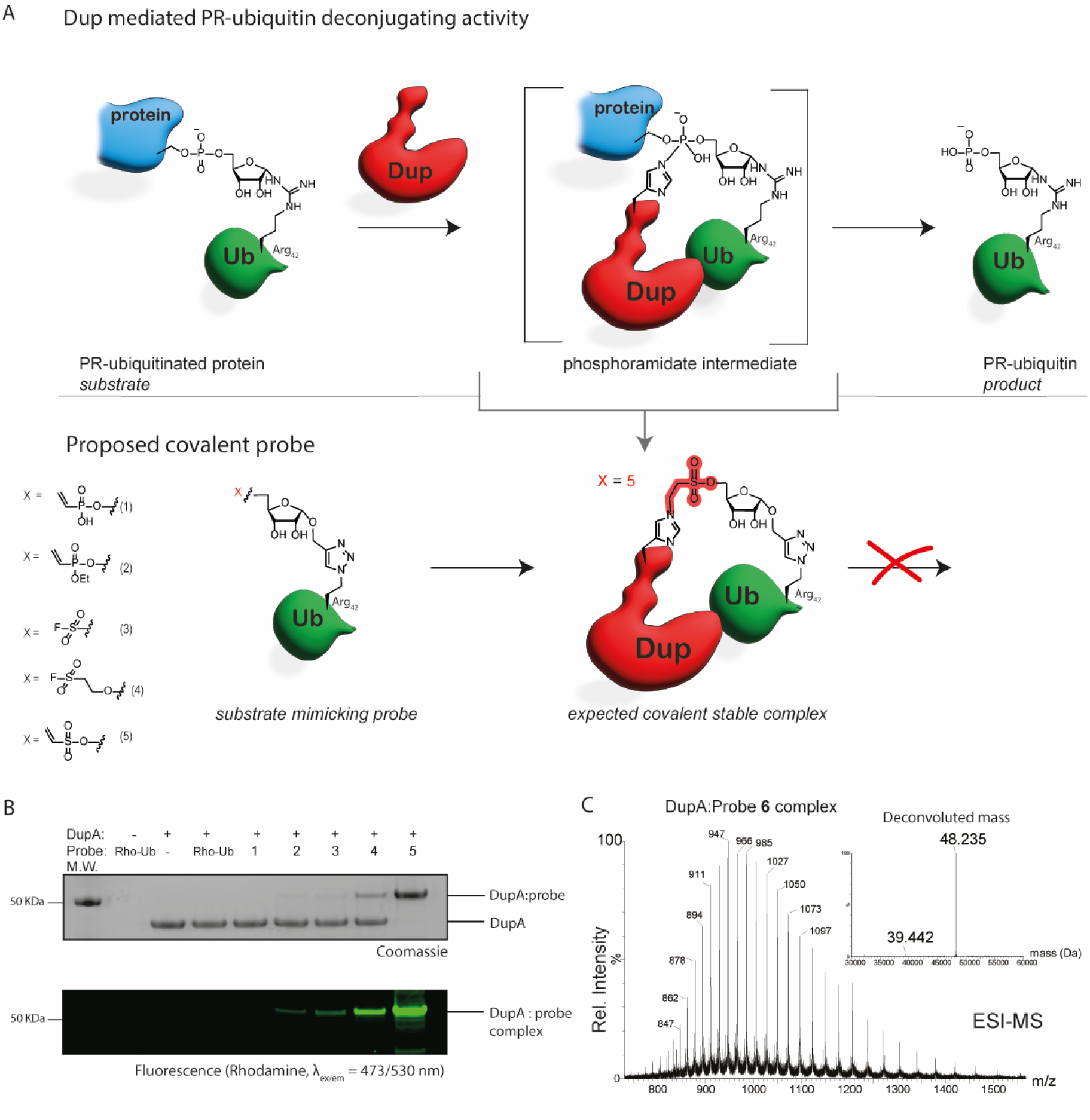
Assessment of the reaction between probe library **1-5** and recombinant DupA WT. **A)** Schematic representation of part of the catalytic mechanism of Dup activity showing the covalent phosphoramidate intermediate that Dup-His67 forms when deconjugating a Ser-PR-ubiquitinated substrate (upper panel) and the expected chemical trapping of the covalent transition state intermediate by our substrate mimicking probe (lower panel). The different warheads installed on the probes **1-5** are depicted on the left. The stable covalent complex formed impedes hydrolysis to the PR-Ub product. **B)** SDS-PAGE gel showing complex formation between probes **1-5** with DupA WT. Probe **1-5** (8 equivalents) were subjected to DupA and incubated at 37 °C for 2 hours and analyzed by SDS-page: Coomassie stain (upper panel), Rhodamine fluorescence scan (λ_ex/em_ = 473/530 nm-bottom panel). **C)** Mass spectrometry analysis of DupA:probe **6** complex, ESI MS of the conjugate reacting vinyl sulfonate probe **6** (deconvoluted mass: 8.793 Da) to DupA wt (deconvoluted mass: 39.442 Da). Deconvoluted mass of complex (found: 48.235 Da, calculated: 48.237 Da) confirming the covalency of the conjugate.

### Chemical probes react covalently with recombinantly expressed DupA

The reactivity of probes **1**-**5** was evaluated by incubating them with recombinantly expressed DupA for two hours at 37 °C (**Fig. 1B**). Both vinyl phosphonate **1** and its ethoxy variant **2** showed little to no complex formation based on in-gel fluorescence and Coomassie staining, in-line with literature reports indicating such vinyl phosphonates to be low in reactivity.^31^ Additionally, the fluorosulfonates **3** and **4** showed moderate complex formation, with the longer spacer probe **4** showing slightly better results than its shorter analogue **3**. Gratifyingly, the vinyl sulfonate probe **5** quickly reacted and reached full conversion even after shorter incubation times of only fifteen minutes (**Supplementary Fig. 3**). To further investigate the labeling efficiency of probe **5**, we performed a time series on ice to get an indication of the rate of binding of our probe, observing close to full conversion as soon as five minutes with five equivalents of probe (**Supplementary Fig. 4**). We hence concluded the vinyl sulfonate warhead used in probe **5** to be our most reactive probe and we applied this warhead in the rest of our studies.

To verify the covalent nature of the formed complex we performed high resolution intact protein MS analysis using a non-tagged version of the vinyl sulfonate probe **6**, to exclude influence of the rhodamine dye on the binding event. The deconvoluted mass of the formed complex (m/z = 48.235 Da) corresponds with the calculated mass of the DupA:probe complex (m/z = 48.237 Da) and verifies the covalent nature of the linkage (**Fig. 1C**). To provide a first indication of selectivity of our probe to Legionella effector enzymes we tested reactivity of the vinyl sulfonate **5** to conventional cysteine family deubiquitinating enzymes OTUB2 and UCHL3 and found no reactivity towards both canonical DUBs. Additionally, NAD^+^ consuming enzymes NMNAT1 and ART1 were examined and probe **5** was unable to covalently react with either of these enzymes (**Supplementary Fig. 5**).

### DupA mutational analysis reveals unexpected site of reactivity

To confirm that our PR-Ub probe **5** binds in the active site of DupA and acts as a suicide inhibitor, we first incubated DupA with probe **5** to form a complex, followed by the addition of Ub^ADPr^, a DupA substrate. DupA activity was visualized using mass spectrometry analysis by the appearance of Ub^Pr^ due to the hydrolysis of the pyrophosphate bond in substrate Ub^ADPr^. Indeed, Ub^ADPr^ was not cleaved by the DupA: probe **5** complex, whereas in the absence of pre-incubation with probe **5**, Ub^ADPr^ hydrolysis mediated by DupA is swiftly observed (**Supplementary Fig. 6**). This indicates that probe **5** abrogates catalytic activity or blocks the active site. In order to pinpoint the exact site of covalent modification, we performed mutational analysis where we mutated His67 or His189 to Ala and unexpectedly observed a retained reactivity of both mutants to probe **5** (**Fig. 2A**). We speculated that when one of the active site histidines was mutated the probe might react with the remaining histidine, that is situated in close proximity. However, even the DupA double mutant (His67Ala/His189Ala) was significantly labelled by probe **5**. When examining the catalytic triad of DupA, the crystal structure of Ub interacting with DupA (PDB:6RYA) revealed that the Cys196 resides in the same helix as His189 and is in close proximity to both active site His67 and His189. Although Cys196 is not part of the catalytic triad and placed slightly outside of the active site, we wondered whether this cysteine might react with probe **5**, as the vinyl sulfonate warhead is a Michael acceptor that could also react with cysteine. Indeed, despite some minor residual labelling activity, reacting probe **5** with the Cys196Ala mutant significantly reduced the formation of the DupA-probe complex. Likewise, treating DupA with a cysteine-alkylating agent (iodoacetamide) prior to incubation with probe **5** abrogates labelling (**Supplementary Fig. 3C**). Next we subjected the formed DupA:probe **5** complex to trypsin digestion followed by tandem MS analysis. When comparing non-treated DupA WT versus DupA:probe **5** complex a change in peptide fragments identief was observed (**Supplementary Fig. 7A** and **7B**). The tryptic peptide fragments that were absent in the latter case might indicate probe **5** covalently crosslinking itself to an amino acid residue in that particular peptide thereby modifying it and hindering its identification. Indeed, the tryptic peptide fragment harbouring Cys196 (C196K197) was observed for DupA wt and was no longer present after reacting with probe **5**. No alterations were observed in the peptide coverage of both the regions harbouring the active site His67 or His189 residues. The peptide belonging to the Ub-remnant (Ub_34-48_) crosslinked to the DupA_196-197_ peptide was subsequently identified (**Supplementary Fig. 7** and **8**).

**Fig. 2.**
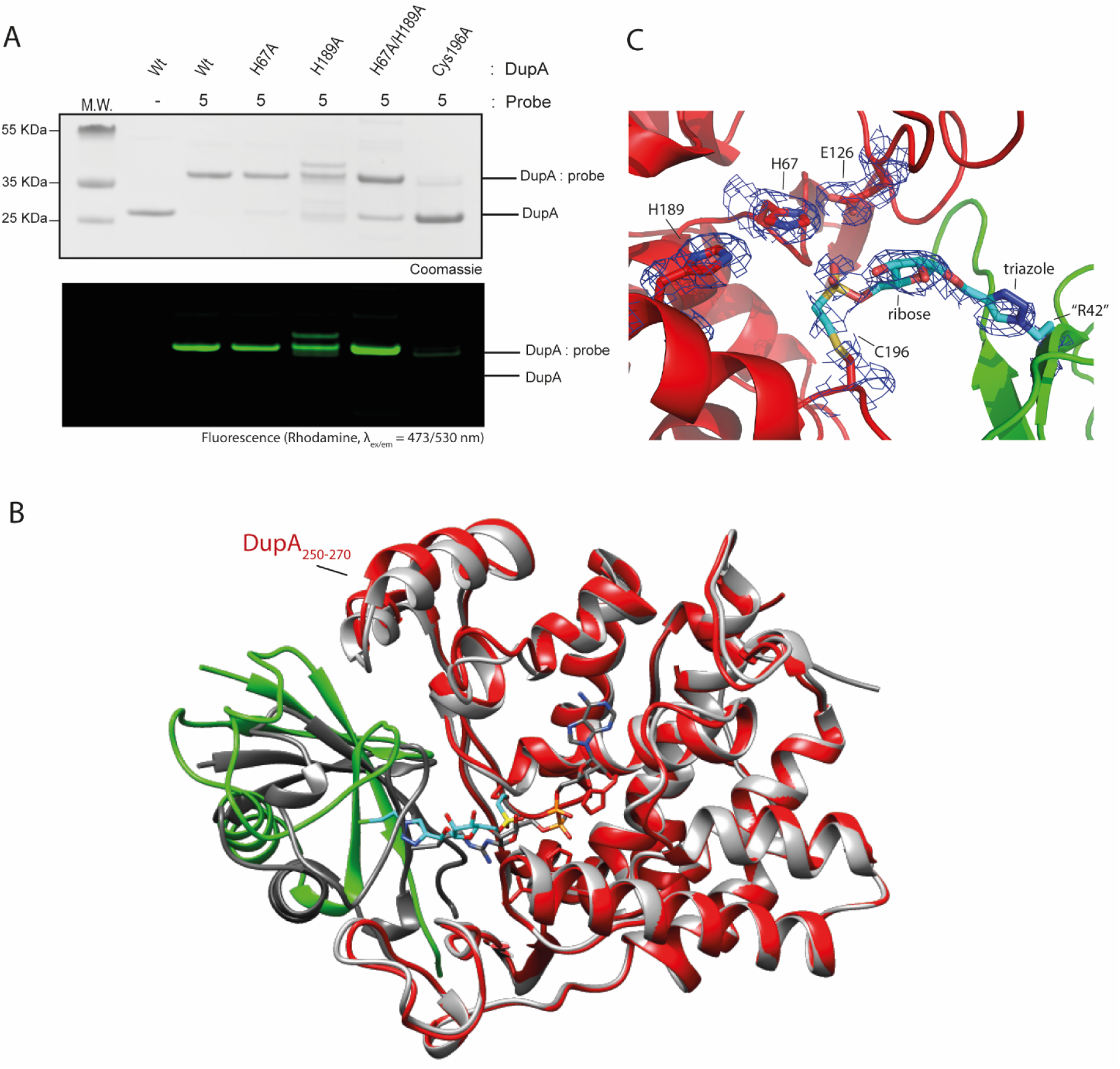
Mutagenesis and crystallographic investigation of covalent DupA:probe complex. **A)** Mutagenesis of DupA and reactivity of the mutants to rhodamine labelled vinyl sulfonate probe **5. B)** Overview of the DupA:probe **6** complex crystal structure (in red and green respectively; PDB: 9EMK), compared to DupA-His67Ala:UbADPr complex (PDB: 6B7O; in grey and dark grey respectively). **C)** Structure and electron density of crosslinked region (cyan) between DupA (red) and Ub probe **6** (green).

### X-ray crystallography confirms reactivity to Cys196

To further consolidate our findings that our probe covalently reacts with Cys196 of DupA, we formed the complex between DupA and probe **6** (the non-tagged version of probe **5**) and were able to obtain diffracting crystals (PDB: 9EMK) (**Supplementary Table 1**). With the P3_2_ space group and the inability of direct molecular replacement with PDB model 6RYA^13^ (space group: C2), we had to resort to do molecular replacement using apo DupA (PDB: 6RYB) and manually place Ub (PDB: 1UBQ) to solve the structure. This still yielded an asymmetric unit with 3 DupA complexes, albeit differently arranged through contacts. Upon closer inspection, the DupA monomer compares very well to those in structures of DupA-apo (PDB: 6RYB), DupA^His67Ala^-Ub (PDB:6RYA) and DupA^His67Ala^-Ub^ADPr^ (PDB:6B7O) with RMSD values <0.4 Å (**Supplementary Fig. 9A**). The main difference is in the DupA alpha-helical lid (a.a. 250-270) that protrudes slightly and makes contact with the bound ubiquitin (**Fig. 2B**). Indeed, when comparing the ubiquitin moiety between the Ub-bound structures (6RYA, 6B7O) we can observe a small tilt and movement for Ub probe **6**, making a more snug interaction with the aforementioned loop_250-270_. The most likely reason for this movement is found at the binding site, where the reactive warhead has inserted itself towards active site His67 (**Supplementary Fig. 9B**), but is caught and bound by Cys196 before reaching the target (**Fig. 2C**). This creates some sterical repulsion because of the local strain in the sulfonate probe when ubiquitin would try and nest itself in the actual binding site, so instead it now binds loop_250-270_, albeit with some flexibility. This flexibility is also seen in the associated B-factors as well as how well the connecting warhead is defined in the electron density. The found density nevertheless further confirms our MS-MS data (**Supplementary Fig. 7** and **8)** and mutational analyses (**Fig. 2A**) and shows the vinylsulfonate probe binds Cys196 of DupA.

### The vinylsulfonate probe is able to enrich both DupA and DupB from Legionella infected HEK293T cell lysate

As probes **5** and **6**, both equipped with the vinylsulfonate warhead, label recombinant DupA with high reactivity, we set out to validate the potential of such a probe to enrich Dups from the lysate of human HEK293T cells infected with Legionella. To facilitate the pulldown and allow for efficient visualization in the optimization process, we prepared vinylsulfonate Ub probe **7**, a probe similar to **5** (and **6**) but equipped with both a rhodamine-fluorophore and biotin-moiety. We used azido-Ub (the Arg42 to azido-homoalanine mutant, not functionalised with the vinysulfonateribosyl moiety) and DMSO treatment as controls, respectively (**Fig. 3A**). Both non-infected HEK293T cells and cells infected with Legionella were lysed four hours post-infection as the optimal time point for detecting the activity of the Dups.^14^ The lysates were incubated with probe **7** and the appropriate controls at 37 °C for two hours (optimization shown in **Supplementary Fig. 10**). Neutravidin beads were added to capture the biotin-Ub-probe-protein conjugates. After stringent washing with buffer containing 2% SDS to remove all non-covalently interacting proteins, the beads were subjected to elution followed by trypsin digestion and mass-spectrometric analysis of the proteome. When comparing proteins enriched by probe **7** in the infected cells compared to non-infected cells, both DupA and DupB are significantly enriched (**Fig. 3B**). When comparing probe **7** and the biotin-Ub control within the infected sample group, DupA and DupB again show significant enrichment (**Fig. 3C**). Comparing azido-Ub versus DMSO (**Supplementary Fig. 11**) showed no significantly enrichment of either Dup, highlighting the necessity of the warhead leading to the formation of the covalent complex between the probe and the Dups under Legionella infected conditions as well as underscoring that non-covalent interaction with Ub is not enough for enrichment of these Dups. Overall, probe **7** shows remarkable specificity towards DupA and DupB since they were specifically enriched among the 2930 distinct proteins in the Legionella proteome.^36^ Upon further analysis of the data we also identified a change in the subset of mammalian (human) proteins captured by probe **7** between infected cells and non-infected cells (**Fig. 3B**). Besides Legionella proteins DupA and DupB we observed significant enrichment for mammalian PARP1 and MacroD1. Interestingly, we observed both the PARP1 and MacroD1 enrichment in the left panel, indicating a marked decrease of protein expression and/or reactivity during Legionella infection. PARP1 has been described to play a role in the immune response during *Salmonella enterica* and *Serovar Typhimurium* infection where it regulates NF-κB-mediated pro-inflammatory gene expression.^37,38^ The role of PARP1 during Legionella infection, however, has not been clarified to date and decreasing activity of PARP1 by the bacterium potentially could affect the initiation of the immune response during infection. The other hit is MacroD1, an ADPribose glycohydrolase, and its decreased reactivity during Legionella infection might suggest that Legionella hinders ADP-ribose hydrolysis to promote its bacterial virulence. Another notable observation is the enrichment of DNA damage response proteins XRCC5 and XRCC6 in the left panel (**Fig 3B**).^39,40^ XRCC5 and XRCC6 are shown to be essential in regulation of the innate immunity response upon viral infections and it could therefore well be that in analogy Legionella acts on these proteins to suppress the host immune response.^41^ Additionally, PARP1 as well as XRCC5 and XRCC6 have been identified prior as potential PR-ubiquitination targets.^18^

**Fig. 3.**
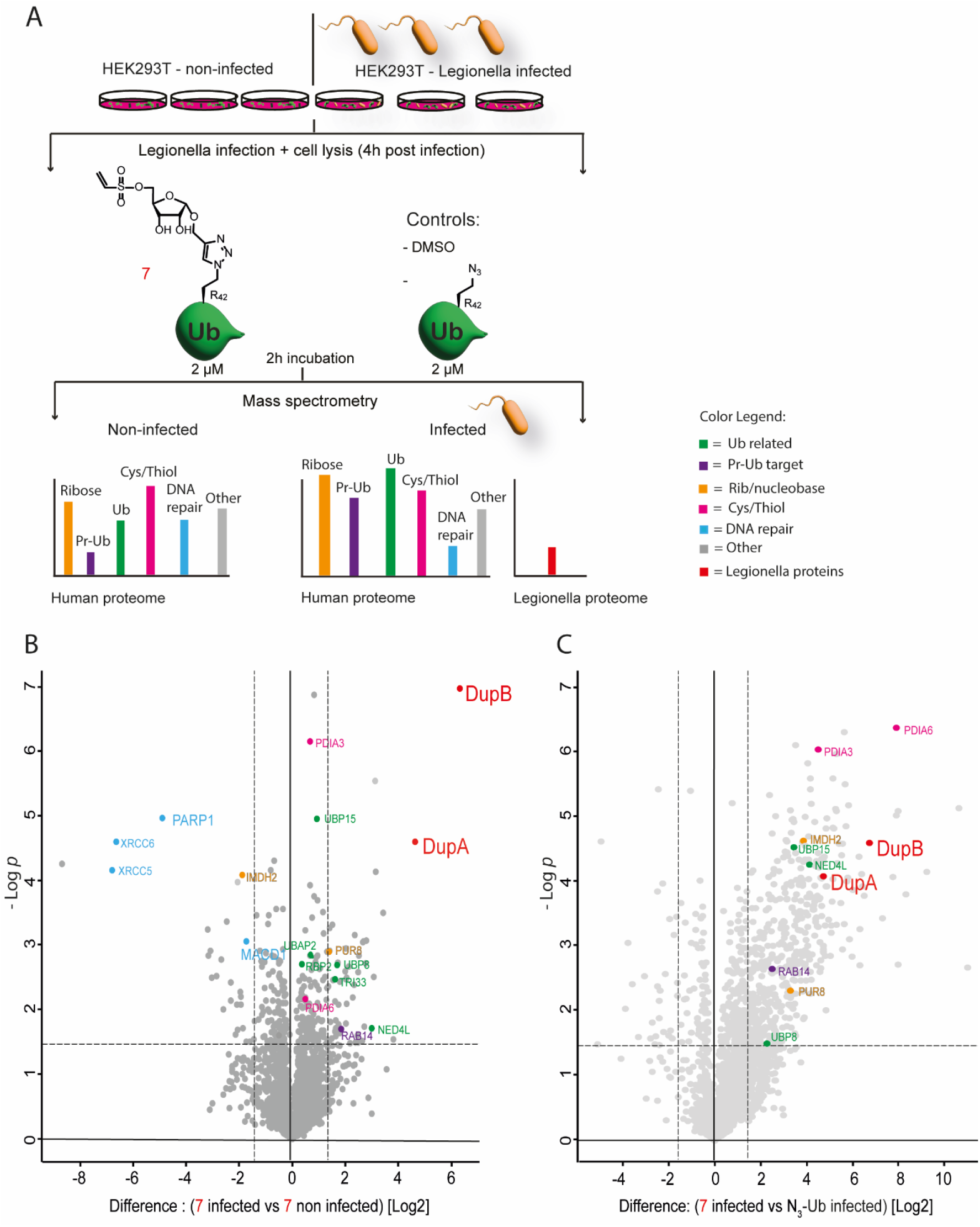
Chemoproteomic assessment of pulldown by the vinylsulfonate probe **7** from non-infected HEK293T cells or cells infected with Legionella, **A)** Schematic representation of the workflow applied in mass spectrometry-based proteomics. The lysates of two sample groups; infected with Legionella and non-infected, were prepared in triplicate. **B)** Volcano plot depicting proteins enriched by probe **7** comparing the infected sample group to the non-infected sample group (dotted line represents significance), **C)** Volcano plot showing the significant enriched proteins by probe **7** compared to the biotin-Ub control within the infected sample group (dotted line represents significance). For every plot holds: Black dotted lines correspond to the thresholds: log_2_ ratio ≥1.5; *p*-value ≤0.05. A color code legend is provided for the clustered proteins. The Legionella enzymes DupA and DupB are marked in red.

Upon closer examination of the proteins we enriched in the human proteome, we classified the identified proteins in different clusters, based on their respective roles in regulating Ub dynamics (green), previously reported SidE-targets (purple), potential reactivity towards the warhead (pink) and recognition of structural properties of the ribosyl moiety (orange) (**Fig. 3B**,**C, Supplementary Fig. 11**). Since our probe has a reactive warhead that captures nucleophilic amino acid residues, we are not surprised to find clusters of proteins that potentially recognize either the Ub-core or the phosphoribosyl-mimic attachment as substrate and subsequently covalently react with the probe in the two hour incubation period. Ub-conjugating enzymes like NEDD4L and TRIM33 and Ub proteases, including ubiquitin-specific proteases (USPs); USP3, USP14, USP48, OTUB1 and USP15 were amongst the enriched human proteins that show reactivity towards the probe (**Supplementary Fig. 11, Fig. 3B and C**). The activity of the conventional ubiquitin machinery is known to be manipulated by bacterial infection, and therefore, it might be interesting to further investigate these Ub-ligases and DUBs that show a change in reactivity towards our probe upon infection by the Legionella bacterium (in green; **Fig. 3B**).

Another cluster of proteins enriched by our probe, including PUR8 and IMDH2 (orange), might be captured due to their recognition of the modified ribose moiety in the probe. PUR8 for instance is an adenosylsuccinate lyase and therefore likely recognizes the sulfonate-ribose functionality in probe **7**. General cysteine reactivity towards the Michael acceptor warhead in our probe might explain the enrichment of PDIA3 and PDIA6 (pink) as these enzymes are disulfide isomerases relying on a Cys-Gly-His-Cys active site motif.^42,43^

We further annotated proteins previously implicated to be PR-ubiquitination targets in purple.^14,18^ Some of these include Ub conjugating enzymes UBR5 and UBA1 and deubiquitinases such as USP5 and USP10, but we also enrich MAP4, CCAR2, PRDX1, ACLY, ATXN2L and EIF3D and several Rab GTPases (RAB14 (**Fig. 3B**,**C**), RAB5C, RABL6 and RAB21 (**Supplementary Fig. 11**)) some of which are known substrates for PR-ubiquitination. These GTPases are involved in membrane trafficking and/or the endocytic pathway and by manipulation of these proteins the bacterium might try to improve stability of the LCV.^44^

The use of the vinylsulfonate probe warrants covalent crosslinking to a significant number of proteins in the lysates and although some of these captured proteins may be off-targets due to the high reactivity of the warhead, the intended Dup family is amongst the most significant enriched proteins. In addition comparison of non-infected and Legionella infected cells leads to some interesting changes of protein reactivity towards our probe that might give insights in how the bacterium manipulates the host cell.

## Discussion

Legionella manipulates its host cell by releasing a wide variety of effector enzymes. Among these effectors, the SidE enzyme family plays a crucial role in deregulating Ub signaling by effecting PR-ubiquitination thereby favoring the replication of Legionella in the Legionella-containing vacuole. This PR-ubiquitination process itself is tightly regulated by SidJ enzymes and reversed by Dup effectors. Measuring or interfering with these different enzyme activities will help to gain fundamental molecular insights into these crucial steps in Legionella infection. We hence developed a chemical methodology for the generation of a collection of ubiquitin phosphoribose analogues, which serve as covalent probes to target Legionella Dup effectors. We successfully generated a highly reactive vinyl sulfonate probe able to covalently react with recombinant DupA, achieving fast and complete conversion. Although this vinyl sulfonate warhead was initially designed to target the catalytic histidine of the Dups, our studies indicate main reactivity of probe **5** towards the Cys196 residue. Structural analysis indeed confirmed covalent crosslinking of the non-catalytic Cys196 forming a stable link between DupA and the PR-Ub derived probe. Although the targeted cysteine is a non-catalytic residue the location near the active site might make this residue an interesting target for future drug development as reacting this cysteine residue with the bulky Ub-based probe but also the small alkylating reagent iodoacetamide reduced catalytic activity. We further successfully enriched both DupA and DupB from Legionella-infected HEK293T lysate, in a pulldown-proteomics experiment, with the protease stable triazole linked probe **7**, confirming the applicability of this probe in a complex proteome. Of note, we did not enrich the SidE effectors in this pulldown experiment and although the PDE domains of SidEs have a high strucutural homology to the Dups, the Cys196 from the Dups is not conserved in the SidEs. In addition the Legionella-infected cells used in our study were lysed four hours post-infection and previous research indicates that the activity of SdeA is concentrated in the initial period after infection and is significantly reduced four hours post-infection.^45^ Recent interest in regulatory mechanisms involving ADPribosylation of Ub or vice versa both in bacteria^17,19,20,46,47^ and mammals^48–51^ has grown and the tools presented here could be adapted and applied in a broader context. Consequently, further investigation of the Ub and ADPr interplay in human cells and the utilization of the probes presented here could help to unveil the molecular mechanisms involved in both normal homeostasis as well as in bacterial infection conditions.^8^

## Methods

### Chemical synthesis of the probes and small molecule ribosides is described in detail in the Supplementary Information

#### Constructs and protein purification

The bacterial expression construct for DupA has been described previously.^13^ The mutations (H67A, H189A, the combination and C196A) were generated using overlapping primers^52^ and all were sequence-verified before expression. The N-terminal GST-tagged DupA expression constructs were transformed into T7 Express cells and cultured at 37 °C in LB supplemented with 100 μg mL^-1^ ampicillin. When OD_600_ reached 0.6, expression was induced using 0.5 mM IPTG (Isopropyl β-D-1-thiogalactopyranoside) and temperature was lowered to 20 °C for 5 hours.

Harvested cell pellets were resuspended in GST buffer (50 mM Tris pH 7.5, 150 mM NaCl and 1 mM TCEP) and sonication on ice, before being spun down at 24 kG for 40 minutes at 4 °C. The resulting supernatant was applied to GST 4B Sepharose beads, followed by extensive washing with GST buffer and DupA elution using elution buffer (50 mM Tris pH 7.5, 50 mM Nacl, 25 mM GSH, 1 mM TCEP). Overnight incubation with TEV protease resulted in cleaved DupA and the protein solution was concentrated using spin filtration before a final purification step over a Superdex 75 16/60 column, equilibrated in a final buffer containing 20 mM HEPES pH 7.5, 100 mM NaCl and 1 mM DTT. All steps were analysed on SDS-PAGE and the final product was conctrated to ∼10 mg mL^-1^ before being flash frozen in liquid nitrogen.

#### Labeling assay on recombinant DupA: Reactivity of the probes (1-5)

The probes **1**-**5** (27 μM, 8 eq.) in buffer (20 mM TRIS, 150 mM NaCl, pH 7.6) were incubated with DupA WT (3.4 μM) at 37 °C in a total volume of 26.7 μL. After 15 min/ 2 h (the indicated time points on SDS page **Supplementary Fig. 3**) of incubation 10 μL sample was taken and added to 5 μL loading buffer including β-mercaptoethanol. The samples were run on a NuPAGE™ 12% Bis-Tris gel in MES buffer, 190 mV, for 45 minutes. A fluorescence scan on a Typhoon FLA 9500 (rhodamine channel, λ_ex/em_ = 473/530 nm) was performed to visualize the complex formed and additionally, the proteins were stained with Coomassie staining.

#### Labeling assay on recombinant DupA; time series on ice

Probe **5** (22 μM, 5 eq.) in buffer (20 mM TRIS, 150 mM NaCl, pH 7.6) was incubated with DupA WT (4.4 μM) which was kept on ice, in a total volume of 20 μL. The labelling reaction was performed at 0 °C and monitored at the indicated time points (0.5, 1, 2 and 5 min). 5 μL of each sample was mixed with loading buffer including β-mercaptoethanol (10 μL) and subsequently ran on a NuPAGE™ 4-12% Bis-Tris gel in MES buffer, 190 mV for 45 minutes. A fluorescence scan on a Typhoon FLA 9500 (rhodamine channel, λ_ex/em_ = 473/530 nm) was performed to visualize the complex formed and additionally, the proteins were stained with Coomassie staining.

#### Labeling assay on recombinant DupA; generating the DupA-vinyl sulfonate probe complex, which upon full conversion is incubated with Ub^ADPr^

Probe **5** (27 μM, 8 eq.) in buffer (20 mM TRIS, 150 mM NaCl, pH 7.6) was incubated with DupA WT (3.4 μM) in a total volume of 26.7 μL. The reaction was incubated at 37 °C and after 2h LC-MS indicated full conversion into the DupA-probe complex (M + H^+^ = 48.595). Ub^ADPr^ (24.6 μM, 7.3 eq.) was added and the conversion of Ub^ADPr^ to Ub^Pr^ was monitored by LC-MS after 15 min and 30 min of incubation. The same reaction procedure was performed in parallel preincubating DupA wt (3.4 μM) with Rhodamine-Ub (27.4 μM, 8.1 eq.) instead of probe **5**, for 2 hours at 37 °C. Subsequently, Ub^ADPr^ (24.6 μM, 7.3 eq.) was added and the Ub^ADPr^ to Ub^Pr^ conversion monitored by LC-MS. The mass spectrometry analysis is depicted in **Supplementary Fig. 6**.

#### Labeling assay of vinyl sulfonate probe 5 to recombinant OTUB2, UCHL3, NMNAT1 and ART-1

Probe **5** (27 μM, 8 eq.) in buffer (20 mM TRIS, 150 mM NaCl, pH 7.6) was incubated with one the enzymes (OTUB2, UCHL3, NMNAT1 or ART-1, 3.4 μM) at 37 °C in a total volume of 26.7 μL for the indicated time points. For the DUBs (OTUB2 and UCHL3) reactivity towards Rhodamine-Ub-Prg (27 μM, 8 eq.) was taken along. After the incubation, 10 μL sample was taken and added to 5 μL loading buffer including β-mercaptoethanol. Samples were ran on a NuPAGE™ 4-12% Bis-Tris gel in MES buffer, 190 mV for 45 minutes. A fluorescence scan on a Typhoon FLA 9500 (rhodamine channel, λ_ex/em_ = 473/530 nm) was performed to visualize the complex formed and additionally, the proteins were stained with Coomassie staining. The SDS PAGE analysis is depicted in **Supplementary Fig. 5**.

#### DupA-probe complexation for structural analysis

Vinyl sulfonate probe **6** (77 μM, 1.4 eq.), which is the un-tagged probe variant, in buffer (20 mM TRIS, 150 mM NaCl, pH 7.6) was incubated with DupA WT (57 μM) in a total volume of 777 μL. The reaction was shaken at 37 °C and monitored by LC-MS. After 30 min, additional probe **6** (48 μM, 37 μmol, 0.8 eq.) was added. Full conversion was reached after 1 hour (M + H^+^ complex = 48.235 Da) and the complex was purified by gel filtration using a Superdex75 10/300 column equilibrated in 20 mM HEPES pH7.5 and 100 mM NaCl.

#### Cell lines

HEK293T expressing CD32 were cultured in DMEM supplemented with 10% FBS, 100 I.U./mL penicillin and 100 mg/mL streptomycin (Pen/Strep) at 37 °C, 5% CO_2_.

#### Legionella pneumophila culture and infection

Wild type L. pneumophila (Lp02) was grown for 3 days on N-(2-acetamido)-2-amino-ethanesulfonic acid (ACES)-buffered charcoal-yeast (BCYE) extract agar, at 37 °C, followed by growth for 20 h in ACES yeast extract media. Bacterial cultures of optical density between 3.2-3.6 were used to infect cells at an MOI of 1:10.

#### Preparation of cell lysate from Legionella infected cells

Legionella infected or non-infected HEK293T cells growing on a 10 cm dish were lysed in KHEM lysis buffer (20 mM Hepes-KOH (pH =7.5), 150 mM KCl, 2 mM MgCl_2_, 0.2 mM EDTA, 1% Triton-X100, protease inhibitor cocktail) 4 hours post infection. For lysis, cells were collected in PBS, centrifuged at 800 rcf to get a cell pellet. Lysis buffer was added to the cell pellet and incubated in ice for 10 min. This was then centrifuged at 15000g for 10 min. The pellet was discarded and the supernatant was used as the cell lysate for the pull-down assay.

#### Pull-down assay with biotin conjugated chemical probes

For each sample, 300 μl of lysate was incubated with 2 μM of one of the following conditions: 1) DMSO control, 0.4 μL. 2) Biotin-ubiquitin (Arg42 → Azido homoalanine) (0.3 μL of a 2 mM solution in DMSO, 2 μM final concentration) 3) Vinyl sulfonate probe **7** (0.3 μL of a 2 mM in DMSO, 2 μM final concentration). Lysates were incubated with the probes and controls at 37 °C for 2 hours. Streptavidin-agarose resin was equilibrated in lysis buffer. 30 μL of equilibrated resin was added to the lysate-probe mixture and the total volume of the reaction was adjusted to 1 mL by adding 700 μL of lysis buffer. This was then incubated overnight at 4 °C on a rotator. On the next day, the resin was washed with wash buffer (Tris 20 mM, NaCl 150 mM, pH 7.5, 2% SDS) by centrifuging at 500 rcf for 1 min followed by removal of the supernatant. The wash was repeated 6 times; the resin was then transferred to new microfuge tubes and boiled in SDS with β-mercaptoethanol for 15 min. Subsequently, the samples were run on SDS-PAGE followed by Coomassie Blue staining. This was then subjected to in-gel trypsin digestion and mass-spectrometry. The experiment was performed in biological triplicates for both non-infected and Legionella-infected HEK293T cells.

#### Bioinformatic analysis

Peptide intensity table with the Label free quantitation (LFQ) values were analyzed in Perseus (v1.6.2.3). Data were log2 transformed and filtered for identification in all three replicates in at least one group. Principal component analysis (PCA) was performed for each analysis with default settings. Intensities were *Z* scored by subtracting the mean and used for hierarchical clustering by Euclidean distance (pre-processed with k-means, 300 clusters, 1000 iterations) (**Supplementary Fig. 14**). Missing values were imputed from the lower end of the normal distribution (default settings). A two-sided student’s *t* test with permutation-based FDR was used to calculate significance between probe pulldown and control with/without infection, at 0.05 FDR (*p* value).

## Data availability

The mass spectrometry proteomics data has been deposited to the ProteomeXchange Consortium via the PRIDE partner repository with the dataset identifier *PXD049797*. The structure coordinates of the DupA:probe **5** complex has been deposited in the PDB with accession code *9EMK*. Full gel images and biochemical assay readings are provided in the Source Data file.

## Acknowledgements

Work in the van der Heden van Noort laboratory was supported by ZonMw (Off-road Grant np. 451001026), Dutch Research Council (NWO VIDI Grant, no. VI.Vidi.192.011) and the European Research Council (ERC CoG Grant, no. 101087582) to GJVDHVN. This work was further supported by the European Union’s Horizon 2020 research and innovation programme under the TRIM-NET project, Marie Skłodowska-Curie Action - Innovative TrainingNetworks (H2020-MSCA-ITN-2018) - GA 813599 to RM and a National Research Foundation of Korea (NRF) grant funded by the Korean government (MSIT and MOE) (Nos. 2021R1C1C1003961 and 2018R1A6A1A03025607) and the Yonsei University Research Fund of 2021 (2021-22-0050) to DS. ESRF (proposal mx2407) beamline scientists are gratefully acknowledged, especially those on beamline ID30A1, for their automated data collection service.

## Author contributions

The concept of this study was conceived by G.J.v.d.H.v.N. Synthesis of Ub probes was executed by M.S.K with technical assistance from C.M.P.T.O. Protein expression and purification was executed by A.M and R.Q.K. Biochemical assays were performed by M.S.K and supervised by A.S. X-ray crystallography was performed by R.Q.K and data processing by M.J., M.M., D.S. and R.Q.K. Cell culture was performed by R.M^b^ and M.M. in the lab of I.D. Proteomic measurements were performed by R.T.N.T supervised by P.A.v.V. Data analysis was performed by R.M.^a^ The manuscript was prepared by M.S.K and G.J.v.d.H.v.N. with input from all authors. The project was supervised by G.J.v.d.H.v.N.

