## Supplementary material for "Development of covalent probes to capture *Legionella pneumophila* effector enzymes": Sup. Info

**Chemical synthesis Probes 1-7** Experimental ............ 22-29

**HRMS data of probes 1-7**............ 30-36

**Riboside Synthesis** Experimental ............ 37-43

**References**............43

**NMR Spectra** ............ 43-58

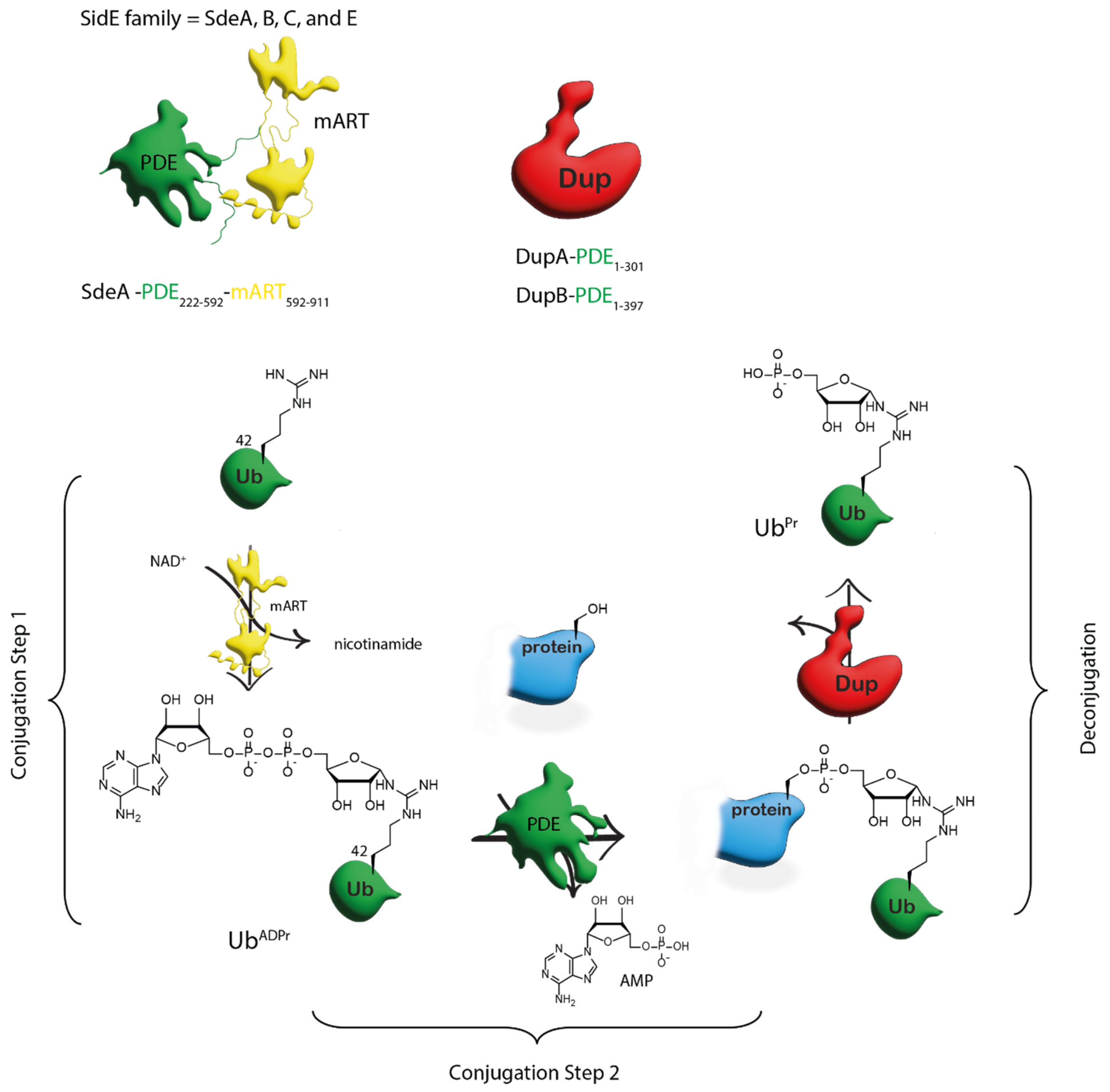

Supplementary Figure 1. Schematic representation of the *Legionella pneumophila* SidE-mediated PR-ubiquitination cascade.

Proposed mechanism

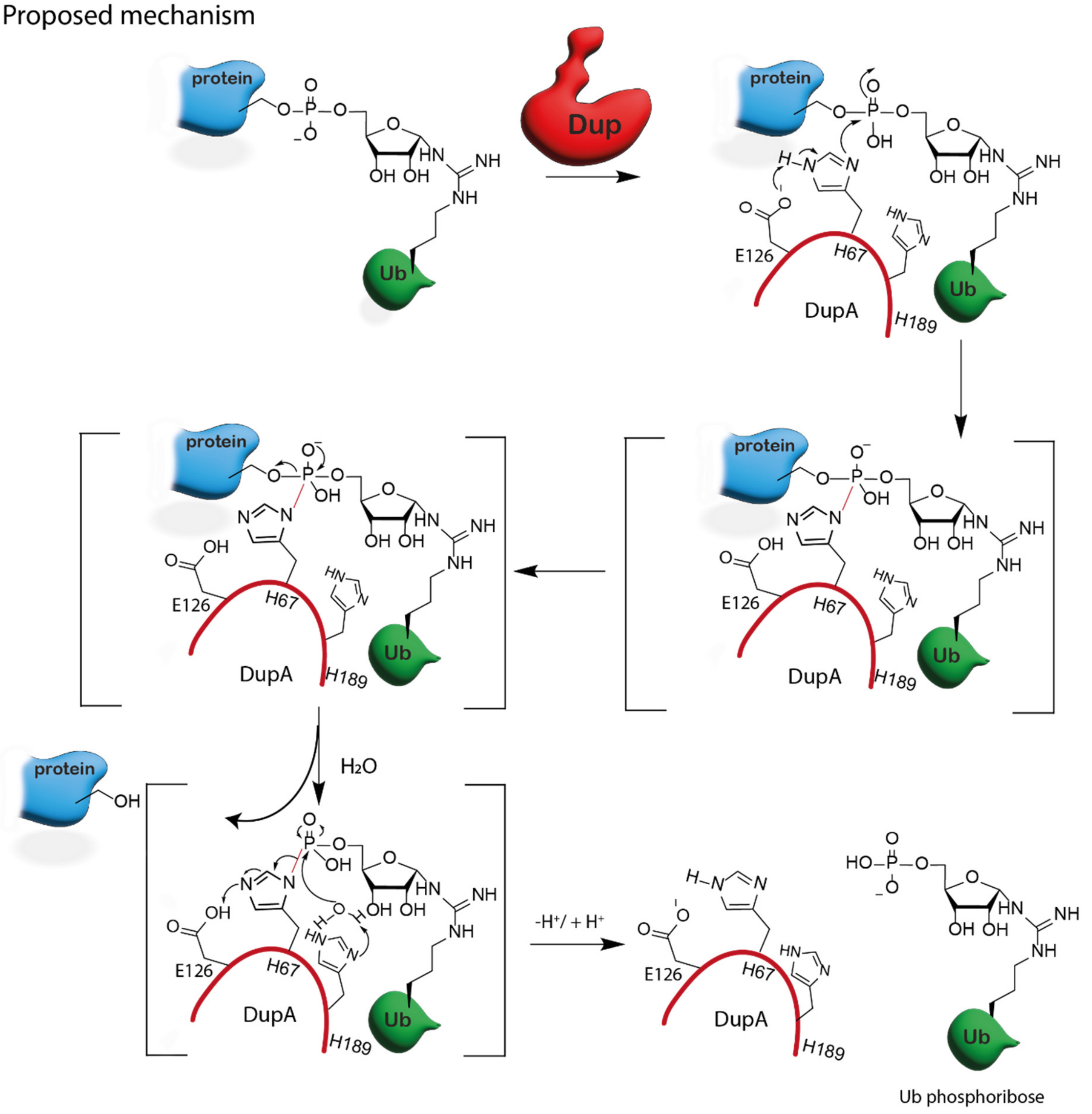

Supplementary Figure 2. The proposed catalytic mechanism involved in DupA mediated substrate-PR-ubiquitin hydrolysis.

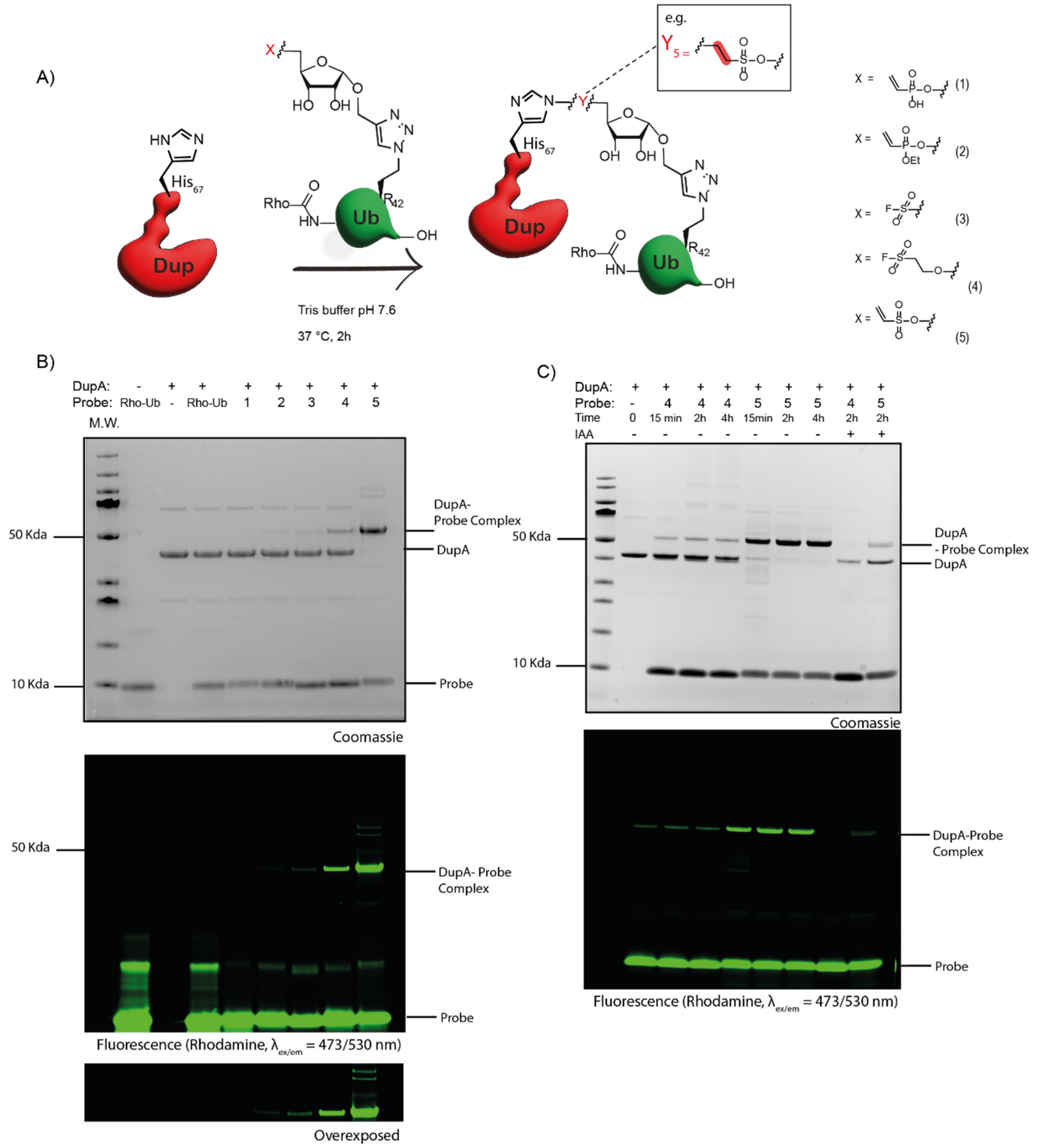

**Supplementary Figure 3.** Assessment of the reaction of probe library **1**-**5** on recombinant DupA WT. **A**) Schematic representation of the reaction between probes **1**-**5** and DupA WT, aiming to target His67 of the DupA active site, **B**) Probe **1-5** (8 equivalents) were incubated with DupA WT at 37 °C for 2 hours. SDS-PAGE gel: Coomassie staining (upper panel) and rhodamine fluorescence scan ($\lambda_{ex/em}$ = 473/530 nm) (bottom panel). **C**) Probe **4** and **5** (8 equivalents) were incubated with DupA wt at 37 °C and followed over time by quenching the samples with 3x sample buffer at the indicated time points. SDS-PAGE gel: Coomassie staining (upper panel) and rhodamine fluorescence scan ($\lambda_{ex/em}$ = 473/530 nm) (bottom panel). Iodo-acetamide (IAA) treatment was performed by preincubating DupA with IAA (10 mM, 30 min at 37 °C).

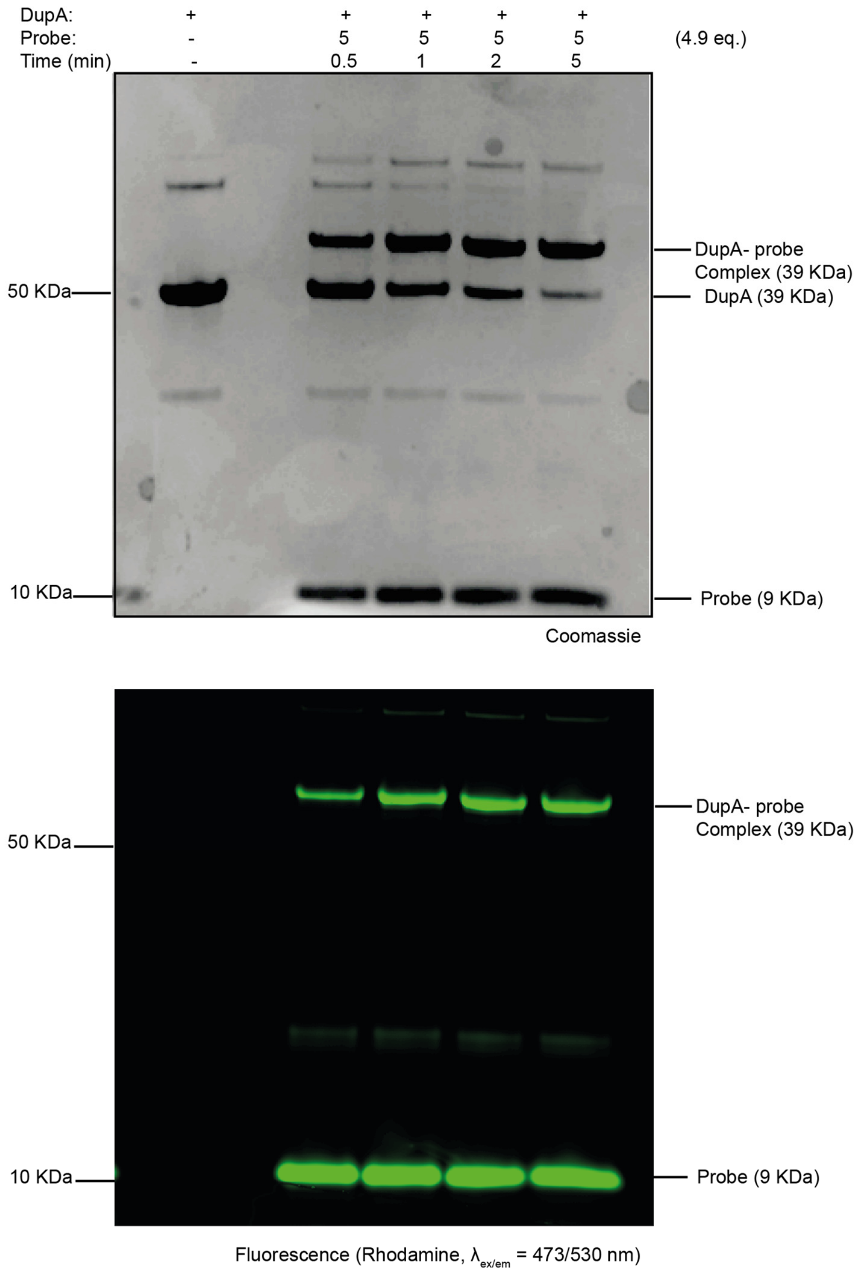

**Supplementary Figure 4.** SDS-PAGE analysis of the reaction between recombinant DupA WT and vinyl sulfonate probe **5** ( 5 equivalents) on ice for the indicated time points. SDS-PAGE gel: Coomassie staining (upper panel) and rhodamine fluorescence scan ($\lambda_{ex/em}$ = 473/530 nm) (bottom panel).

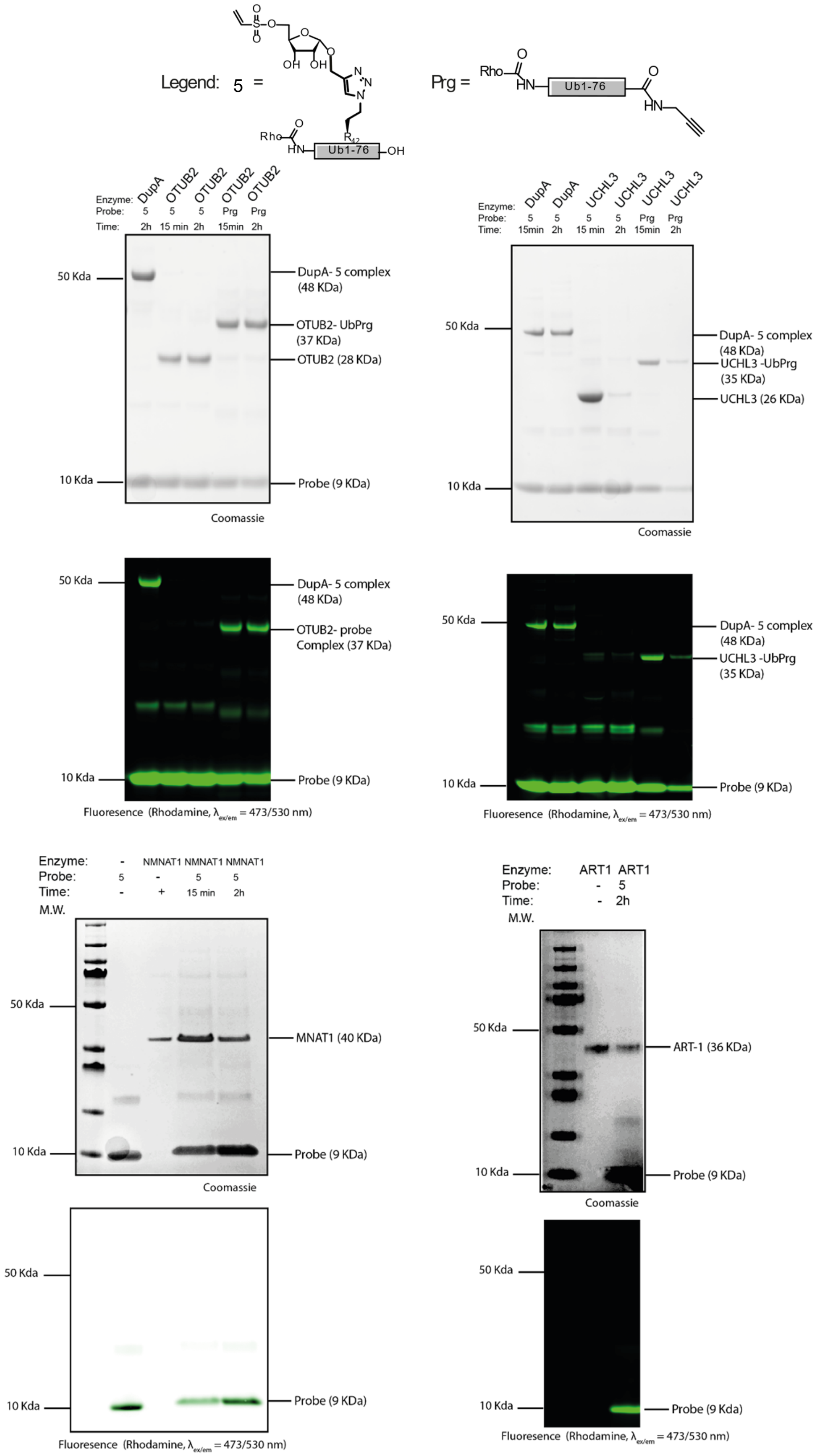

**Supplementary Figure 5.** Labeling assay of vinyl sulfonate probe **5** on recombinant proteins other then DupA. Probe **5** (8 equivalents) was incubated with one of the recombinant enzymes (OTUB2, UCHL3, NMNAT1 or ART-1) for the indicated time points at 37 °C. For the DUBs (OTUB2 and UCHL3) activity was confirmed with reactivity towards Rhodamine-Ub-Prg (8 eq.).

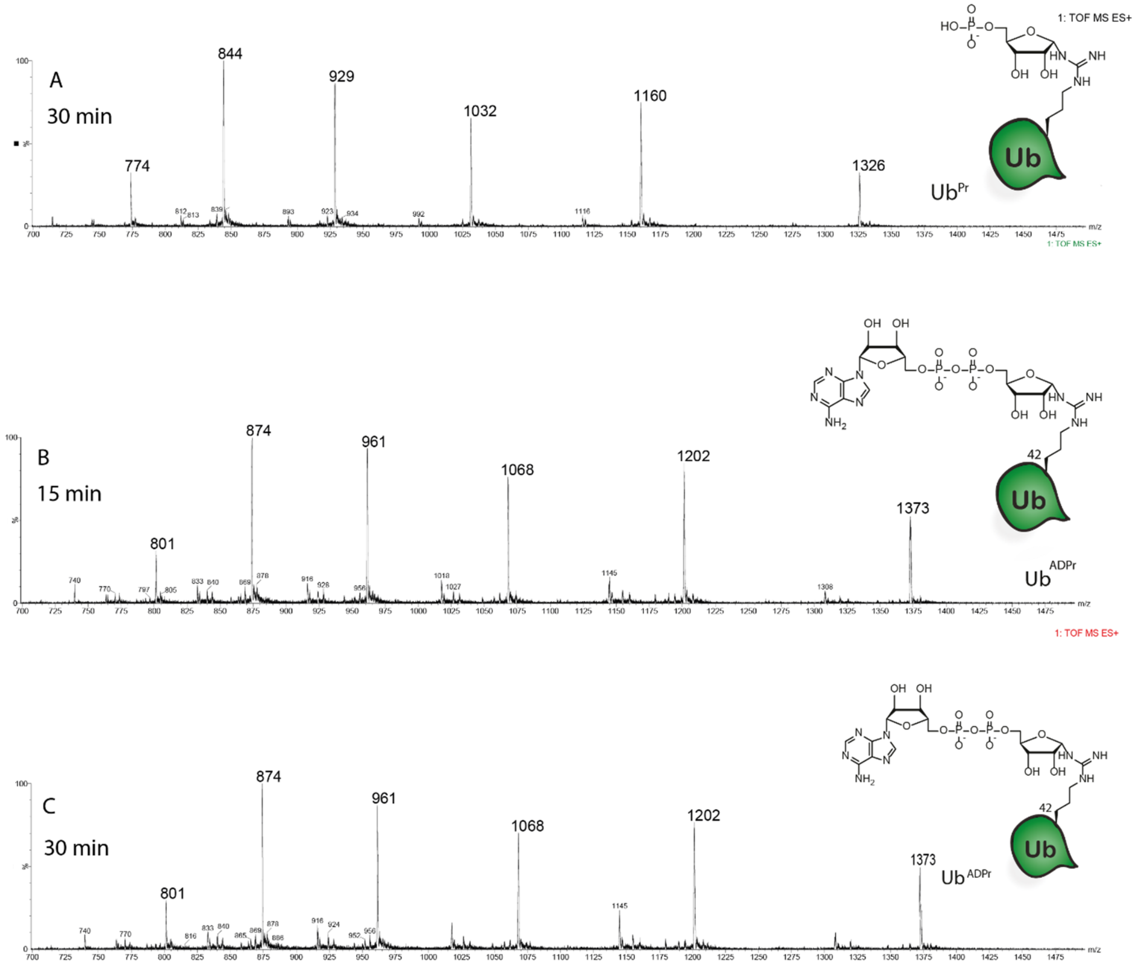

**Supplementary Figure 6.** Mass spectrometry analysis of the DupA:Probe **5** complex incubated with $Ub^{ADPr}$ as DupA substrate. The processing of $Ub^{ADPr}$ to $Ub^{PR}$ was analyzed. **A)** DupA was pre-incubated for 2 hours with Rhodamine-Ub (8 eq.) at 37 °C. Subsequently, $Ub^{ADPr}$ (7.3 eq.) was added and the mixture incubated for an additional 30 min at 37 °C before analyses by mass spectrometry, showing full consumption of $Ub^{ADPr}$ to $Ub^{PR}$. **B, C)** DupA was incubated with vinylsulfonate probe **5** (8 eq.) for 2 hours resulting in the formation of the DupA-vinylsulfonate-Ub complex. Subsequently, $Ub^{ADPr}$ (7.3 eq) was added and $Ub^{ADPr}$ to $Ub^{Pr}$ processing analyzed by mass spectrometry, showing no consumption of $Ub^{ADPr}$ **B)** 15 min and **C)** 30 min.

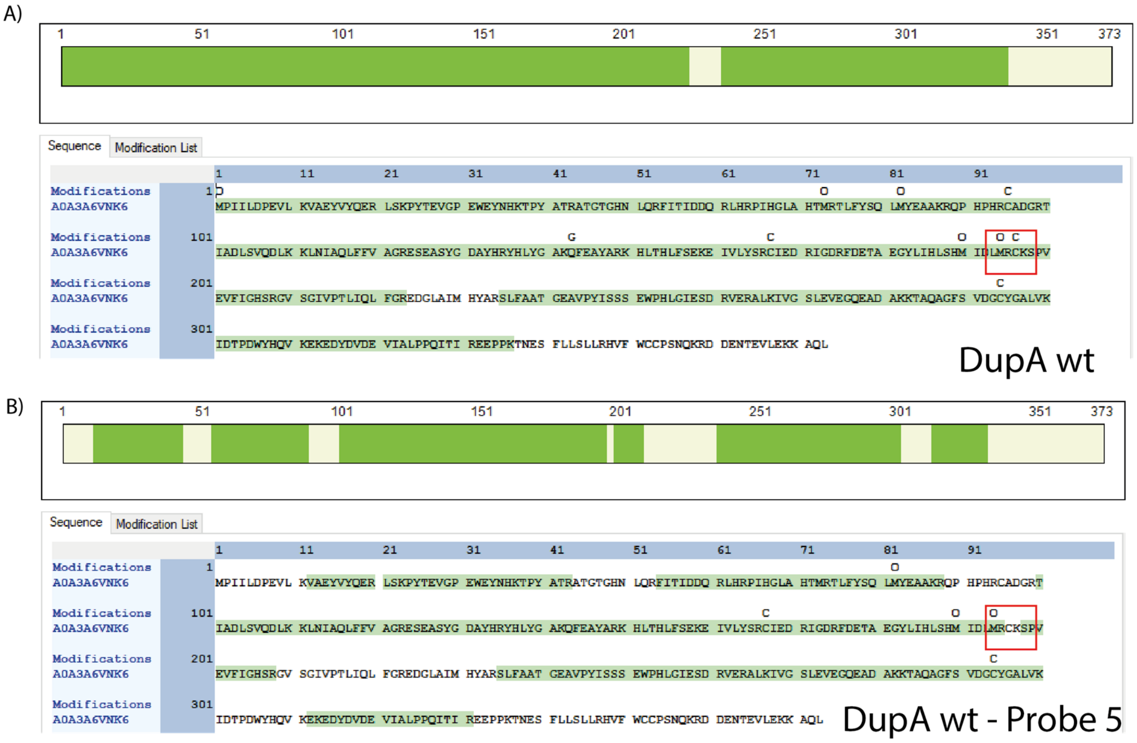

**Supplementary Figure 7**. Mapping of tryptic peptide sequence coverage by MS analysis of A) DupA WT and **B**) the DupA:probe **5** complex. The highlighted green sections in the sequence indicates the identified tryptic peptides. Red box indicates the difference in identification of tryptic peptide $C_{196}K_{197}$.

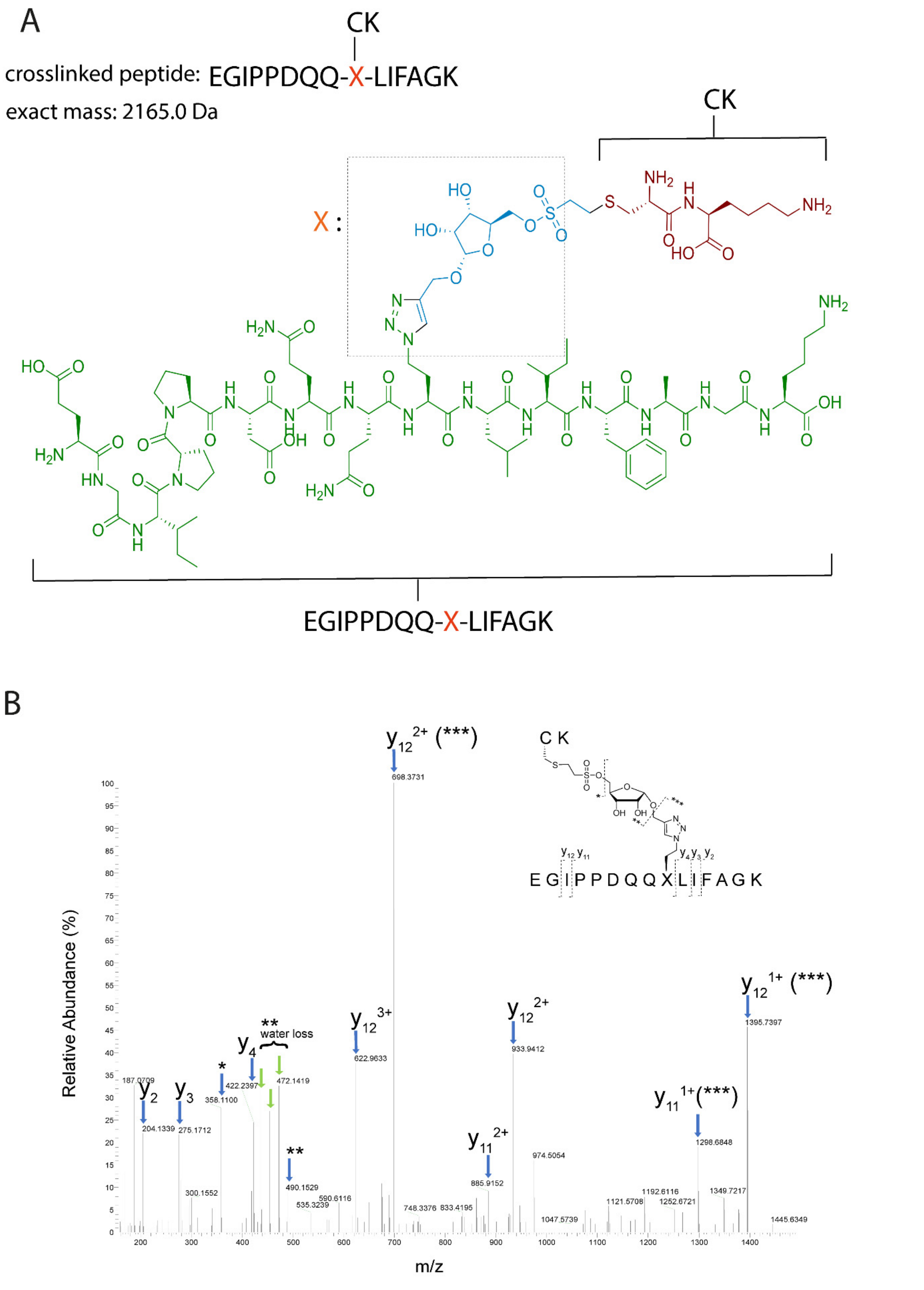

**Supplementary Figure 8**. **A)** Chemical structure of the DupA-probe **5** crosslinked tryptic peptide fragment. **B)** Annotated MS/MS spectrum identifying the crosslink between cysteine (DupA at position 196) and Ub probe **5** (at position 42).

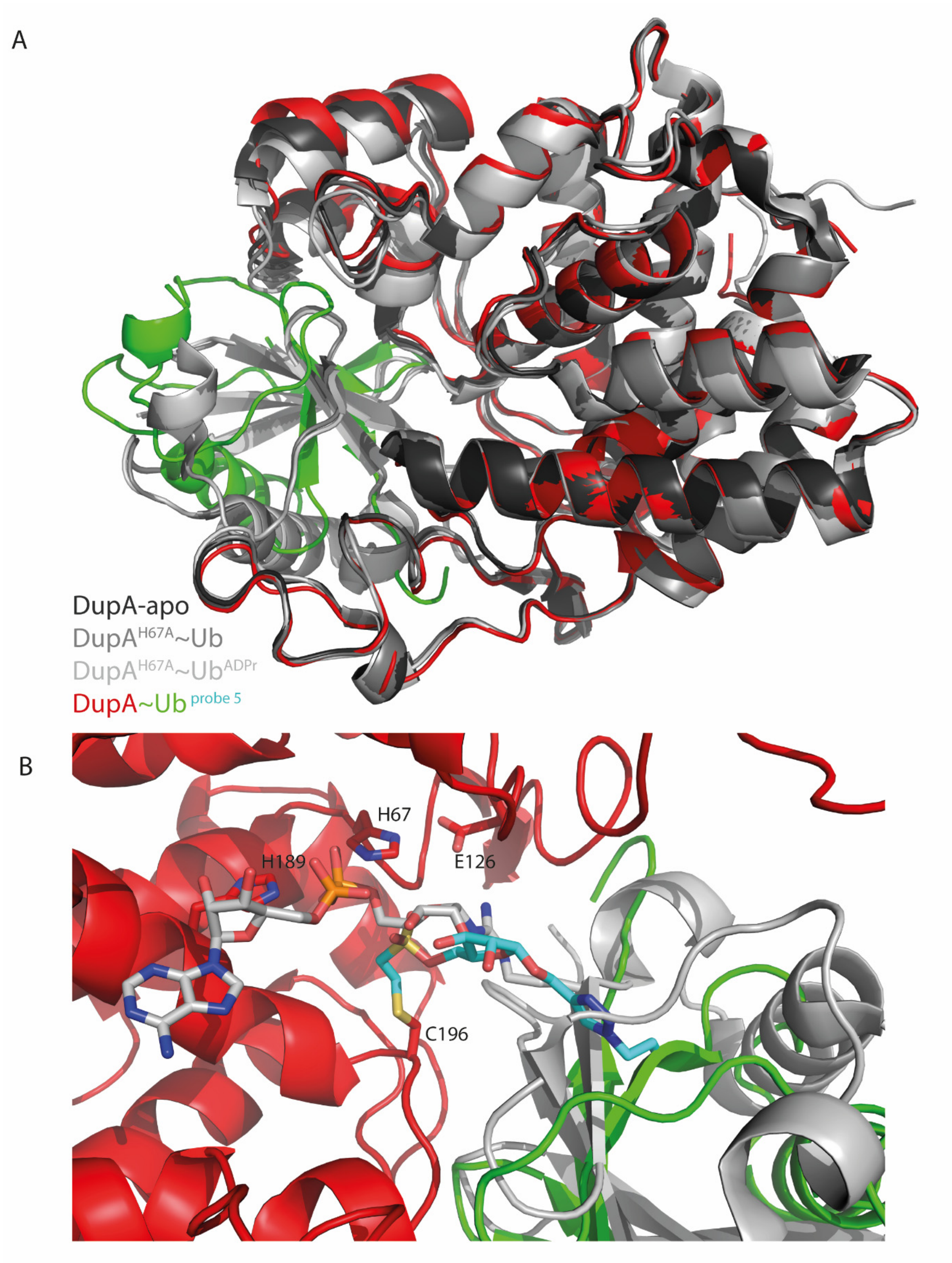

**Supplementary Figure 9**. **A**) Overlay of DupA$_{APO}$ (PDB: 6RYB) in dark grey, DupA$^{H67A}$:Ub (PDB:6RYA) in grey, DupA$^{H67A}$:Ub$^{ADPr}$ (PDB:6B7O) in light grey, DupA:Ub$^{Probe\ 5}$ (PDB: 9EMK) in red/green. **B**) Zoom in on active site; overlay of DupA$^{H67A}$:Ub$^{ADPr}$ (PDB:6B7O) in light grey, DupA:Ub$^{Probe\ 5}$ (PDB: 9EMK) in red/green.

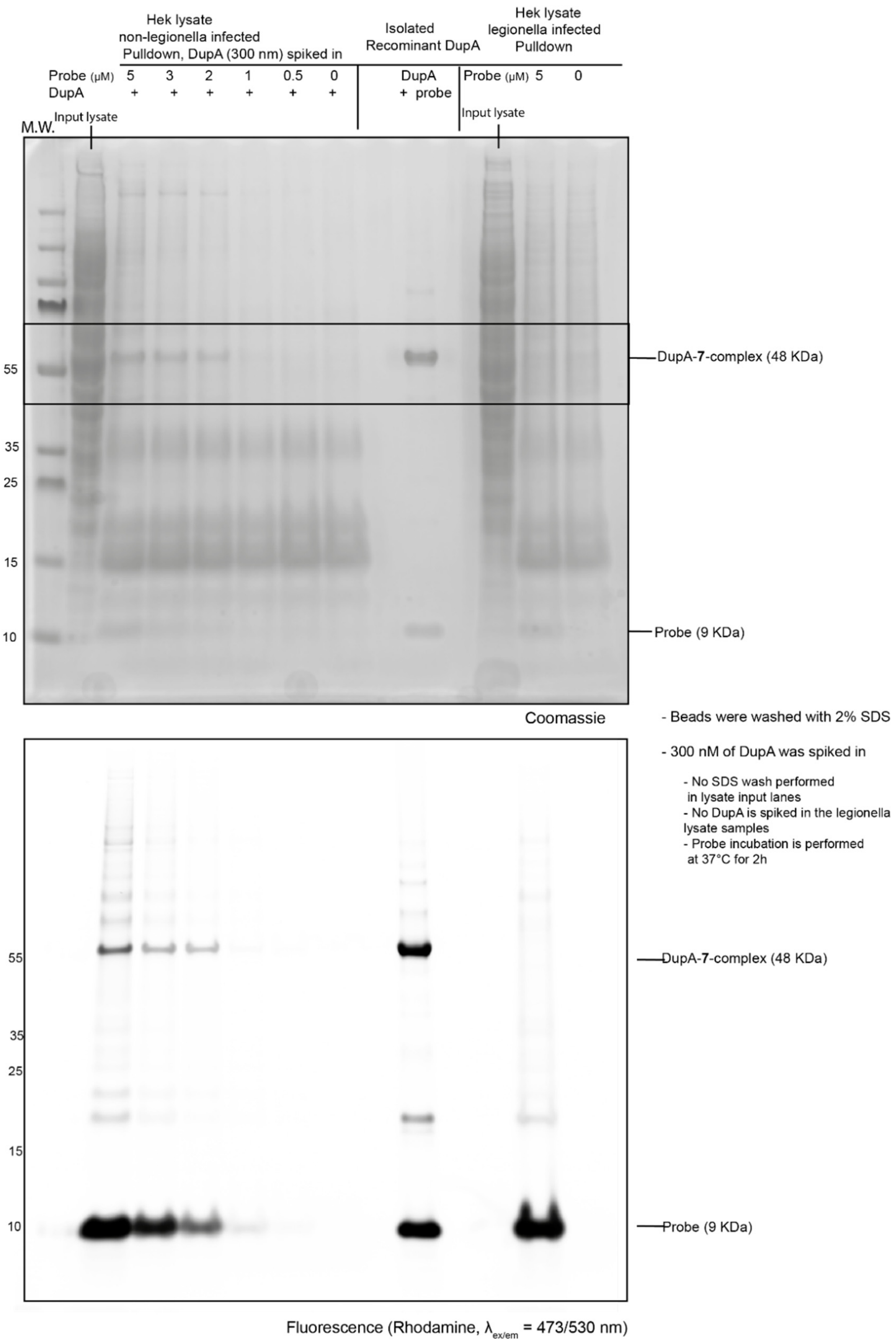

**Supplementary Figure 10**. DupA-Probe **7** complex recovery via biotin-streptavidin pulldown in HEK293T lysate. The pulldown was performed using different concentrations of probe **7** (0 – 5 µM) in non-infected HEK293T cells or HEK293T cells infected with *Legionella pneumophila*. For the non-infected HEK293T lysate, DupA (300 nM) was spiked in. The probe incubation was performed at 37 °C for 2 hours. After probe incubation the beads were washed with buffer (Tris 20 mM, NaCl 150 mM, pH 7.5, including 2% SDS). Upper panel: Coomassie stain, bottom panel: rhodamine fluorescence scan ($\lambda_{ex/em}$ = 473/530 nm).

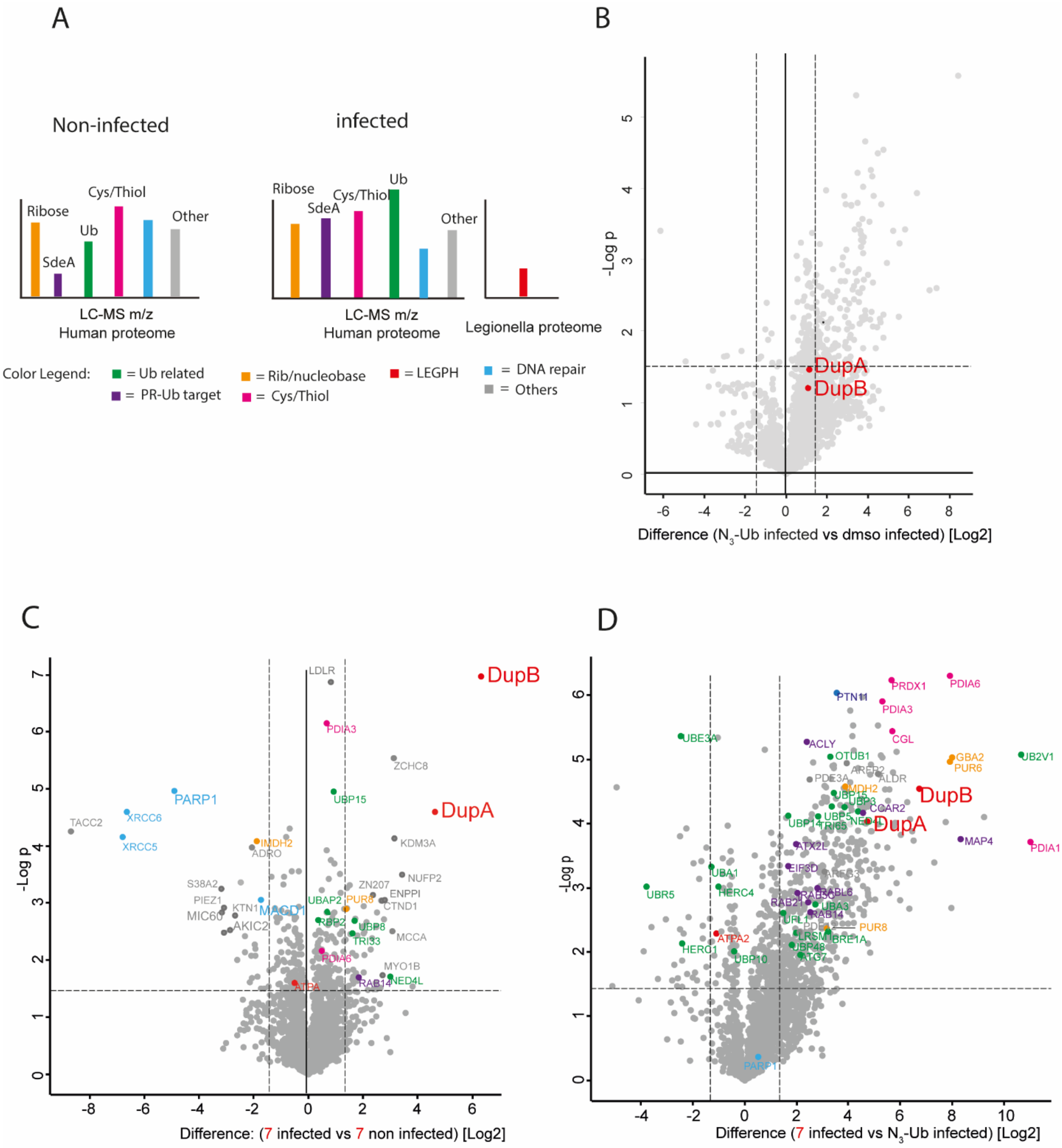

**Supplementary Figure 11**. Chemoproteomic assessment of the pulldown by the vinylsulfonate probe **7** in non-infected- or HEK293T cells infected with Legionella **A**) Color legend representing clusters of significantly enriched proteins **B**) volcano plot comparing biotin-Ub and DMSO in the legionella infected sample group, highlighting DupA and DupB being present in the lysate but not being significantly enriched using either of the controls. **C**) volcano plot highlighting the significant enrichment of proteins by probe **7** comparing the infected sample group to the non-infected group. **D**) volcano plot showing the significant enriched proteins with probe 7 in the infected sample group compared to biotin-Ub. For every plot holds: black dashed lines correspond to the thresholds: $\log_2$ ratio ≥1.5; p-value ≤0.05.

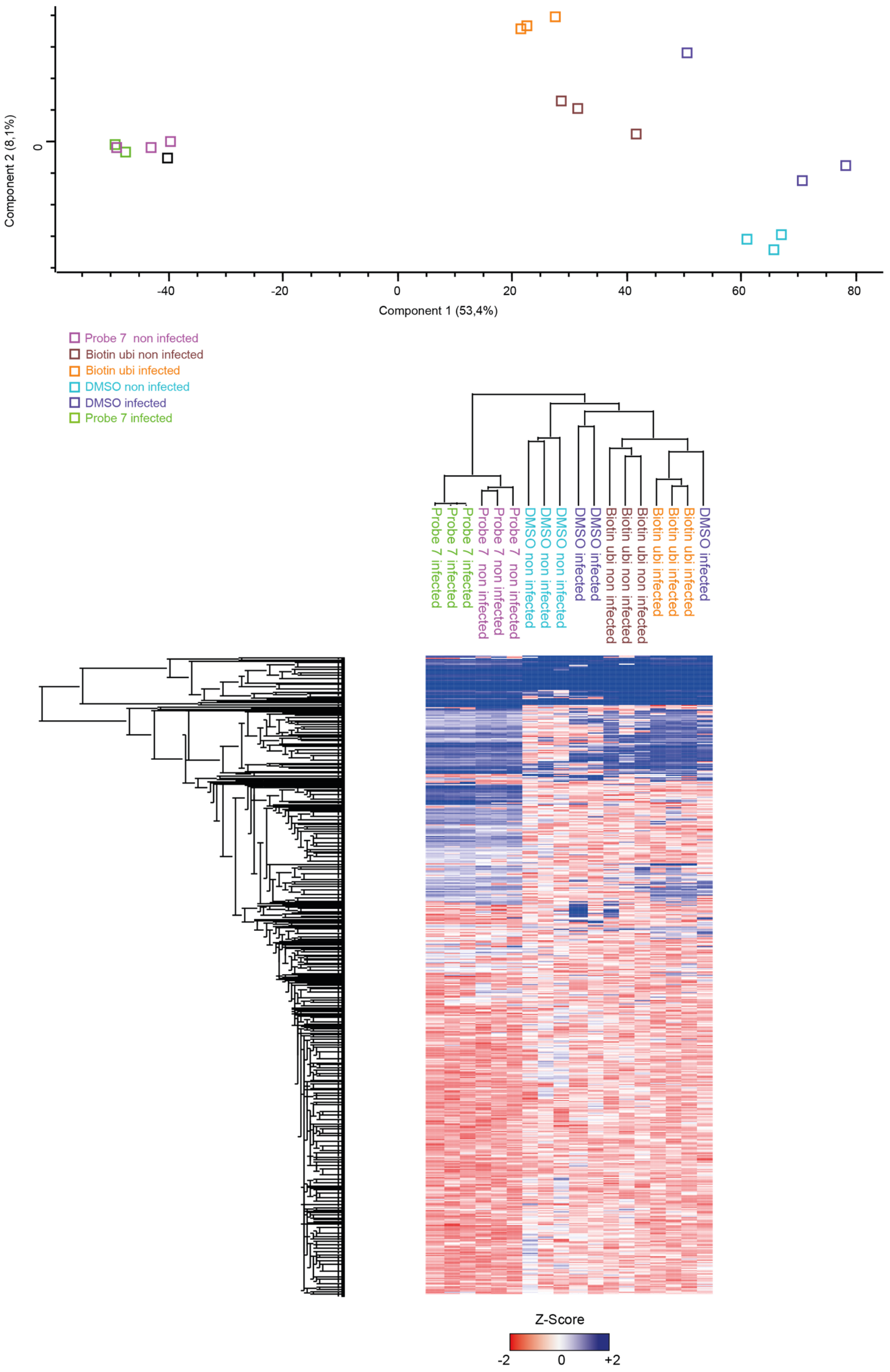

**Supplementary Figure 12.** PCA analysis and Hierarchical clustering. Figure legend for PCA analysis: Principal component analysis of the total proteome DIA data using components with the highest explained variance, n = 3. Figure legend for heat map: Heat map shows Hierarchical clustering of the significantly affected genes (p value < 0.05) from the proteome data set. Clustering done via Euclidean distance (pre-processed with k-means, 300 clusters, 1000 iterations), where each column in the heatmap is a sample and each row is a protein. The LFQ intensity values were normalized by Z-score.

**Supplementary Table 1.** Statistics for data processing and refinement of the co-crystal structure of DupA and compound **6**.

| | |
|---|---|
| Crystallization conditions | 20 %w/v PEG 6K, 0.1 M HEPES pH 7, 0.2 M $CaCl_2$ |
| PDB accession code | 9EMK |
| Space group | $P3_2$ |
| *Cell dimensions* | |
| a (Å) | 86.8 |
| b (Å) | 86.8 |
| c (Å) | 146.5 |
| α (°) | 90 |
| β (°) | 90 |
| γ (°) | 120 |
| *Processing statistics* | |
| Resolution (Å)<br>Outer shell (Å) | 75.18-2.17<br>2.22-2.17 |
| Beamline | ESRF ID30A-1 (MASSIF1) |
| Wavelength (Å) | 0.966Å |
| Observed reflections | 328821 (22513) |
| Unique reflections | 65360 (4611) |
| $R_{pim}$ | 0.043 (0.763) |
| CC(1/2) | 0.998 (0.388) |
| Multiplicity | 5.0 (4.9) |
| Completeness | 100 (100) |
| Mean (I/σ(I)) | 10.7 (0.9) |
| *Refinement statistics* | |
| Monomers in ASU | 6 |
| N° of protein atoms | 9221 (non-H) |
| Average B-factor | 66.0 |
| $R_{work}$ | 0.185 |
| $R_{free}$ | 0.239 |
| *RMSD from ideality* | |
| Bond lengths (Å) | 0.0083 |
| Bond angles (°) | 1.835 |
| *Values within parentheses are for the outer resolution shell* | |

### Biochemical procedures: using probe library 1-7 on recombinant protein, crystallization and in chemoproteomics.

#### Constructs and protein purifications

The bacterial expression construct for DupA has been described previously.[1] The mutations (H67A, H189A, the combination and C196A) were generated using overlapping primers[2] and all were sequence-verified before expression. The N-terminal GST-tagged DupA expression constructs were transformed into T7 Express cells and cultured at 37 °C in LB supplemented with 100 µg $mL^{-1}$ ampicillin. When $OD_{600}$ reached 0.6, expression was induced using 0.5 mM IPTG (Isopropyl β-D-1-thiogalactopyranoside) and temperature was lowered to 20 °C for 5 hours.

Harvested cell pellets were resuspended in GST buffer (50 mM Tris pH 7.5, 150 mM NaCl and 1 mM TCEP) and sonication on ice, before being spun down at 24 kG for 40 minutes at 4 °C. The resulting supernatant was applied to GST 4B Sepharose beads, followed by extensive washing with GST buffer and DupA elution using elution buffer (50 mM Tris pH 7.5, 50 mM NaCl, 25 mM GSH, 1 mM TCEP). Overnight incubation with TEV protease resulted in cleaved DupA and the protein solution was concentrated using spin filtration before a final purification step over a Superdex 75 16/60 column, equilibrated in a final buffer containing 20 mM HEPES pH 7.5, 100 mM NaCl and 1 mM DTT. All steps were analysed on SDS-PAGE and the final product was concentrated to ~10 mg $mL^{-1}$ before being flash frozen in liquid nitrogen.

Amino acids sequence DupA: (UNIPROT ID: A0A3A6VNK6)

```
GAMGS⁴ILDPEVLKVAEYVYQERLSKPYTEVGPEWEYNHKTPYATRATGTGHNLQRFITIDDQRLHRP
IHGLAHTMRTLFYSQLMYEAAKRQPHPHRCADGRTIADLSVQDLKKLNIAQLFFVAGRESEASYGDAY
HRYHLYGAKQFEAYARKHLTHLFSEKEIVLYSRCIEDRIGDRFDETAEGYLIHLSHMIDLMRCKSPVE
VFIGHSRGVSGIVPTLIQLFGREDGLAIMHYARSLFAATGEAVPYISSSEWPHLGIESDRVERALKIV
GSLEVEGQEADAKKTAQAGFSVDGCYGALVKIDTPDWYHQVKEKEDYDVDEVIALPPQITIREEPPKT
NESFLLSL³⁴⁵L
```

Calculated mass: 39.442 Da.
Observed mass: 39.442 Da.

#### Labeling assay on recombinant DupA; Reactivity of the probes (1-5)

The probes **1**-**5** (27.4 µM, 8.1 eq.) in buffer (20 mM TRIS, 150 mM NaCl, pH 7.6) were incubated with DupA (3.4 µM) at 37 °C in a total volume of 26.66 µL. After 15 min/ 2h (the indicated time points on SDS page **Supplementary Fig. 3**) of incubation 10 µL sample was taken and added to 5 µL loading buffer including β-mercaptoethanol. The samples were run on a NuPAGE™ 12% Bis-Tris gel in MES buffer, 190 mV, for 45 minutes. A fluorescence scan on a Typhoon FLA 9500 (rhodamine channel, $\lambda_{ex/em}$ = 473/530 nm) was performed to visualize the complex formed and additionally, the proteins were stained with Coomassie staining.

#### Labeling assay on recombinant DupA; time series on ice

Probe **5** (21.9 µM, 4.9 eq.) in buffer (20 mM TRIS, 150 mM NaCl, pH 7.6) was incubated with DupA wt (4.4 µM) which was kept on ice, in a total volume of 20 µL. The labelling reaction was performed at 0 °C and monitored at the indicated time points (0.5 min, 1 min, 2 min and

5 min). 5 µL of each sample was mixed with loading buffer including β-mercaptoethanol (10 µL) and subsequently ran on a NuPAGE™ 4-12% Bis-Tris gel in MES buffer, 190 mV for 45 minutes. A fluorescence scan on a Typhoon FLA 9500 (rhodamine channel, $\lambda_{ex/em}$ = 473/530 nm) was performed to visualize the complex formed and additionally, the proteins were stained with Coomassie staining.

**Labeling assay on recombinant DupA; generating the DupA-vinyl sulfonate probe complex, which upon full conversion is incubated with $Ub^{ADPr}$**

Probe **5** (27.4 µM, 8.1 eq.) in buffer (20 mM TRIS, 150 mM NaCl, pH 7.6) was incubated with DupA wt (3.4 µM) in a total volume of 26.66 µL. The reaction was incubated at 37°C and after 2h LC-MS indicated full conversion into the DupA-probe complex ($M + H^+$ = 48595). $Ub^{ADPr}$ (24.6 µM, 7.3 eq.) was added and the conversion of $Ub^{ADPr}$ to $Ub^{Pr}$ was monitored by LC-MS after 15 min and 30 min of incubation. The same reaction procedure was performed in parallel preincubating DupA wt (3.4 µM) with Rhodamine-Ub (27.4 µM, 8.1 eq.) instead of probe **5**, for 2 hours at 37°C. Subsequently, $Ub^{ADPr}$ (24.6 µM, 7.3 eq.) was added and the $Ub^{ADPr}$ to $Ub^{Pr}$ conversion monitored by LC-MS. The mass spectrometry analysis is depicted in **Supplementary Fig. 6.**

**Labeling assay of vinyl sulfonate probe 5 to recombinant OTUB2, UCHL3, NMNAT1 and ART-1**

Probe **5** (27.4 µM, 8.13 eq.) in buffer (20 mM TRIS, 150 mM NaCl, pH 7.6) was incubated with one the enzymes (OTUB2, UCHL3, NMNAT1 or ART-1, 3.4 µM) at 37 °C in a total volume of 26.66 µL for the indicated time points. For the DUBs (OTUB2 and UCHL3) reactivity towards Rhodamine-Ub-Prg (27.4 µM, 8.1 eq.) was taken along. After the incubation, 10 µL sample was taken and added to 5 µL loading buffer including β-mercaptoethanol. Samples were ran on a NuPAGE™ 4-12 % Bis-Tris gel in MES buffer, 190 mV for 45 minutes. A fluorescence scan on a Typhoon FLA 9500 (rhodamine channel, $\lambda_{ex/em}$ = 473/530 nm) was performed to visualize the complex formed and additionally, the proteins were stained with Coomassie staining. The SDS PAGE analysis is depicted in **Supplementary Fig. 5.**

**DupA-probe complexation for structural analysis**

Vinyl sulfonate probe **6** (77.2 µM, 1.4 eq.), which is the un-tagged probe variant, in buffer (20 mM TRIS, 150 mM NaCl, pH 7.6) was incubated with DupA wt (57.1 µM) in a total volume of 777.6 µL. The reaction was shaken at 37°C and monitored by LC-MS. After 30 min, additional probe **6** (48 µM, 37.5 µmol, 0.8 eq.) was added. Full conversion was reached after 1 hour ($M + H^+$ complex = 48.235 Da) and the complex was purified by gel filtration using a Superdex75 10/300 column equilibrated in 20 mM HEPES pH 7.5 and 100 mM NaCl.

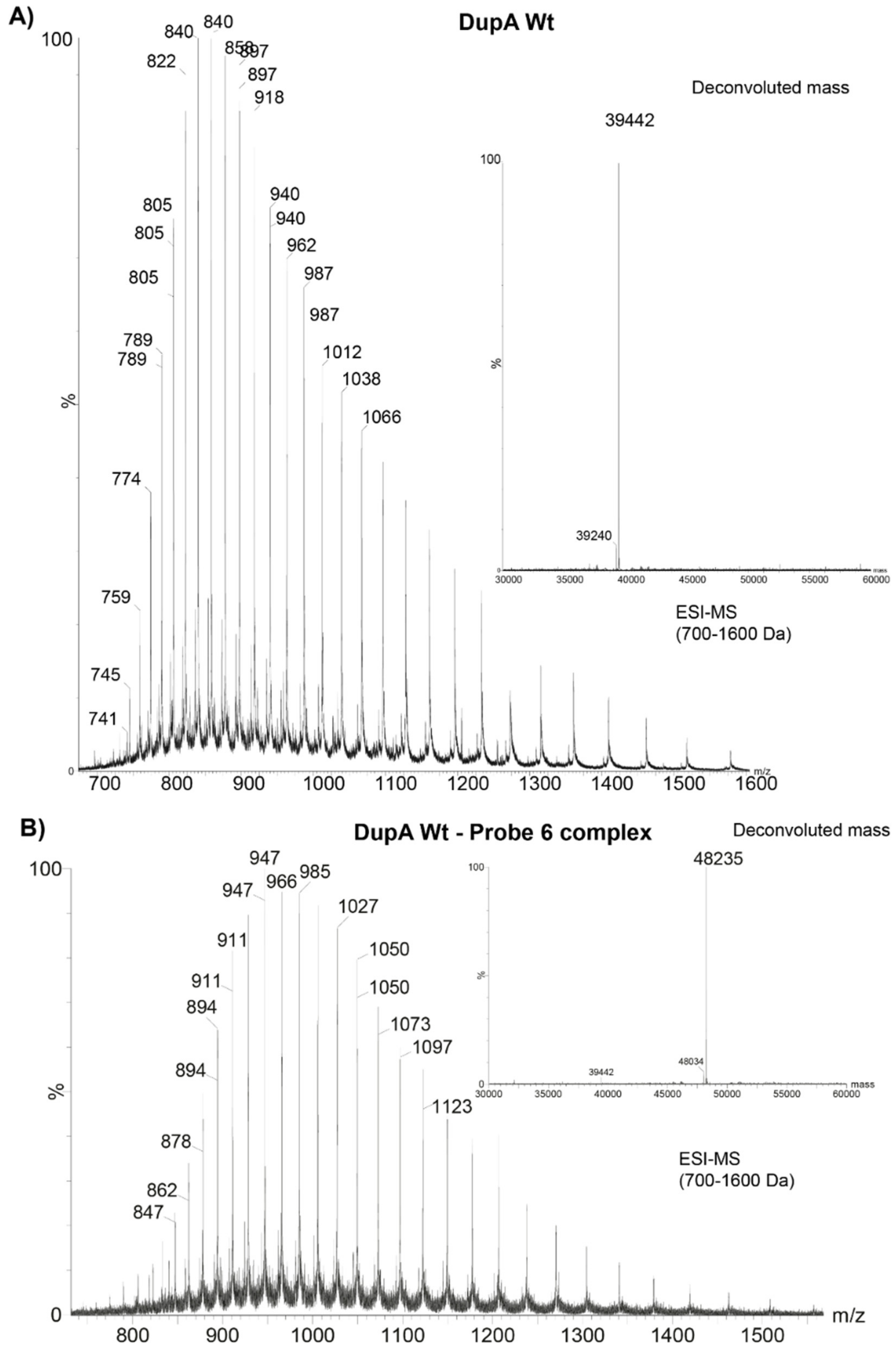

**Supplementary Figure 13.** HRMS spectra of **A**) DupA WT, prior to reaction with probe **6**. **B**) The DupA-:probe **5** complex formed after 2 hours incubation with probe **6**. ESI and deconvoluted masses (inset) are shown.

**Crystallization**

The complex of DupA and compound **6** was set up at a concentration of 7.2 mg $mL^{-1}$ in sitting drops using a 1:1 ratio. Crystals appeared within 3 days in a condition with 0.1M HEPES pH 7, 0.2 M $CaCl_2$ and 20 % (w/v) PEG-6000 and grew about 2 weeks. Single crystals were harvested in mother liquor supplemented with 30 % ethylene glycol (cryo-protectant) and shipped to ESRF for diffraction experiments at beamline ID30A1 (MASSIF1[3]). Diffraction data were processed using DIALS[4] before going through a CCP4[5] pipeline consisting of Aimless[6] data reduction, PHASER[7] molecular replacement using PDB 6RYA and iterative refinement and building cycles using COOT[8] and REFMAC.[9] The remaining resulting density could be built with probe **6** using restraints generated by the GRADE webserver (Global Phasing Ltd.), before final refinement optimization using PDB-REDO.[10] Statistics on both processing and refinement are aggregated in Supplemental Table 1 and final model is deposited at the Protein Data Bank.

**Pulldown in Legionella-infected HEK293T lysate and chemoproteomics**

**Cell lines**

HEK293T expressing CD32 were cultured in DMEM supplemented with 10 % FBS, 100 I.U./mL penicillin and 100 mg/mL streptomycin (Pen/Strep) at 37 °C, 5 % $CO_2$.

**Legionella pneumophila culture and infection**

Wild type L. pneumophila (Lp02) was grown for 3 days on N-(2-acetamido)-2-amino-ethanesulfonic acid (ACES)-buffered charcoal-yeast (BCYE) extract agar, at 37 °C, followed by growth for 20 h in ACES yeast extract media. Bacterial cultures of optical density between 3.2-3.6 were used to infect cells at an MOI of 1:10.

**Preparation of cell lysate from Legionella infected cells.**

Legionella infected or non-infected HEK293T cells growing on a 10 cm dish were lysed in KHEM lysis buffer (20 mM Hepes-KOH (pH =7.5), 150 mM KCl, 2 mM $MgCl_2$, 0.2 mM EDTA, 1% Triton-X100, protease inhibitor cocktail) 4 hours post infection. For lysis, cells were collected in PBS, centrifuged at 800 rcf to get a cell pellet. Lysis buffer was added to the cell pellet and incubated in ice for 10 min. This was then centrifuged at 15000g for 10 min. The pellet was discarded and the supernatant was used as the cell lysate for the pull-down assay.

**Pull-down assay with biotin conjugated chemical probes**

For each sample, 300 µl of lysate was incubated with 2 µM of one of the following conditions: 1) DMSO control, 0.36 µL. 2) Biotin-ubiquitin (R42 → Azido homoalanine) (0.3 µL of a 2 mM solution in DMSO, 2 µM final concentration) 3) Vinyl sulfonate probe **7** (0.3 µL, of a 2 mM in DMSO, 2 µM final concentration). Lysates were Incubated with the probes and controls at 37°C for 2 hours. Streptavidin-agarose resin was equilibrated in lysis buffer. 30 µL of equilibrated resin was added to the lysate-probe mixture and the total volume of the reaction was adjusted to 1 mL by adding 700 µL of lysis buffer. This was then incubated overnight at 4 °C on a rotator. On the next day, the resin was washed with wash buffer (Tris 20 mM, NaCl 150 mM, pH 7.5, 2 % SDS) by centrifuging at 500 rcf for 1min followed by removal of the supernatant. The wash was repeated 6 times; the resin was then transferred to new microfuge tubes and boiled in SDS with β-mercaptoethanol for 15 min. Subsequently, the samples were run on SDS-PAGE followed by Coomassie Blue staining. This was then subjected to in-gel trypsin digestion and mass-spectrometry.

**Proteomics mass spectrometry measurements**

Samples were loaded onto a 4-12 % Bis-Tris gradient gel (Invitrogen) and run for 1.5 cm. Subsequently, each lane was cut into four bands. Gel slices were first washed 3x with water, and subsequently subjected to reduction with 10 mM dithiothreitol, alkylation with 50 mM of iodoacetamide, and in-gel trypsin digestion using a Proteineer DP digestion robot (Bruker). After addition of trypsin (at 12.5 ng/µL) and swelling of the bands, the bands were transferred to Eppendorf vials and the bands were covered in 25 mM $NH_4HCO_3$ pH 8.3. Tryptic digestion took place overnight at 37 C and the peptides were extracted from the gel slices with with 50/50/0.1 v/v/v water/acetonitril/formic acid. Finally peptides were lyophilized. The experiment was performed in biological triplicates for both non-infected and Legionella infected HEK293T cells.

Tryptic peptides were dissolved in water/formic acid (100/0.1 v/v) and subsequently analyzed by on-line C18 nanoHPLC MS/MS with a system consisting of an Ultimate3000nano gradient HPLC system (Thermo, Bremen, Germany), and an Exploris480 mass spectrometer (Thermo). Fractions were injected onto a cartridge precolumn (300 µm × 5 mm, C18 PepMap, 5 um, 100 A, and eluted via a homemade analytical nano-HPLC column (50 cm × 75 µm; Reprosil-Pur C18-AQ 1.9 um, 120 A (Dr. Maisch, Ammerbuch, Germany). The gradient was run from 2% to 40 % solvent B (20/80/0.1 water/acetonitrile/formic acid (FA) v/v) in 30 min. The nano-HPLC column was drawn to a tip of ∼10 µm and acted as the electrospray needle of the MS source. The mass spectrometer was operated in data-dependent MS/MS mode for a cycle time of 3 seconds, with a HCD collision energy at 30 V and recording of the MS2 spectrum in the orbitrap, with a quadrupole isolation width of 1.2 Da. In the master scan (MS1) the resolution was 120,000, the scan range 400-1500, at standard AGC target @maximum fill time of 50 ms. A lock mass correction on the background ion m/z = 445.12 Da was used. Precursors were dynamically excluded after n=1 with an exclusion duration of 10 s, and with a precursor range of 20 ppm. Charge states 2-5 were included. For MS2 the first mass was set to 110 Da, and the MS2 scan resolution was 30,000 at an AGC target of 100% at maximum fill time of 60 ms. In a post-analysis process, raw data were first converted to peak lists using Proteome Discoverer version 2.2 (Thermo Electron), and submitted to the combined Uniprot database (Homo sapiens, 20596 entries, and the Legionella pneumophila subsp. pneumophila (strain Philadelphia 1 / ATCC 33152 / DSM 7513), UP000000609, 2930 entries), using Mascot v. 2.2.07 (www.matrixscience.com) for protein identification. Mascot searches were with 10 ppm and 0.02 Da deviation for precursor and fragment mass, respectively, and trypsin as enzyme. Up to two missed cleavages were allowed. Methionine oxidation and acetyl on protein N-terminus were set as a variable modification; carbamidomethyl on Cys, were set as a fixed modification. Protein FDR was set to 1 %. Normalization was on total peptide amount.

**Bioinformatic analysis:**

Peptide intensity table with the Label free quantitation (LFQ) values were analyzed in Perseus (v1.6.2.3). Data were log2 transformed and filtered for identification in all three replicates in at least one group. Principal component analysis (PCA) was performed for each analysis with default settings. Intensities were *Z* scored by subtracting the mean and used for hierarchical clustering by Euclidean distance (pre-processed with k-means, 300 clusters, 1000 iterations) (**Supplementary Fig. 13**). Missing values were imputed from the lower end of the normal distribution (default settings). A two-sided student's *t* test with permutation-based FDR was used to calculate significance between probe pulldown and control with/without infection, at 0.05 FDR (*p* value).

### Chemical synthesis

#### General synthetic procedures

All reagents were used as received unless stated otherwise. Solvents used in synthesis were dried and stored over 4Å molecular sieves, except for MeOH and MeCN which were stored over 3Å molecular sieves. Triethylamine (TEA) and diisopropylethylamine (DIPEA) were stored over KOH pellets. Column chromatography was performed on silica gel 60 Å (40-63 µm, Macherey-Nagel). TLC analysis was performed on Macherey-Nagel aluminium sheets (silica gel 60 $F_{254}$). TLC was used to visualize compounds by UV at wavelength 254 nm and by spraying with either cerium molybdate spray (25 g/L $(NH_4)_6Mo_7O_{24}$, 10 g/L $(NH_4)_4Ce(SO_4)_4 \cdot H_2O$ in 10% $H_2SO_4$ water solution) or $KMnO_4$ spray (20 g/L $KMnO_4$ and 10 g/L $K_2CO_3$ in water) followed by charring at c.a. 250 $^oC$. NMR spectra were recorded on a Bruker AV-300 NMR. Chemical shifts (δ) are given in ppm relative to tetramethyl silane. Coupling constants (*J*) are given in Hz. All given $^{13}C$-APT spectra are proton decoupled.

#### LC-MS measurements and HPLC purifications

LC-MS measurements were conducted on a Waters ACQUITY UPLC H-class System equipped with a Waters ACQUITY Quaternary Solvent Manager (QSM), Waters ACQUITY UPLC Photodiode Array (PDA) eλ Detector (λ = 210-800 nm), Waters ACQUITY UPLC Protein BEH C18 column (1.7 µM, 2.1 x 50 mm) and LCT Premier Orthogonal acceleration Time of Flight Mass Spectrometer (m/z = 100-1600) in ES+ mode. Samples were run for 3min at 40 $^oC$ using 2 mobile phases: A: MQ + 0.1% formic acid, B: MeCN + 0.1% formic acid. Gradient: 0 - 95% B at a flow rate of 0.5 mL/min. Data processing was performed using Waters MassLynx Mass Spectrometry Software 4.1 (deconvolution with MaxEnt1 function).

HPLC purification was performed on a **A)** Shimadzu semi-preparative RP-HPLC system, equipped with a Waters C18-Xbridge 5 µm OBD (10 x 150 mm) column at a flowrate of 6.5 mL/min. using 2 mobile phases: A: MQ + 0.05 % FA, B: MeCN + 0.05 % FA. Gradient: 10 -> 70 % B. HPLC system **B**) Waters preparative RP-HPLC system, equipped with a Waters C18-Xbridge 5 µm OBD (30 x 150 mm) column at a flowrate of 37.5 mL/min using 3 mobile phases: A: MQ, B: $CH_3CN$ and C: 1% TFA in MQ. Gradient: 20 -> 45 % B, 5 % C. High resolution mass spectra were recorded on a Waters XEVO-G2 XS Q-TOF mass spectrometer equipped with an electrospray ion source in positive mode (source voltage 3.0 kV, desolvation gas flow 900 L/hr, temperature 250 °C) with resolution R = 22000 (mass range m/z = 50-2000) and 200 pg/µL Leu-Enk (m/z = 556.2771) as a “lock mass”.

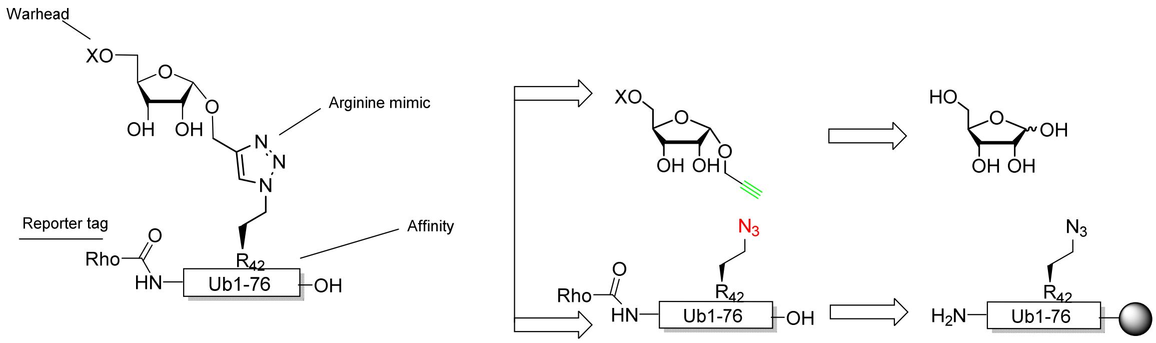

**Supplementary Scheme 1**. Retrosynthesis of warhead equipped Ub-phosphoribose mimics **1**-**5** starting from D-ribose and $Ub_{1\text{-}76}$ Arg42 → azido homoalanine on wang resin.

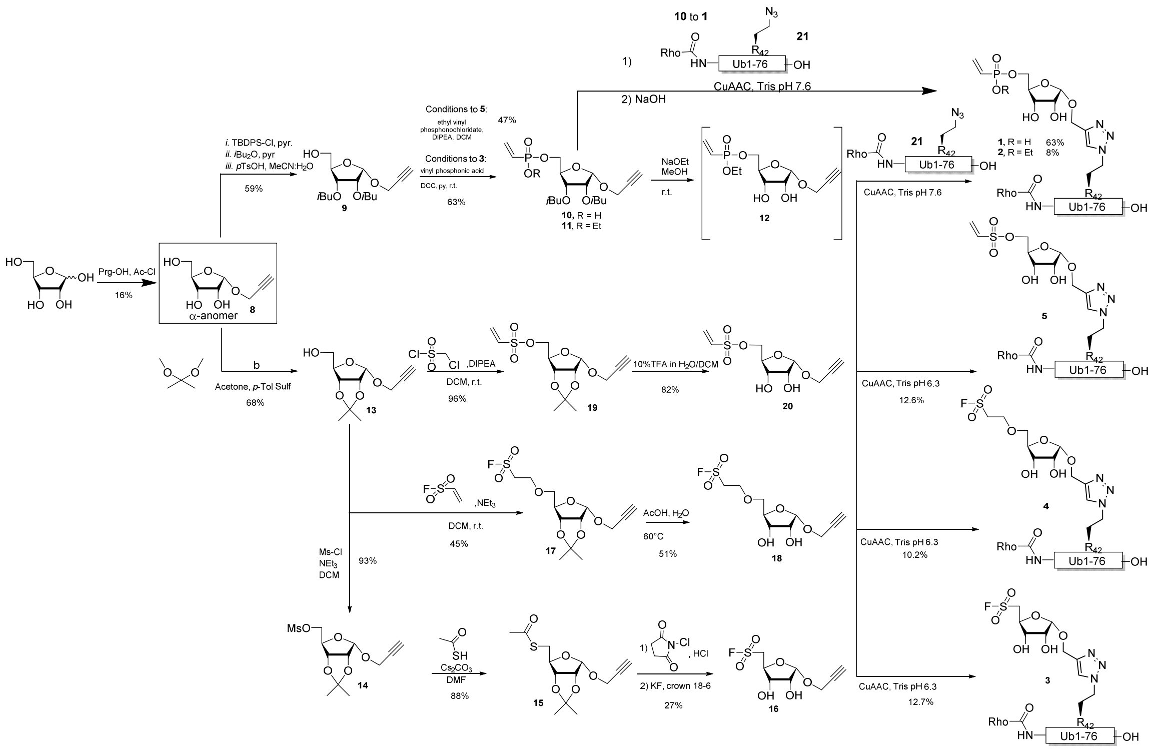

**Supplementary Scheme 2**. Synthesis of ribosides equipped with warheads and final conjugation to Ub via CuAAC.

### Chemical synthesis Probes 1-7 – Experimental

Rhodamine-(Arg42 → triazole linked vinyl-phosphonate ribose) $Ub_{76}$ (**1**)

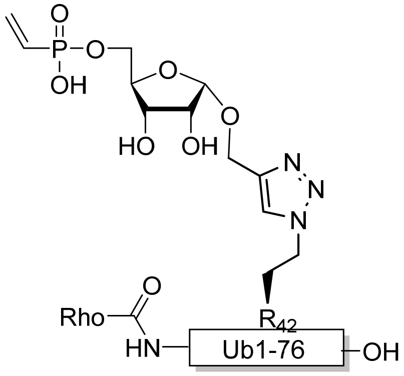

Protected vinyl phosphate **10** (7.3 µL, 0.15 M in DMSO, 1.1 µmol, 3.3 eq.) was added to Arg42 → azido homoalanine modified rhodamine ubiquitin **21** (M + $H^+$ = 9018 Da, 3 mg, 0.33 µmol, 1 eq.) dissolved in DMSO (50 µL) and TRIS buffer (150 µl, 20 mM TRIS, 150 mM NaCl, pH 7.6). To this solution was added 15 µL of freshly prepared click-mixture (1:1:1 v/v/v, $CuSO_4$ (100 mM in $H_2O$): Sodium Ascorbate (600 mM in $H_2O$): TBTA ligand (100 mM in MeCN). The reaction mixture was shaken at 37 °C for 1 hour after which LC-MS verified complete conversion to the clicked product (mass found: (M+$H^+$ = 9437 Da). EDTA (20 µL, 0.5 M in $H_2O$) was added quenching the reaction. Treatment with NaOH (150 µL, 0.5 M) resulted in complete deprotection of the secondary alcohols after shaking for 30 min, as verified by LC-MS (mass found: M + $H^+$ = 9296 Da). Subsequently, AcOH (100 µL) was added and the conjugate was purified by RP-HPLC. Pure fractions were pooled and lyophilized obtaining ubiquitin-ribose conjugate **1** (1.92 mg, 0.21 µmol, 63 %) as a red powder. HRMS: $[C_{414}H_{663}N_{108}O_{132}P + 7H]^{7+}$ found: 1329.2037, calculated: 1329.0657, $[C_{414}H_{663}N_{108}O_{132}P + 8H]^{8+}$ found: 1163.0448, calculated: 1163.0575, $[C_{414}H_{663}N_{108}O_{132}P + 9H]^{9+}$ found: 1033.9700, calculated: 1033.9400, $[C_{414}H_{663}N_{108}O_{132}P + 10H]^{10+}$ found: 930.7431, calculated: 930.6460, $[C_{414}H_{663}N_{108}O_{132}P + 11H]^{11+}$ found: 846.1327, calculated: 846.1614.

Rhodamine-(Arg42 → triazole linked vinyl-ethoxy-phosphonate ribose) $Ub_{76}$ (**2**)

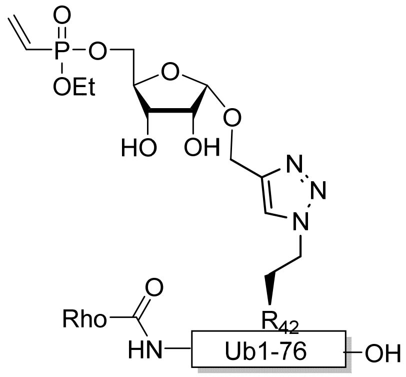

Etoxy vinyl phosphate **12** (5 µL, 0.15 M in DMSO, 0.75 µmol, 3.3 eq.) was added to Arg42 → azido homoalanine modified rhodamine ubiquitin **21** (M + $H^+$ = 8873 Da, 2 mg, 0.22 µmol, 1 eq.) dissolved in DMSO (33.3 µL) and TRIS buffer (100 µl, 20 mM TRIS/150 mM NaCl, pH 7.6). To this solution was added 10 µL of freshly prepared click-mixture (1:1:1 v/v/v, $CuSO_4$ (100 mM in $H_2O$): Sodium Ascorbate (600 mM in $H_2O$): TBTA ligand (100 mM in MeCN). The reaction mixture was shaken at 37 °C for 2 hours before the addition of another portion of freshly prepared click mixture (10 µL). After shaking for another hour LC-MS verified complete conversion to the clicked product (mass found: M+$H^+$ = 9180 Da). The conjugate was purified by RP-HPLC and the pure fractions were pooled and lyophilized obtaining ubiquitin-ribose conjugate **2** (0.17 mg, 0.018 µmol, 8.2 %) as a red powder. HRMS: $[C_{410}H_{656}N_{107}O_{129}P + 7H]^{7+}$ found: 1312.3680, calculated: 1312.3357. $[C_{410}H_{656}N_{107}O_{129}P + 8H]^{8+}$ found: 1148.5343, calculated: 1148.5343. $[C_{410}H_{656}N_{107}O_{129}P + 9H]^{9+}$ found: 1020.9572, calculated: 1020.9277. $[C_{410}H_{656}N_{107}O_{129}P + 10H]^{10+}$ found: 918.9562, calculated: 918.9350. $[C_{410}H_{656}N_{107}O_{129}P + 11H]^{11+}$ found: 835.4969 calculated: 835.4863.

Rhodamine-(Arg42 → triazole linked fluorosulfonate ribose) $Ub_{76}$ (**3**)

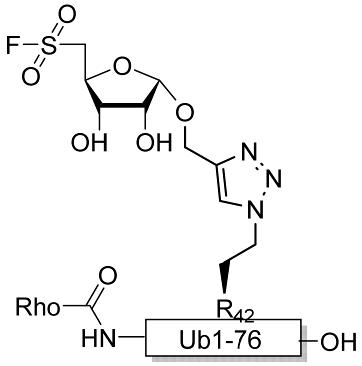

Fluoro-sulfonate **16** (5 µL, 0.15M in DMSO, 0.75 µmol, 3.3 eq.) was added to R42 → azido homoalanine modified rhodamine ubiquitin **21** (M + $H^+$ = 8874 Da, 2 mg, 0.22 µmol, 1 eq.) dissolved in DMSO (33.3 µL) and TRIS buffer (100 µL, 20 mM TRIS/150 mM NaCl, pH 6.3). The addition of 10 µL freshly prepared click mix (1:1:1 v/v/v, $CuSO_4$ (100 mM in $H_2O$): Sodium Ascorbate (600 mM in $H_2O$): TBTA ligand (100 mM in MeCN) started the reaction, which was subsequently shaken at 37 °C for 2 hours. LC-MS verified complete conversion to the clicked product (mass found: M+$H^+$ = 9127 Da) and the conjugate was purified by RP-HPLC. The pure fractions were pooled and lyophilized obtaining ubiquitin-ribose conjugate **3** (0.26 mg, 0.028 µmol, 12.7 %) as a red powder. HRMS: $[C_{406}H_{648}FN_{107}O_{128}S + 7H]^{7+}$ found: 1304.8702, calculated: 1304.9043. $[C_{406}H_{648}FN_{107}O_{128}S + 8H]^{8+}$ found: 1141.8917 , calculated: 1141.9163. $[C_{406}H_{648}FN_{107}O_{128}S + 9H]^{9+}$ found: 1015.1213, calculated: 1015.1477. $[C_{406}H_{648}FN_{107}O_{128}S + 10H]^{10+}$ found: 913.7167, calculated: 913.733. $[C_{406}H_{648}FN_{107}O_{128}S + 11H]^{11+}$ found: 830.7404, calculated: 830.7573.

Rhodamine-(Arg42 → triazole linked fluorosulfonate ribose) $Ub_{76}$ (**4**)

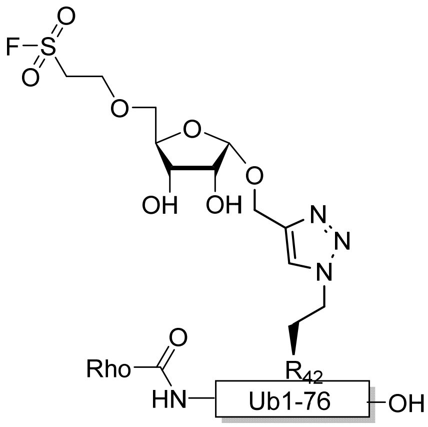

Fluorosulfonate **18** (10 µL, 0.15 M in DMSO, 1.5 µmol, 6.6 eq.) was added to R42 → azido homoalanine modified Rhodamine ubiquitin **21** (M + $H^+$ = 8874 Da, 2 mg, 0.22 µmol, 1 eq.) dissolved in DMSO (33.3 µL) and TRIS buffer (100 µL, 20 mM TRIS/150 mM NaCl, pH 6.3). 15 µL freshly prepared click mix (1:1:1 v/v/v, $CuSO_4$ (100 mM in $H_2O$): Sodium Ascorbate (600 mM in $H_2O$): TBTA ligand (100 mM in MeCN) was added and reaction was shaken at 37 °C for 1.5 hours. LC-MS verified complete conversion to the clicked product (mass found: M+$H^+$ = 9171 Da). The conjugate was purified by RP-HPLC and the pure fractions were pooled and lyophilized obtaining ubiquitin-ribose conjugate **4** (0.21 mg, 0.023 µmol, 10.2 %) as a red powder. HRMS: $[C_{408}H_{652}FN_{107}O_{129}S + 7H]^{7+}$ found: 1311.1853, calculated: 1311.1971. $[C_{408}H_{652}FN_{107}O_{129}S + 8H]^{8+}$ found: 1147.2726, calculated: 1147.4225. $[C_{408}H_{652}FN_{107}O_{129}S + 9H]^{9+}$ found: 1020.0854, calculated: 1020.0422. $[C_{408}H_{652}FN_{107}O_{129}S + 10H]^{10+}$ found: 918.0056, calculated: 918.1383. $[C_{408}H_{652}FN_{107}O_{129}S + 11H]^{11+}$ found: 834.5984, calculated: 834.7618.

Rhodamine-(Arg42 → triazole linked vinyl-sulfonate ribose) $Ub_{76}$ (**5**)

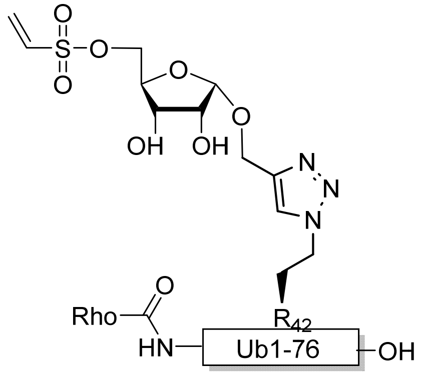

Vinyl sulfonate **20** (5 µL, 0.15 M in DMSO, 0.75 µmol, 3.3 eq.) was added to Arg42 → azido homoalanine modified rhodamine ubiquitin **21** (M + $H^+$ = 8874 Da, 2 mg, 0.225 µmol, 1 eq.) dissolved in DMSO (33.3 µL) and TRIS buffer (100 µL, 20 mM TRIS/150 mM NaCl, pH 6.3). To this solution was added 10 µL of freshly prepared click-mixture (1:1:1 v/v/v, $CuSO_4$ (100 mM in $H_2O$): Sodium Ascorbate (600 mM in $H_2O$): TBTA ligand (100 mM in MeCN). The reaction mixture was shaken at 37 °C for 2 hours before the addition of another portion of freshly prepared click mixture (10 µL) and **20** (2 µL, 0.15M in DMSO, 0.3 µmol, 1.3 eq.). After shaking for another hour LC-MS verified complete conversion to the clicked product (mass found: M+$H^+$ = 9151 Da). The conjugate was purified by RP-HPLC and the pure fractions were pooled and lyophilized obtaining ubiquitin-ribose conjugate **5** (0.26 mg, 0.028 µmol, 12.6 %) as a red powder. HRMS: $[C_{408}H_{651}N_{107}O_{129}S + 7H]^{7+}$ found: 1308.2732, calculated: 1308.3400. $[C_{408}H_{651}N_{107}O_{129}S + 8H]^{8+}$ found: 1144.8230, calculated: 1144.9225. $[C_{408}H_{651}N_{107}O_{129}S + 9H]^{9+}$ found: 1017.7263, calculated: 1017.8200. $[C_{408}H_{651}N_{107}O_{129}S + 10H]^{10+}$ found: 916.0435, calculated: 916.1383. $[C_{408}H_{651}N_{107}O_{129}S + 11H]^{11+}$ found: 832.8512, calculated: 832.9436.

Arg42 → triazole linked vinyl-sulfonate ribose) $Ub_{76}$ (**6**) synthesis (no tag on N-terminus)

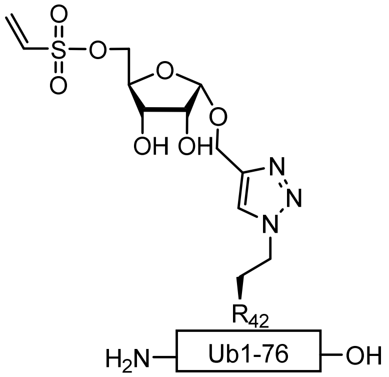

$Ub_{1-76}$ (Arg42 → azido homoalanine, 20 µmol) was prepared on wang resin according to literature procedure[11] of which 8 µmol was treated with TFA/TIS/$H_2O$/Phenol (90.5/2/5/2.5, v/v) for 2.5 hours. The solution was filtrated in an ice-cold solution of $Et_2O$:pentane (1:1, v/v) before the formed precipitate was centrifuged (5 min, 3500 rpm) and the supernatant decanted. The pellet was subsequently dried with $N_2$, taken up in warm DMSO (1 ml) and diluted with warm water before purified by RP-HPLC. Pure fractions were pooled and lyophilized affording $Ub_{1-76}$ (Arg42 → azido homoalanine) as a white powder. Deconvoluted mass found: ($M + H^+ = 8517$). Additionally, CuAAC was performed. $Ub_{1-76}$ (Arg42 → azido homoalanine, 15 mg, 1.76 µmol) was dissolved in DMSO (250 µL) before diluted in TRIS buffer (750 µL, 20 mM TRIS, 150 mM NaCl, pH 6.3). To this mixture was added vinyl sulfonate ribose **20** (40 µL, 0.15 M in DMSO, 6 µmol, 3.4 eq.) followed by 40 µL freshly prepared click mix (1:1:1 v/v/v, $CuSO_4$ (100 mM in $H_2O$): Sodium Ascorbate (600 mM in $H_2O$): TBTA ligand (100 mM in MeCN). After 30 min of shaking at 37°C, another portion of **20** (20 µL, 0.15M in DMSO, 3 µmol, 1.7 eq.) and 60 µL click mix were added. Full conversion ($M + H^+ = 8794$ Da) was observed after 3 hours, monitored by LC-MS, and the mixture was purified by RP-HPLC. Pure fractions were pooled and lyophilized to obtain vinyl sulfonate ubiquitin-ribose conjugate **6** (5.1 mg, 0.57 µmol, 32.2 %). HRMS: $[C_{387}H_{639}N_{105}O_{125}S + 7H]^{7+}$ found: 1257.3403, calculated: 1257.4343, $[C_{387}H_{639}N_{105}O_{125}S + 8H]^{8+}$ found: 1100.2778, calculated: 1100.3802, $[C_{387}H_{639}N_{105}O_{125}S + 9H]^{9+}$ found: 978.1292, calculated: 978.2266, $[C_{436}H_{699}N_{113}O_{138}S_2 + 10H]^{10+}$ found: 880.3937, calculated: 880.5043, $[C_{436}H_{699}N_{113}O_{138}S_2 + 11H]^{11+}$ found: 800.3965, calculated: 800.5508

Rhodamine-$PEG_2$-Lys(biotin)-$PEG_2$-(Arg42 → triazole linked vinyl-sulfonate ribose) $Ub_{76}$ (**7**)

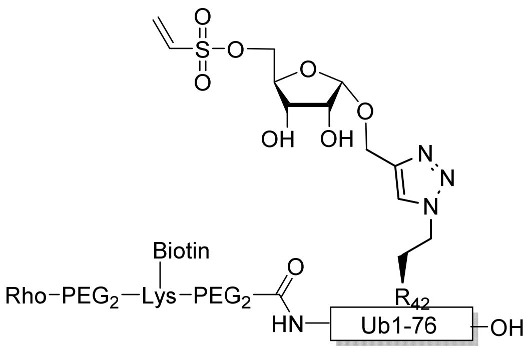

$Ub_{1-76}$ (Arg42 → azido homoalanine, 20 µmol) was prepared on wang resin according to literature procedure[11] and subsequently treated with HOBt (6.7 mg, 50 µmol, 5 eq.), HBTU (19 mg, 50 µmol, 5 eq.) and Fmoc-$PEG_2$-COOH (19.3 mg, 50 µmol, 5 eq.) in DMF (2 ml). After 5 min of shaking, DiPEA (26 µL, 150 µmol, 15 eq.) was added. The reaction mixture was shaken overnight, after which a test cleavage confirmed full conversion of the conjugation (mass found: ($M + H^+$ = 8884 Da). The resin was washed with DMF and DCM before deprotecting the Fmoc with 20% piperidine in DMF (2 mL, 3 min, 3x). After washing (DMF), Fmoc-Lys(Biotin)-OH (40.4 mg, 68 µmol, 6.8 eq.) dissolved in NMP, PyBOP (35.4 mg, 68 µmol, 6.8 eq.) and DIPEA (26 µL, 150 µmol, 15 eq.) were added. After 1 hour of shaking a test cleavage was performed ($M + H^+$ = 9238 Da). Additionally, the Fmoc-group was removed as described before and another Fmoc-$PEG_2$-COOH (50 µmol) coupling was performed. After 1 hour a test cleavage confirmed successful conjugation ($M + H^+$ = 9382 Da). The Fmoc was deprotected and subsequently the final coupling was performed. DiBoc-Rhodamine (28.73 mg, 50 µmol, 5 eq) was added together with PyBOP (26 mg, 50 µmol, 5 eq.) and DiPEA (26 µL, 150 µmol, 15 eq.) and after 1 hour of shaking formation of the biotin and rhodamine N-terminus modified ubiquitin was verified ($M + H^+$ = 9518 Da). The resin was treated with TFA/TIS/$H_2O$/Phenol (90.5/2/5/2.5, v/v) for 2.5 hours before filtrated in an ice-cold solution of $Et_2O$:pentane (1:1, v/v). The precipitate formed was centrifuged (5 min, 3500 rpm) and the supernatant decanted. The pellet was subsequently dried with $N_2$, taken up in warm DMSO (1 ml) and diluted with warm water before purification by RP-HPLC. Pure fractions were pooled and lyophilized to the obtain rhodamine-biotinylated ubiquitin (5.4 mg, 0.57 µmol, 5.7 %) compound as a red powder.

Additionally, the click reaction was performed utilizing this rhodamine-biotinylated Arg42 → azido homoalanine ubiquitin conjugate (1.5 mg, 0.16 µmol, 1 eq.) dissolved in DMSO (25 µL) and TRIS buffer (75 µL, 20 mM TRIS/150 mM NaCl, pH 6.3). To initiate the reaction vinyl sulfonate **20** (5 µL, 0.15 M in DMSO, 0.75 µmol, 4.7 eq.) and 15 µL freshly prepared click mix (1:1:1 v/v/v, $CuSO_4$ (100 mM in $H_2O$): Sodium Ascorbate (600 mM in $H_2O$): TBTA ligand (100 mM in MeCN) were added. The reaction was shaken for 1 hour at 37 °C reaching full conversion as observed by LC-MS (mass found: $M + H^+$ = 9796 Da). The conjugate was subsequently purified by RP-HPLC and the pure fractions were pooled and lyophilized obtaining ubiquitin-ribose conjugate **7** (0.36 mg, 0.037 µmol, 23.3 %) as a red powder. HRMS: $[C_{436}H_{699}N_{113}O_{138}S_2 + 7H]^{7+}$ found: 1400.4207, calculated: 1400.4514. $[C_{436}H_{699}N_{113}O_{138}S_2 + 8H]^{8+}$ found: 1225.4253, calculated: 1225.5232. $[C_{436}H_{699}N_{113}O_{138}S_2 + 9H]^{9+}$ found: 1089.4050, calculated: 1089.4622. $[C_{436}H_{699}N_{113}O_{138}S_2 + 10H]^{10+}$ found: 980.5313, calculated: 980.616. $[C_{436}H_{699}N_{113}O_{138}S_2 + 11H]^{11+}$ found: 891.4427, calculated: 891.5636.

Rhodamine-(Arg42 → azido homoalanine) $Ub_{76}$ (**21**)

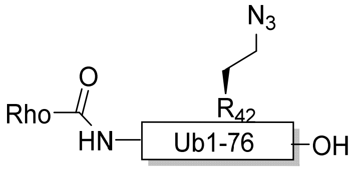

$Ub_{1-76}$ (Arg42 → azido homoalanine) was prepared on wang resin according to literature procedure[11] and subsequently treated with HOBT (6.7 mg, 50 µmol, 5 eq.), HBTU (19 mg, 50 µmol, 5 eq.) and DiBoc-Rhodamine (28.7 mg, 50 µmol, 5 eq.) in DMF (2 ml). After 5 min of shaking, DIPEA (26 µL, 150 µmol, 15 eq.) was added. The reaction mixture was shaken overnight, after which a test cleavage confirmed full conversion of the conjugation (mass found: ($M + H^+$ = 8874 Da). The resin was treated with TFA/TIS/$H_2O$/Phenol (90.5/2/5/2.5, v/v) for 2.5 hours before filtrated in an ice-cold solution of $Et_2O$:pentane (1:1, v/v). The precipitate formed was centrifuged (5min, 3500 rpm) and the supernatant decanted. The pellet was subsequently dried with $N_2$, taken up in warm DMSO (1 mL) and diluted with warm water before purified by RP-HPLC. Pure fractions were pooled and lyophilized affording ubiquitin conjugate **21** (9.6 mg, 1.08 µmol, 10.8 %) as a red powder. Deconvoluted mass found: ($M + H^+$ = 8874 Da).

### HRMS data of probes (1-7)

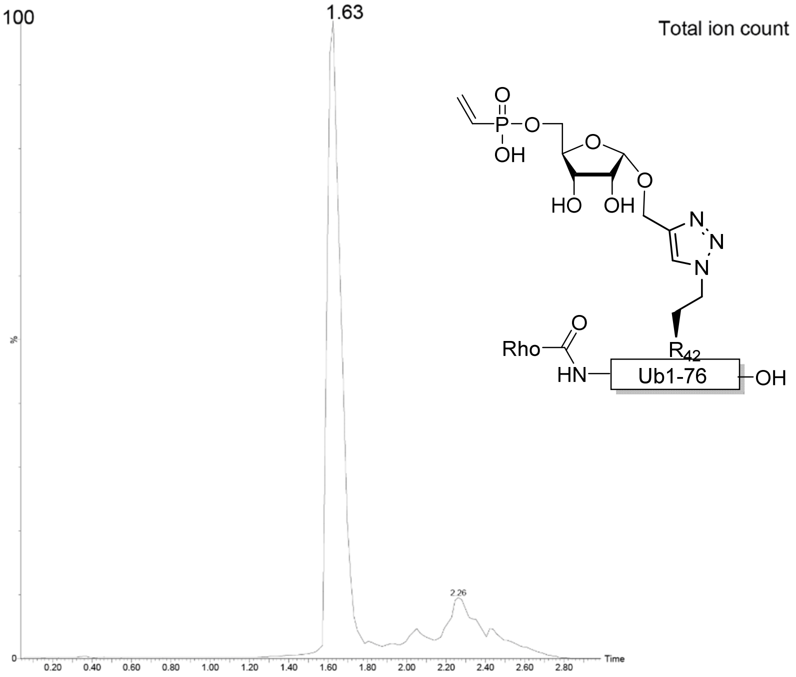

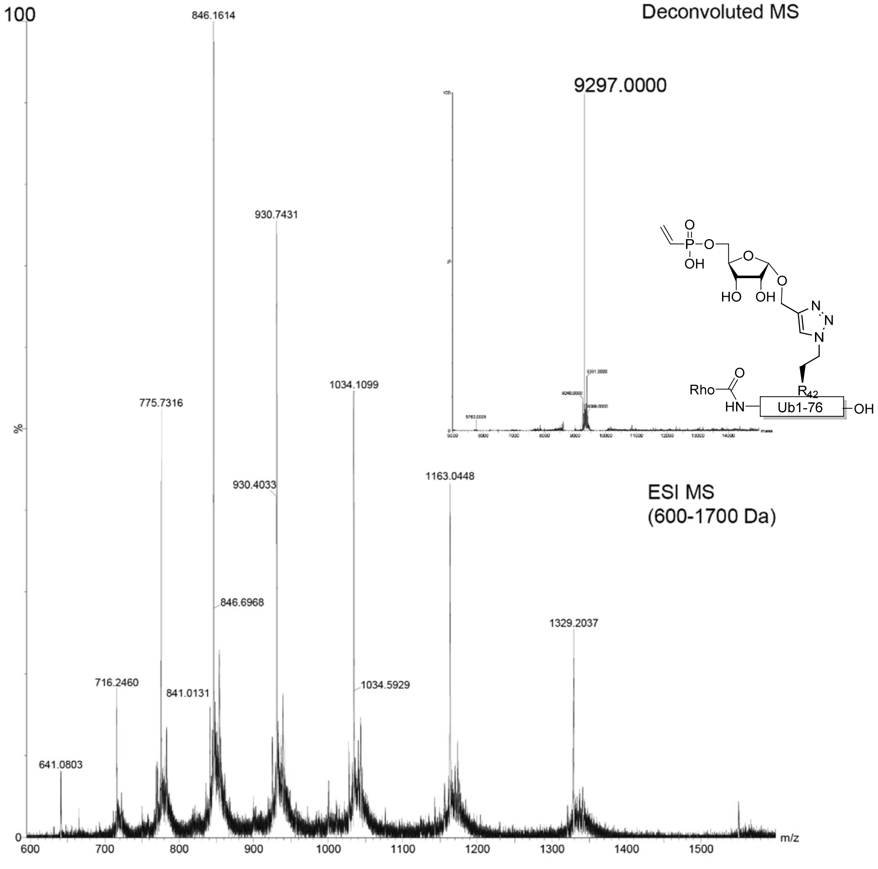

**Supplementary Figure 14.** HRMS spectra of Probe **1**.

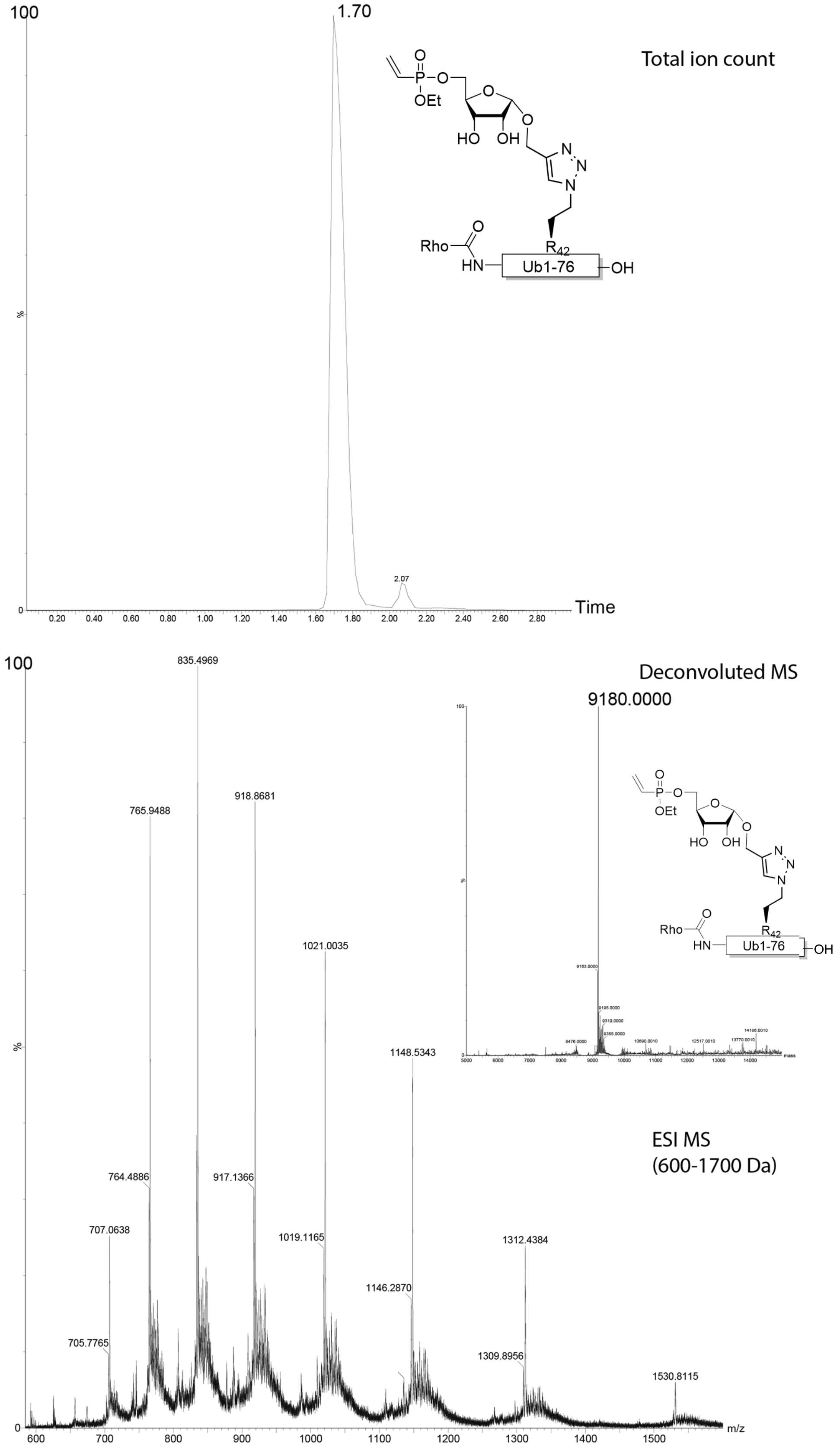

**Supplementary Figure 15.** HRMS spectra of Probe **2**.

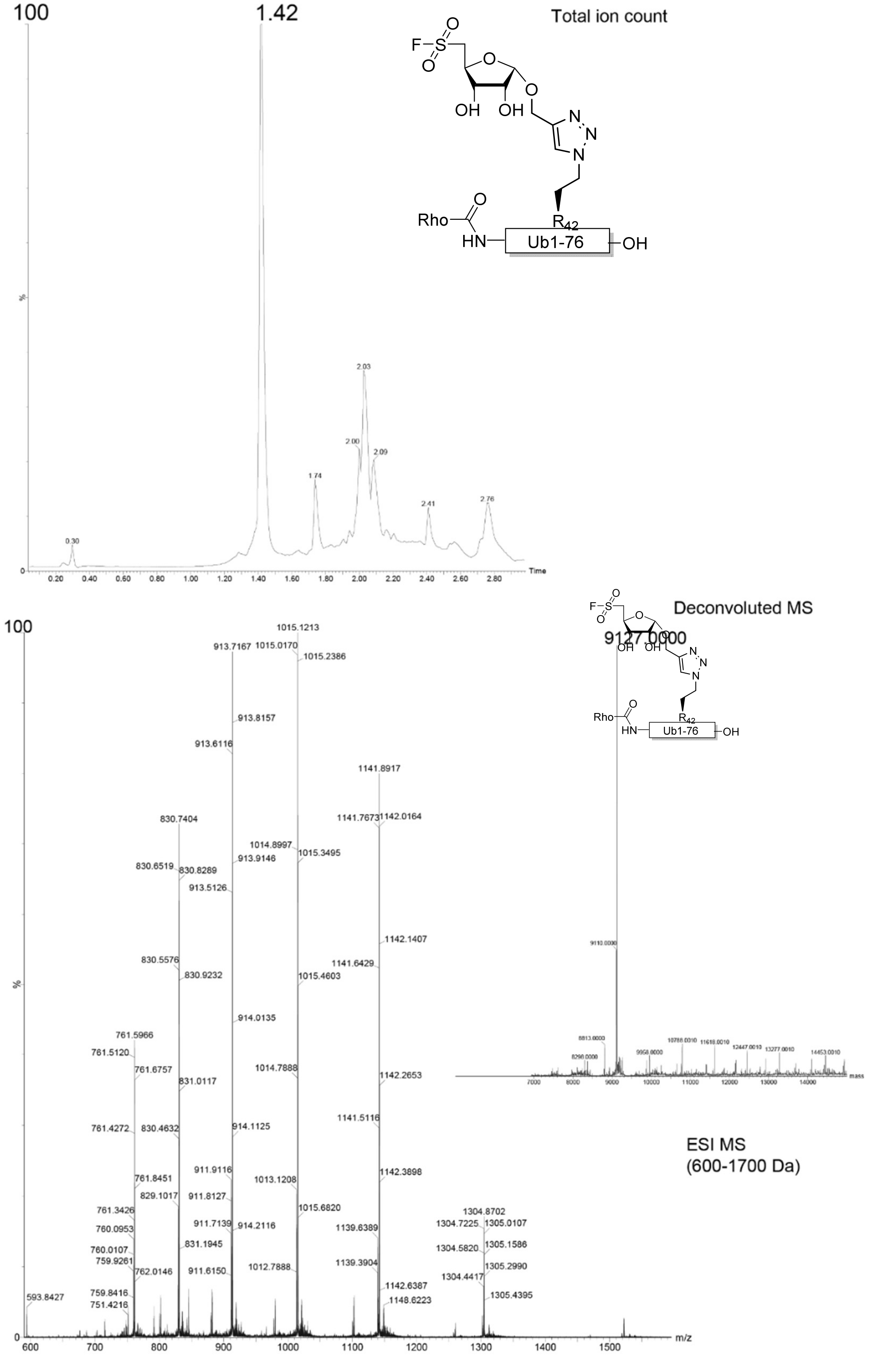

**Supplementary Figure 16.** HRMS spectra of Probe **3**.

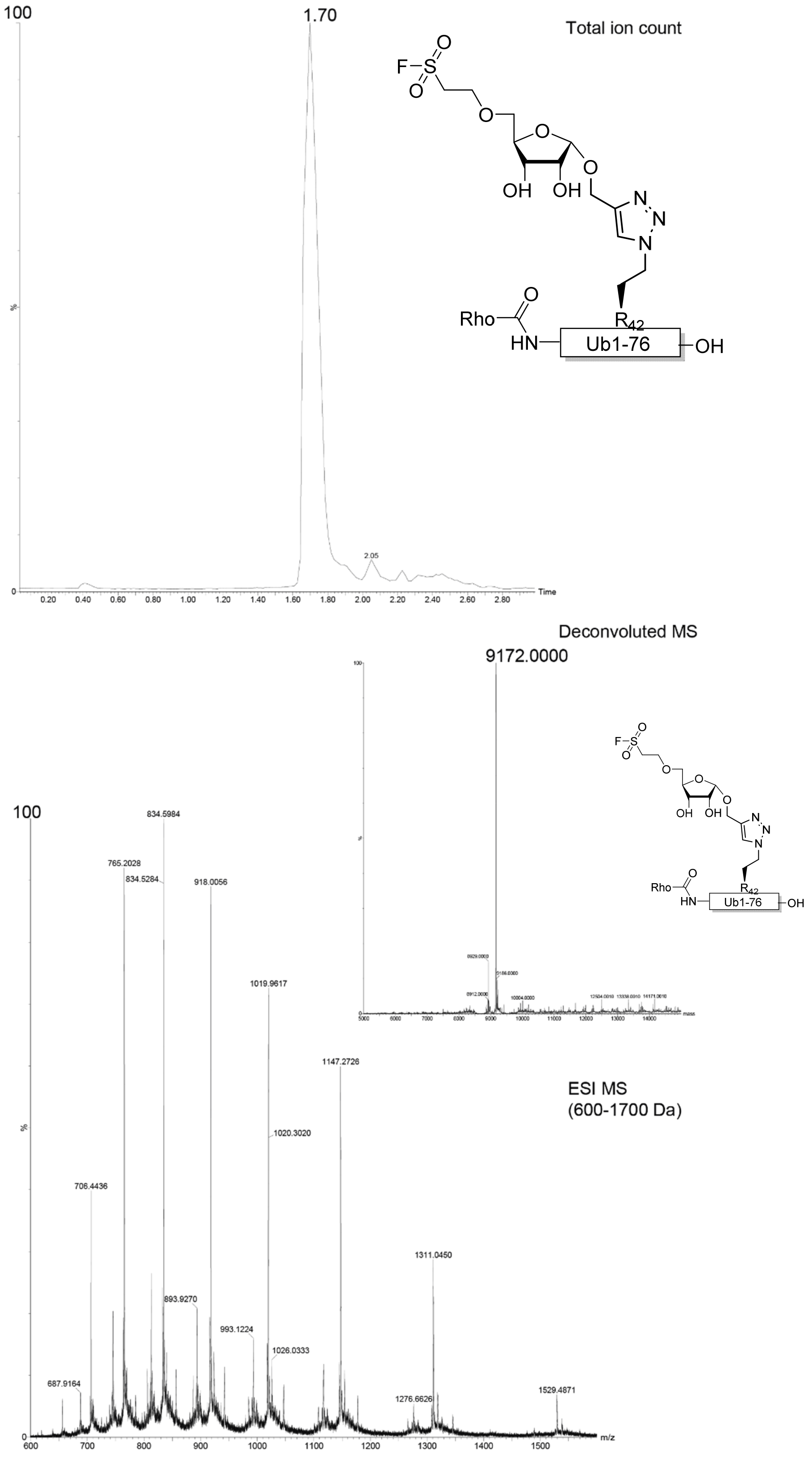

**Supplementary Figure 17.** HRMS spectra of Probe **4**.

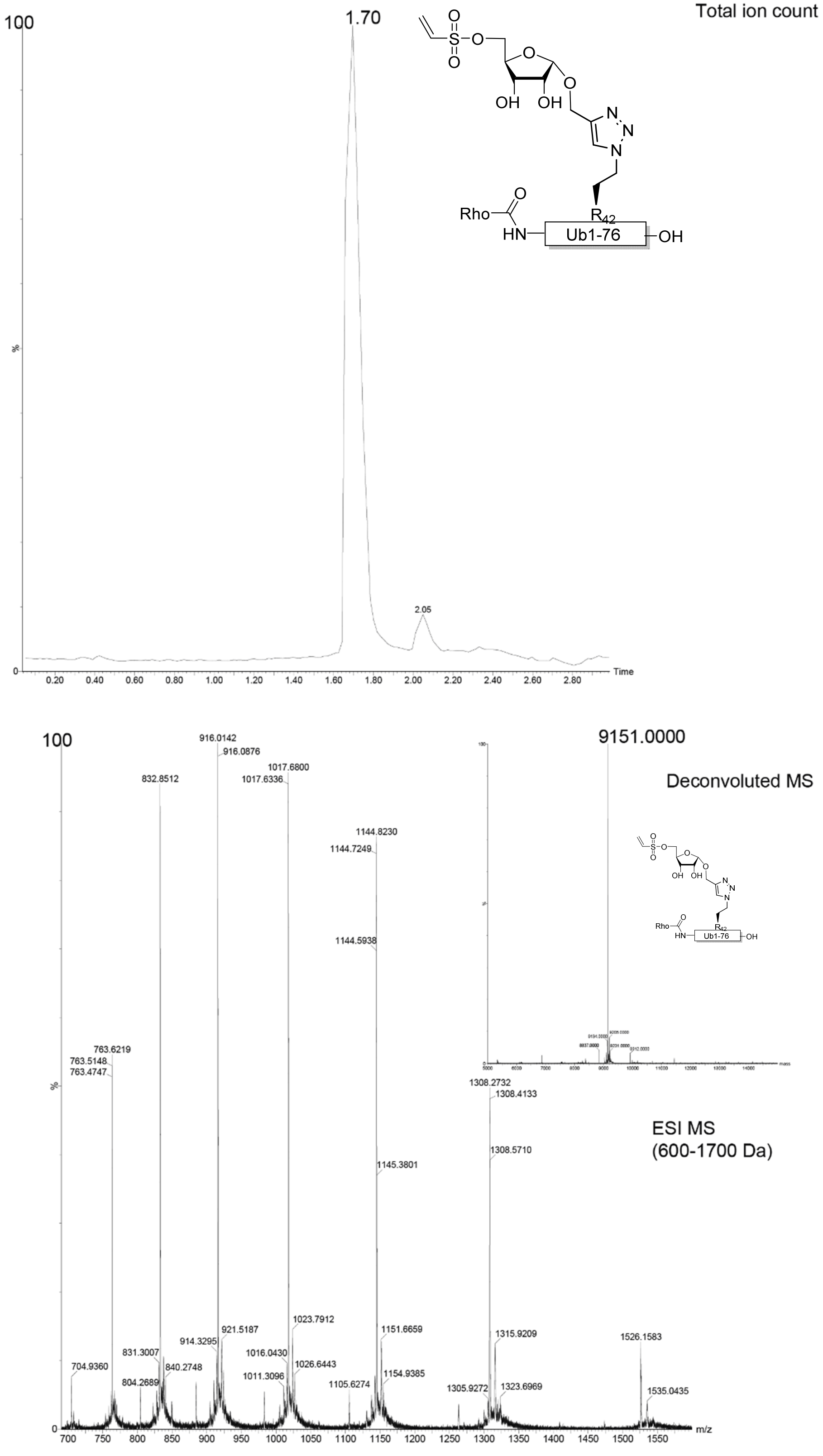

**Supplementary Figure 18.** HRMS spectra of Probe **5**.

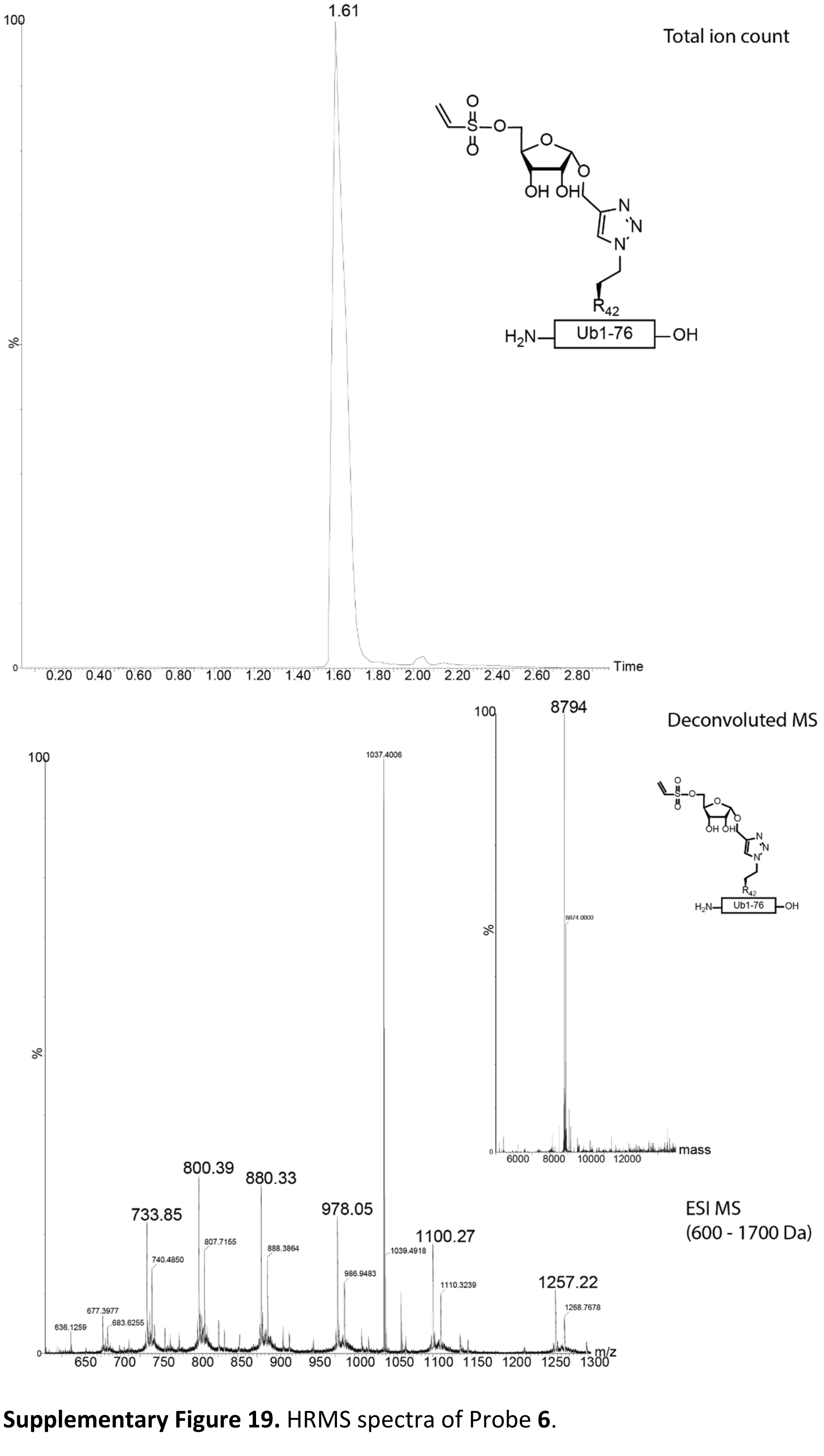

**Supplementary Figure 19.** HRMS spectra of Probe **6**.

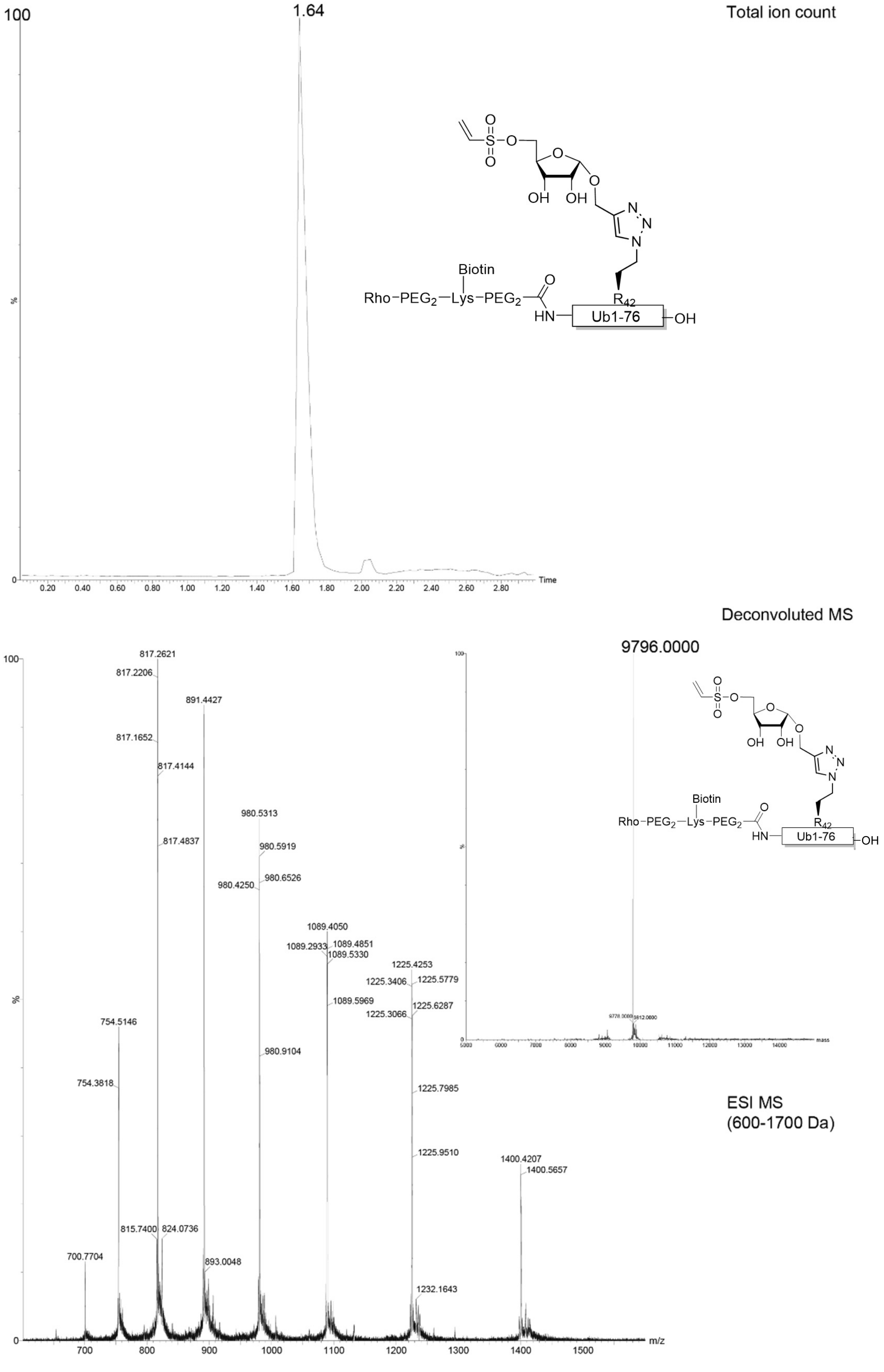

**Supplementary Figure 20.** HRMS spectra of Probe **7**.

### Ribosisde Synthesis

α-1-O-Propargyl-ribofuranose (**8**)

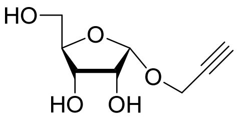

To a suspension of D-ribose (6 g, 40 mmol) in propargyl alcohol (100 mL) under argon atmosphere was added acetyl chloride (2.0 mL, 28 mmol, 0.7 eq.). The reaction was stirred at rt for 0.5 hours and neutralized by addition of pyridine (0.5 mL). The reaction mixture was concentration *in vacuo* and purified by silica column chromatography (0 → 100 % EtOAc in Heptane). Column chromatography was conducted multiple times separating the α- and ß-anomer, obtaining α-glycosylated D-ribose **8** (1.2 g, 64 mmol, 16 %) as a colorless oil. Spectroscopic data was in agreement with literature.[12] $^{1}$H NMR (300 MHz, $D_2O$), δ: 5.39 (d, *J* = 4.4 Hz, 1H), 4.56 – 4.38 (m, 2H), 4.30 – 4.20 (m, 2H), 4.16 (dd, *J* = 6.4, 3.1 Hz, 1H), 3.83 (qd, *J* = 12.3, 3.9 Hz, 2H), 3.10 (t, *J* = 2.3 Hz, 1H). $^{13}$C-NMR (75 MHz, $D_2O$), δ: 100.6, 85.1, 79.7, 76.3, 71.4, 69.6, 61.7, 55.0.

α-1-O-Propargyl-2',3'-di-*O*-*iso*-butyryl-ribofuranose (**9**)

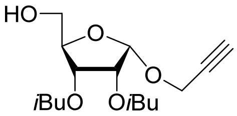

Ribose **8** (253 mg, 1.34 mmol) was co-evaporated with pyridine and subsequently placed under argon atmosphere and dissolved in anhydrous pyridine (6.72 mL). TBDMS-Cl (421 mg, 2.8 mmol, 2.08 eq) was added and the reaction mixture was stirred at rt for 30 minutes. To this solution was added *iso*butyric anhydride (1.03 mL, 5.59 mmol, 4.16 eq) followed by overnight stirring. Next, the reaction mixture was concentrated *in vacuo* and taken up in EtOAc (20 mL). The organic layer was washed with 5 % (w/v) citric acid (10 mL) and $H_2O$ (10 mL), dried over $MgSO_4$, filtrated and concentrated *in vacuo*. Subsequently, the residue was taken up in MeCN: $H_2O$ (14 mL, 4:1, v/v) and *p*TsOH monohydrate (266 mg, 1.4 mmol, 1.04 eq) was added. The reaction mixture was stirred for 1 hour at rt before TLC indicated full conversion. Subsequently, $Et_3N$ (300 µL, 2.13 mmol, 1.59 eq.) was added to neutralize the reaction followed by concentration *in vacuo*. The residue was purified by silica column chromatography (0 → 60 % EtOAc in heptane) to obtain *i*Bu protected D-ribose **9** (258 mg, 0.79 mmol, 59 %) as a colorless oil. $^{1}$H NMR (300 MHz, $CDCl_3$) δ 5.48 (d, *J* = 4.4 Hz, 1H), 5.22 (dd, *J* = 7.2, 3.7 Hz, 1H), 4.97 (dd, *J* = 7.3, 4.4 Hz, 1H), 4.29 (dd, *J* = 2.4, 0.7 Hz, 2H), 4.17 (q, *J* = 3.5 Hz, 1H), 3.81 (qd, *J* = 12.1, 3.4 Hz, 2H), 2.61 (pd, *J* = 7.0, 3.9 Hz, 2H), 2.40 (t, *J* = 2.4 Hz, 1H), 1.23 – 1.14 (m, 12H). $^{13}$C NMR (75 MHz, $CDCl_3$) δ 176.8, 176.2, 99.1, 82.8, 79.0, 74.6, 71.3, 69.9, 62.3, 54.9, 33.9, 33.7, 19.0, 18.9, 18.9.

α-1-O-Propargyl-2',3'-di-*O*-*iso*-butyryl-5'-O-vinyl-phosphate-ribofuranose (**10**)

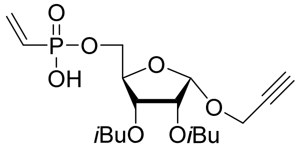

To a flame dried flask were added **9** (55 mg, 0.18 mmol) and DCC (148 mg, 0.72 mmol, 4 eq). The reagents were co-evaporated with anhydrous pyridine (1 mL) and subsequently placed under argon atmosphere. Vinylphosphonic acid (34 µL, 0.43 mmol, 2.4 eq) was added and the reaction mixture stirred overnight at 50 °C. Upon full conversion (as indicated by TLC), $H_2O$ was added (5 mL) and the mixture was cooled on ice for 15 min before filtration over a syringe with filter. The precipitate was washed with $H_2O$ (2 mL) before concentration of the filtrate *in vacuo* and impregnation onto Celite®. Silica column chromatography (0 → 60 % MeOH in DCM) afforded phosphorylated ribose **10** (47 mg, 0.11 mmol, 63 %) as a colorless oil. $^1$H NMR (300 MHz, MeOD) δ 6.27 – 6.07 (m, 2H), 6.07 – 5.86 (m, 1H), 5.42 (d, *J* = 4.5 Hz, 1H), 5.29 (dd, *J* = 6.9, 3.1 Hz, 1H), 4.99 (dd, *J* = 7.0, 4.5 Hz, 1H), 4.34-4.22 (m, 3H), 4.10 (dd, *J* = 6.1, 3.2 Hz, 2H), 2.83 (t, *J* = 2.4 Hz, 1H), 2.59 (dhept, *J* = 10.7, 7.0 Hz, 2H), 1.24 – 1.08 (m, 12H). $^{13}$C NMR (75 MHz, Methanol-$d_4$) δ 177.8, 177.3, 134.2, 132.6, 100.3, 82.5, 80.0, 75.9, 72.5, 71.1, 65.6, 55.5, 49.3, 35.0, 34.9, 19.4, 19.1. $^{31}$P NMR (121 MHz, MeOD) δ 14.5. This compound was used in the formation of compound **1** and *i*Bu protection was removed after CuAAC as described in the experimental of probe **1**.

α-1-O-Propargyl-2',3'-di-*O*-*iso*-butyryl-5'-O-vinyl-ethoxy-phosphate-ribofuranose (**11**)

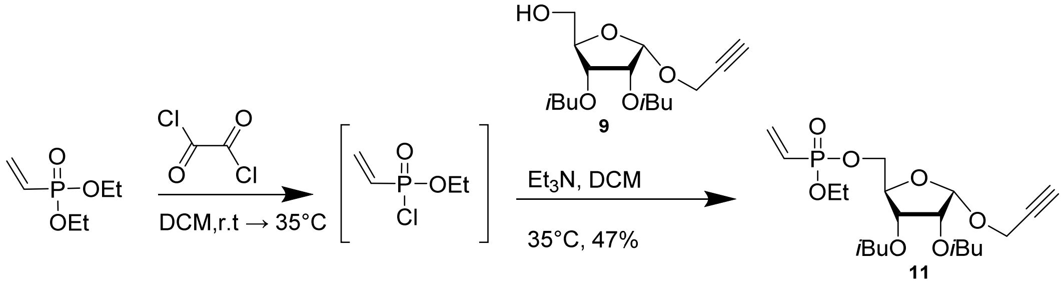

Compound **11** was synthesized based on a procedure by Liautard *et al.*[13] To a flame dried flask under argon atmosphere was added diethyl vinylphosphonate (189 µL, 1.2 mmol) and a solution of oxalyl chloride (115 µL, 1.3 mmol, 1.1 eq.) in anhydrous DCM (5 mL). The reaction was stirred overnight stirred at rt. Subsequently, the temperature was increased to 35°C and stirred for an additional 24 h. The conversion was monitored by $^1$H- and $^{31}$P-NMR, which indicated consumption of the starting material. The formed ethyl vinylphosphonyl chloride reagent was subsequently concentrated in *vacuo*, and 120 mg (0.78 mmol) was transferred to a flame dried flask. The reagent was co-evaporated with anhydrous toluene (2 mL) before placed under argon atmosphere and dissolved in anhydrous DCM (1.3 mL). This solution was added dropwise to a flame dried flask containing ribose **9** (160 mg, 0.5 mmol) and $Et_3N$ (144 µL, 1.0 mmol, 2 eq.). After addition, the temperature was increased to 35 °C and the reaction stirred for 48 h. The reaction mixture was diluted with $H_2O$ (2 mL) and DCM (10 mL) before transferred to a separatory funnel. The water layer was then extracted three times with DCM and the combined organics were dried over $MgSO_4$, filtrated and concentrated *in vacuo*. Purification by silica column chromatography (30 → 100 % EtOAc in heptane) yielded phosphorylated ribose **11** (109 mg, 0.24 mmol, 47 % yield). $^{1}$H-NMR (300 MHz, Chloroform-*d*), δ: 6.39 – 6.18 (m, 1.5H), 6.14 – 5.93 (m, 1.5H), 5.41 (dd, *J* = 5.8, 4.5 Hz, 1H), 5.22 (ddd, *J* = 9.2, 7.1, 3.2 Hz, 1H), 4.91 (ddd, *J* = 7.1, 4.4, 3.8 Hz, 1H), 4.28 – 4.16 (m, 3H).4.15 – 4.01 (m, 2H), 2.56 (hept, *J* = 7.0 Hz, 2H), 2.37 (t, *J* = 2.4 Hz, 1H), 1.31 (t, *J* = 7.2 Hz, 3H), 1.15 (dt, *J* = 7.0, 4.7 Hz, 12H). $^{13}$C-NMR (75 MHz, $CDCl_3$), δ: 176.3, 175.9, 136.2, 126.5, 124.1, 98.9, 80.8, 80.7, 80.7, 80.7, 78.9, 74.4, 70.8, 69.5, 64.7, 62.2, 54.8, 33.8, 18.9, 16.3. $^{31}$P-NMR (121 MHz, $CDCl_3$), δ: 18.2.

α-1-O-Propargyl-5'-O-vinyl-ethoxy-phosphate-ribofuranose (**12**)

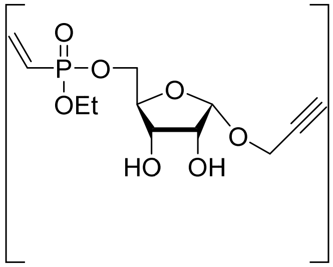

To a stirring solution of ribose **11** (7 mg, 0.016 mmol) in anhydrous MeOH (0.4 mL) was added NaOEt (2.66 mg, 0.039 mmol, 2.5 eq.) After TLC indicated full consumption of the starting material 4M HCl in dioxane (10 µL) was added and the reaction reached pH = 7. The mixture was diluted with toluene (2 mL) before concentrated *in vacuo* to obtain **12**. This ribose molecule was used as a crude in the synthesis of probe **2**. $^{1}$H NMR (300 MHz, $CDCl_3$) δ 6.40 – 6.30 (m, 1H), 6.28 – 6.22 (m, 1H), 6.14 – 6.02 (m, 1H), 5.27 (dd, *J* = 4.5, 1.2 Hz, 1H), 4.43 – 4.28 (m, 2H), 4.20 – 4.08 (m, 5H), 4.07 – 4.00 (m, 1H), 2.46 (t, *J* = 2.4 Hz, 1H), 1.59 (s, 2H), 1.34 (t, *J* = 7.1 Hz, 3H).

α-1-O-Propargyl-2',3'-*O*-*iso*propylidene-ribofuranose (**13**)

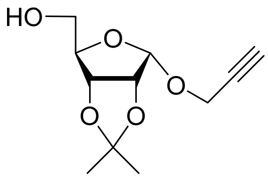

To an ice cooled solution of ribose **8** (174 mg, 0.93 mmol) in acetone (4.65 mL) was added *p*-Toluene sulfonic acid monohydrate (5.5 mg, 0.029 mmol, 0.03 eq.) followed by 2'2'dimethoxy propane (124 µl, 1.02 mmol, 1.1 eq.). After 30 min of stirring at rt TLC indicated full conversion. The reaction mixture was subsequently neutralized by the addition of $Et_3N$ (20 µL) and concentrated *in vacuo*. Purification by silica column chromatography (20 → 70 % EtOAc in heptane) yielded ribose **13** (143 mg, 0.63 mmol, 68 % yield) as a colorless oil. $^{1}$H NMR (300 MHz, $CDCl_3$) δ 5.26 (d, *J* = 3.4 Hz, 1H), 4.74 – 4.64 (m, 2H), 4.36 (dd, *J* = 2.4, 1.7 Hz, 2H), 4.25 – 4.18 (m, 1H), 3.89 – 3.66 (m, 2H), 2.42 (t, *J* = 2.4 Hz, 1H), 1.73 (s, 1H), 1.58 (s, 3H), 1.55 (s, 3H). $^{13}$C NMR (75 MHz, $CDCl_3$) δ 115.9, 99.7, 82.2, 81.2, 80.4, 79.1, 77.6, 77.2, 76.7, 74.8, 62.8, 54.8, 26.1, 25.9.

α-1-O-Propargyl-2’,3’-*O-iso*propylidene-5’-O-mesylate-ribofuranose (**14**)

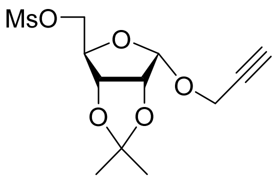

Synthesis of the fluorosulfonate warhead and precursors has previously been described in literature. [14–17]

Ribose **13** (400 mg, 1.75 mmol) was placed under $N_2$ atmosphere before dissolved in anhydrous DCM (5.83 mL). $Et_3N$ (292 µL, 2.1 mmol, 1.2 eq.) was added and the mixture was cooled on ice. Subsequently, mesyl-chloride (188 µL, 2.1 mmol, 1.2 eq.) was added dropwise and the reaction was stirred for 1h allowing to attain rt. $H_2O$ (5 mL) was added to dilute the mixture followed by an extraction with DCM (10 mL). The organic layer was separated and washed with brine (5 mL, 2x), dried over $MgSO_4$, filtrated and concentrated *in vacuo*. The crude mesylate was purified by silica column chromatography (10 → 70 % EtOAc in Heptane) to obtain ribose **14** (497 mg, 1.62 mmol, 93 %) as a colorless oil. $^1$H NMR (300 MHz, $CDCl_3$) δ 5.23 (d, *J* = 4.3 Hz, 1H), 4.77 – 4.57 (m, 2H), 4.41 – 4.23 (m, 5H), 3.04 (s, 3H), 2.42 (t, *J* = 2.4 Hz, 1H), 1.55 (s, 3H), 1.34 (s, 3H). $^{13}$C NMR (75 MHz, $CDCl_3$) δ 116.2, 99.9, 80.8, 80.1, 79.3, 75.0, 69.1, 56.9, 37.7, 26.0, 25.9. $^{13}$C NMR (75 MHz, Chloroform-*d*) δ 116.1, 99.8, 80.7, 80.0, 79.2, 78.8, 74.9, 69.0, 55.0, 37.6, 26.0, 25.8.

α-1-O-Propargyl-2’,3’-*O-iso*propylidene-5’-C-thioacetate-ribofuranose (**15**)

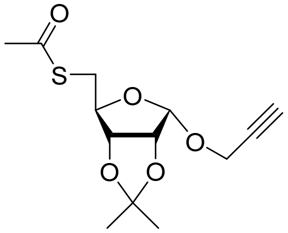

To a suspension of $Cs_2CO_3$ (207 mg, 0.64 mmol, 0.65 eq.) in anhydrous DMF (5 mL) under argon atmosphere was added thioacetic acid (86 µL, 1.20 mmol, 1.23 eq.). The mixture was stirred for 30 min at rt before Ribose **14** (300 mg, 0.98 mmol) dissolved in anhydrous DMF (2 mL) was added. The reaction mixture was covered in aluminium foil and stirred overnight at 50 °C. The reaction mixture was concentrated *in vacuo* and the residue taken up in EtOAc (10 mL). The organic layer was washed with 5 % aq. $NaHCO_3$ (8 mL) and $H_2O$ (5 mL), dried over $MgSO_4$ and concentrated *in vacuo*. The crude was purified by silica column chromatography (0 → 50 % EtOAc in Heptane) to obtain **15** (246 mg, 0.86 mmol, 88 %) as a colorless oil. $^1$H NMR (300 MHz, $CDCl_3$) δ 5.14 (d, *J* = 4.4 Hz, 1H), 4.65 (dd, *J* = 7.1, 4.4 Hz, 1H), 4.34 (dd, *J* = 7.1, 3.4 Hz, 1H), 4.27 (dd, *J* = 3.7, 2.4 Hz, 2H), 4.25 – 4.14 (m, 1H), 3.12 (qd, *J* = 13.9, 5.9 Hz, 2H), 2.35 (t, *J* = 2.4 Hz, 1H), 2.32 (s, 3H), 1.51 (s, 3H), 1.28 (s, 3H). $^{13}$C NMR (75 MHz, $CDCl_3$) δ 194.8, 116.2, 99.5, 82.7, 81.1, 80.3, 79.1, 74.8, 54.9, 31.3, 30.7, 26.1, 26.0.

α-1-O-Propargyl-5′-C-fluorosulfonate-ribofuranose (**16**)

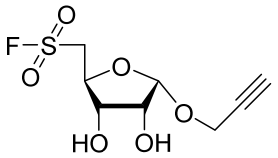

N-Chlorosuccinimide (93 mg, 0.70 mmol, 4 eq.) was dissolved in an ice-cooled solution of MeCN (215 µL) and 2M HCl (43 µL). After stirring for 15 min an ice cooled solution of Ribose **15** (50 mg, 0.18 mmol) in MeCN (46 µL) was added and the reaction was stirred for 30 min allowing to attain rt. TLC confirmed the formation of a lower running spot, which was assumed to be the intermediate sulfonyl chloride. The reaction mixture was diluted with Ether (4 mL) and washed with brine (0.5 mL, 2x). The organic layer was subsequently dried over $MgSO_4$, filtrated and concentrated *in vacuo*. Next, was the colorless sulfonyl chloride intermediate co-evaporated with anhydrous MeCN before dissolved in anhydrous MeCN (875 µL) and placed under argon atmosphere. Subsequently, KF (20 mg, 0.35 mmol, 2 eq.) and 18-Crown-6 (2 mg, 0.008 mmol, 0.05 eq.) were added and the mixture was stirred at rt. After overnight stirring, a $^{19}$F NMR of the crude confirmed the formation of the fluorosulfonate ($^{19}$F: δ = 60 ppm) and the mixture was concentrated *in vacuo*. The residue was purified by silica column chromatography (0 → 100 % EtOAc in Heptane) to obtain **16** (12 mg, 0.047 mmol, 27 %) as a colorless oil. NMR indicated that the formation of **16** was associated with the loss of the isopropylidene group under these conditions. $^{1}$H NMR (300 MHz, $CDCl_3$) δ 5.39 (d, *J* = 4.6 Hz, 1H), 4.40 (dd, *J* = 3.2, 2.4 Hz, 2H), 4.37 – 4.30 (m, 1H), 4.23 (dd, *J* = 6.6, 4.6 Hz, 1H), 3.92 (t, *J* = 6.4 Hz, 1H), 3.81 (ddd, *J* = 14.9, 6.3, 3.7 Hz, 1H), 3.61 (ddd, *J* = 14.9, 8.2, 1.6 Hz, 1H), 2.49 (t, *J* = 2.4 Hz, 1H), 2.41 (s, 2H). $^{13}$C NMR (75 MHz, $CDCl_3$) δ 99.5, 78.1, 76.9, 75.8, 73.1, 70.0, 55.3, 53.4. $^{19}$F NMR (282 MHz, $CDCl_3$) δ 60.51.

α-1-O-Propargyl-2′,3′-*O*-*iso*propylidene-5′-O-ethanesulfonyl fluoride-ribofuranose (**17**)

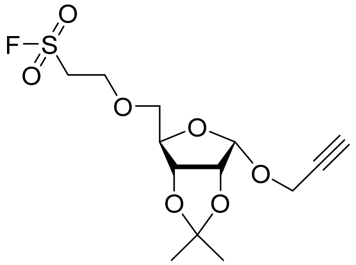

Ribose **13** (60 mg, 0.26 mmol) was co-evaporated with anhydrous MeCN before placed under argon atmosphere and dissolved in anhydrous DCM. $Et_3N$ (109 µL, 0.79 mmol, 3 eq.) and ethenesulfonyl fluoride (43µL, 0.53 mmol, 2 eq.) were added and the reaction mixture was stirred at rt. After 8 hours of stirring, additional ethenesulfonyl fluoride (21.5 µL, 0.26 mmol, 1 eq.) and $Et_3N$ (109 µL, 0.79 mmol, 3 eq.) were added and the reaction was stirred overnight. The reaction mixture was concentrated *in vacuo* and the residue was purified by silica column chromatography (20 → 70 % EtOAc in Heptane) to obtain **17** (40 mg, 0.118 mmol, 45 %) as a colorless oil. $^{1}$H NMR (300 MHz, $CDCl_3$) δ 5.20 (d, *J* = 4.1 Hz, 1H), 4.64 (qd, *J* = 6.9, 3.5 Hz, 2H), 4.30 (dd, *J* = 3.1, 2.4 Hz, 2H), 4.21 (d, *J* = 3.1 Hz, 1H), 4.02 – 3.88 (m, 2H), 3.65 (d, *J* = 3.4 Hz, 2H), 3.63 – 3.54 (m, 2H), 2.36 (t, *J* = 2.4 Hz, 1H), 1.52 (s, 3H), 1.31 (s, 3H). $^{13}$C NMR (75 MHz, $CDCl_3$) δ 115.5, 100.2, 80.8, 80.7, 80.6, 79.1, 74.8, 71.9, 64.7, 55.2, 51.5, 51.3. $^{19}$F NMR (282 MHz, $CDCl_3$) δ 59.6.

α-1-O-Propargyl-5’-O-ethanesulfonyl fluoride-ribofuranose (**18**)

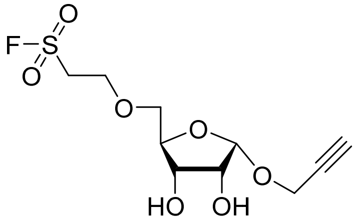

Ribose **17** (20 mg, 0.059 mmol) was dissolved in a mixture of AcOH:$H_2O$ (1 mL, 1:1 v/v) and stirred at 60 °C for 1 hour. Upon consumption of the starting material, the reaction mixture was diluted with toluene (3 mL) and concentrated *in vacuo*. The residue was purified by silica column chromatography (40 → 100 % EtOAc in Heptane) to obtain **18** (9 mg, 0.03 mmol, 51 %) as a colorless oil. $^1$H NMR (300 MHz, $CDCl_3$) δ 5.27 (d, *J* = 4.5 Hz, 1H), 4.36 (dd, *J* = 3.9, 2.4 Hz, 2H), 4.18 – 4.12 (m, 2H), 4.02 – 3.99 (m, 2H), 3.99 – 3.97 (m, 1H), 3.70 (d, *J* = 3.2 Hz, 2H), 3.64 (td, *J* = 5.9, 4.0 Hz, 2H), 2.46 (t, *J* = 2.4 Hz, 1H), 2.13 (s, 2H). $^{19}$F NMR (282 MHz, $CDCl_3$) δ 59.41.

α-1-O-Propargyl-2’,3’-*O-iso*propylidene-5’-O-vinylsulfonate-ribofuranose (**19**)

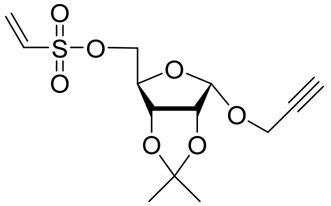

Ribose **13** (150 mg, 0.66 mmol) and $Et_3N$ (463 µL 3.3, mmol, 5 eq.) were placed under argon atmosphere before dissolved in anhydrous DCM (3.3 mL). This mixture was added to a stirring ice-cooled solution of 2-chloroethanesulfonyl chloride (138 µL, 1.3 mmol, 2 eq.) in anhydrous DCM (6.6 mL). The reaction mixture was stirred for 2 hours while cooled by a water-ice bath. Sat. aq. $NaHCO_3$ was added, the organic phase separated and washed with $H_2O$ before dried over $MgSO_4$ and concentrated *in vacuo*. The crude product was purified by silica column chromatography (0 → 60 % EtOAc in Heptane) to yield vinyl sulfonate **19** (201 mg, 0.63 mmol, 96 %) as colorless oil. $^1$H NMR (300 MHz, $CDCl_3$) δ 6.62 – 6.37 (m, 2H), 6.17 (d, *J* = 9.5 Hz, 1H), 5.23 (d, *J* = 4.3 Hz, 1H), 4.70 (dd, *J* = 23.4, 3.8 Hz, 2H), 4.33 (dd, *J* = 4.0, 2.4 Hz, 3H), 4.25 (d, *J* = 3.0 Hz, 2H), 2.42 (t, *J* = 2.4 Hz, 1H), 1.56 (s, 3H), 1.35 (s, 3H). $^{13}$C NMR (75 MHz, $CDCl_3$) δ 132.1, 130.9, 116.0, 99.8, 80.8, 79.9, 79.1, 78.9, 74.9, 69.6, 55.1, 26.0, 25.8.

α-1-O-Propargyl-5’-O-vinylsulfonate-ribofuranose (**20**)

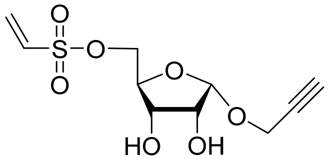

Ribose **19** (117 mg, 0.346 mmol) was dissolved in DCM (3.46 mL) before a 10 % aqueous solution of TFA (9:1 v/v, 3.46 mL) was added. The final TFA concentration was 5 %. After stirring for 1 hour at rt no conversion was observed. Subsequently, TFA (346 µL) was added to increase the TFA concentration to 10 %. The reaction was stirred for 18 h, after which TLC and LC-MS indicated full conversion. The reaction was subsequently diluted with toluene and concentrated *in vacuo*. The crude product was purified by silica column chromatography (0 → 7 % MeOH in DCM) to obtain diol **20** (79 mg, 0.28 mmol, 82 %) as a colorless oil. $^{1}$H NMR (300 MHz, $CDCl_3$) δ 6.56 (dd, *J* = 16.6, 9.6 Hz, 1H), 6.42 (d, *J* = 16.6 Hz, 1H), 6.16 (d, *J* = 9.6 Hz, 1H), 5.27 (d, *J* = 4.3 Hz, 1H), 4.37 – 4.32 (m, 2H), 4.26 (dd, *J* = 6.0, 3.2 Hz, 2H), 4.20 (d, *J* = 3.6 Hz, 1H), 4.17 – 4.11 (m, 1H), 4.01 (dd, *J* = 6.5, 3.6 Hz, 1H), 2.88 (s, 2H), 2.48 (t, *J* = 2.4 Hz, 1H). $^{13}$C NMR (75 MHz, $CDCl_3$) δ 132.1, 132.1, 100.0, 81.9, 78.5, 75.4, 71.3, 70.2, 69.4, 55.0.

### NMR Spectra

$^{1}$H NMR (**8**)

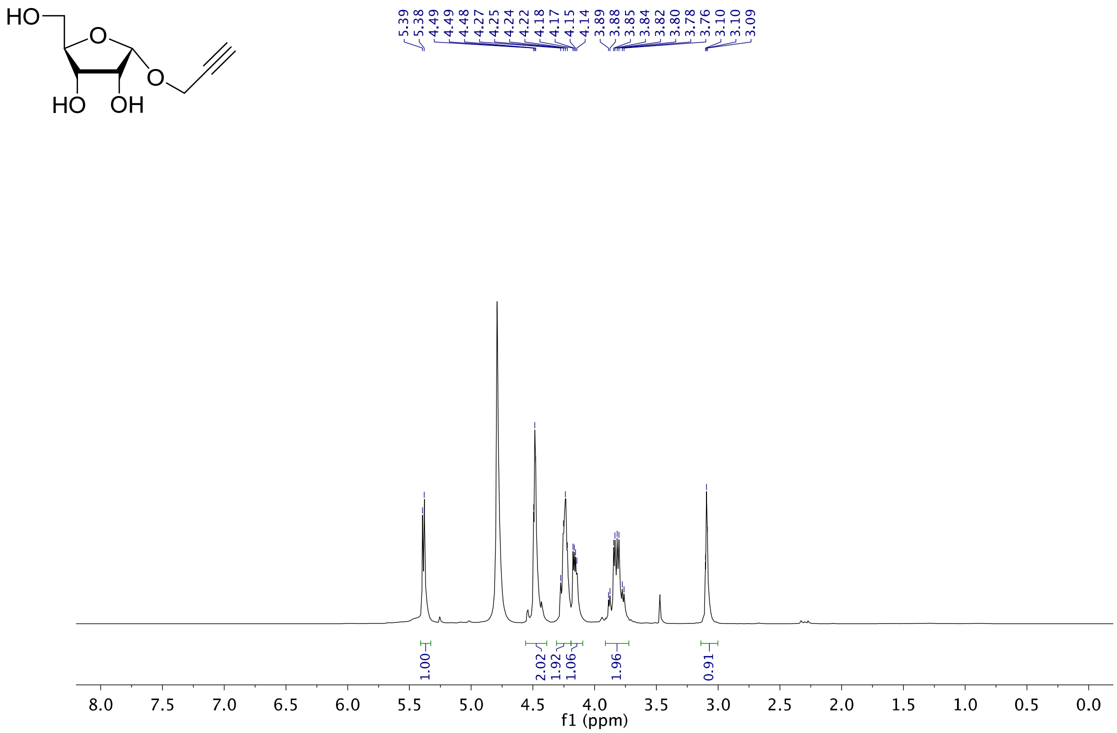

$^{13}$C NMR (**8**)

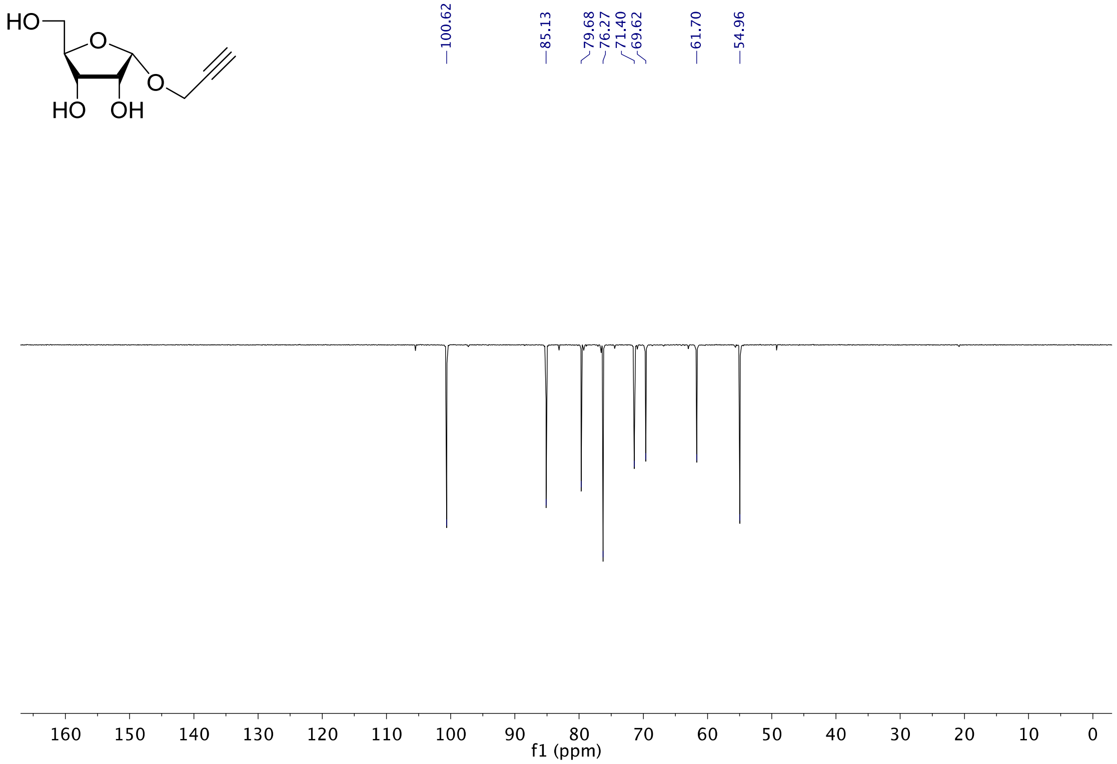

[1]H NMR (**9**)

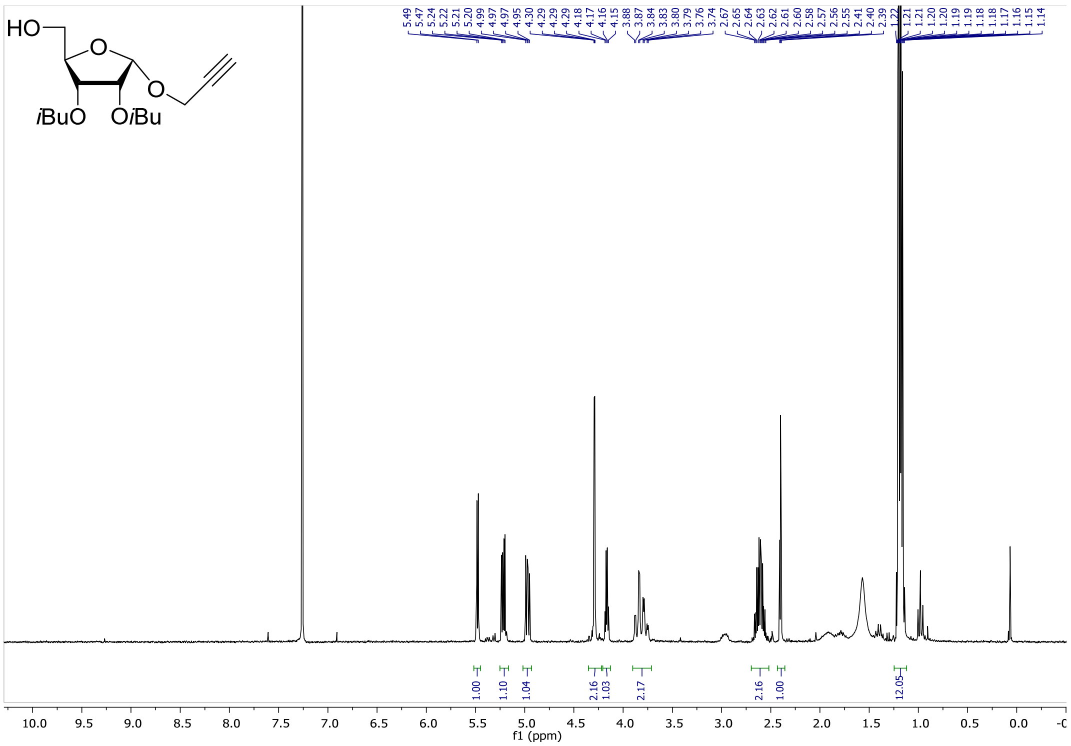

[13]C NMR (**9**)

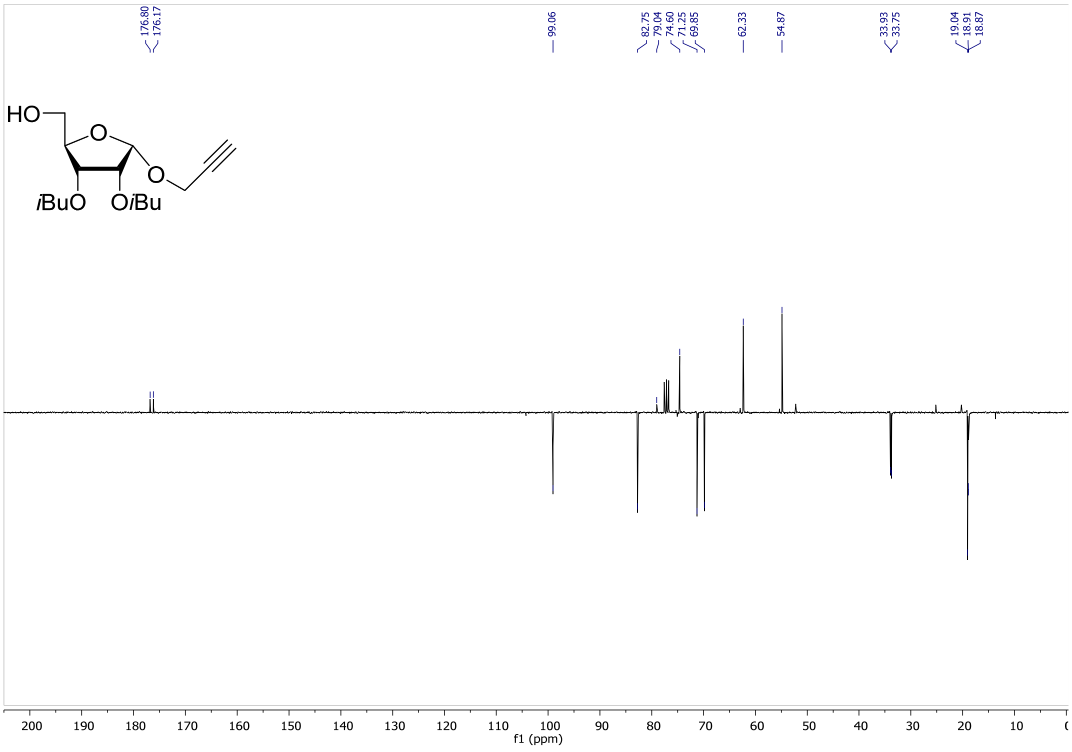

$^1$H NMR (**10**)

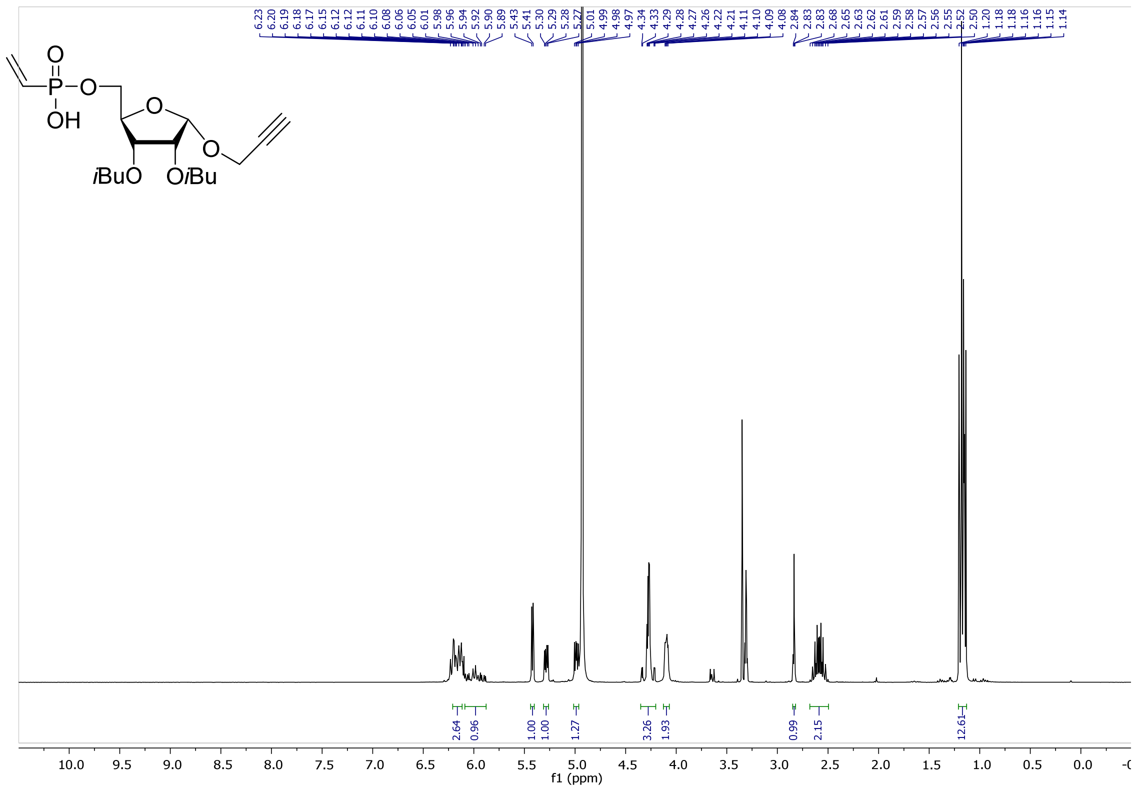

$^{13}$C NMR (**10**)

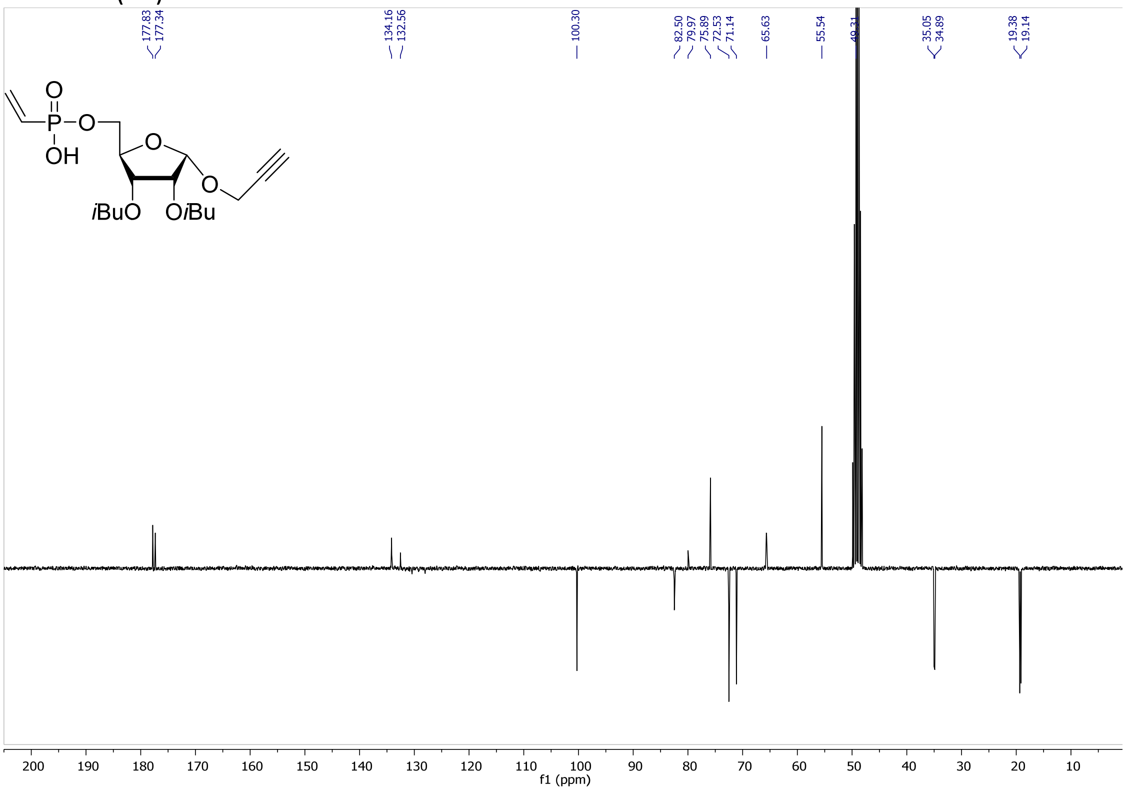

### $^{31}P$ NMR (**10**)

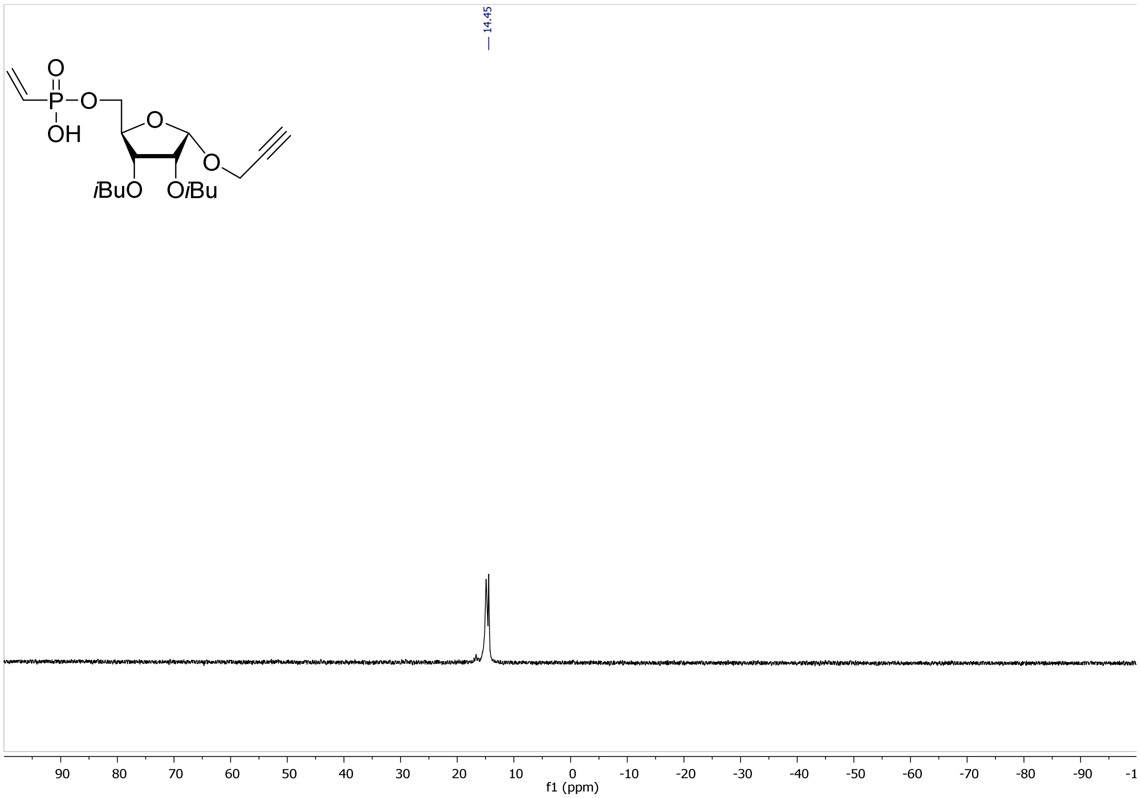

### $^{1}H$ NMR (**11**)

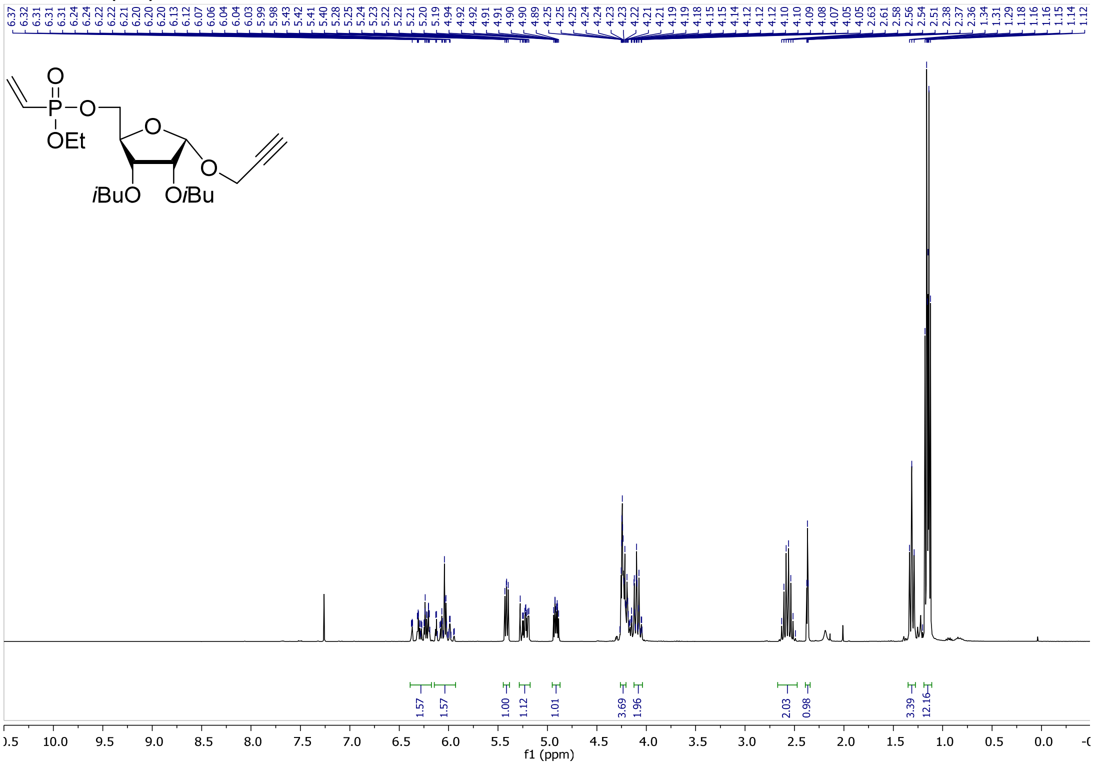

$^{13}C$ NMR (**11**)

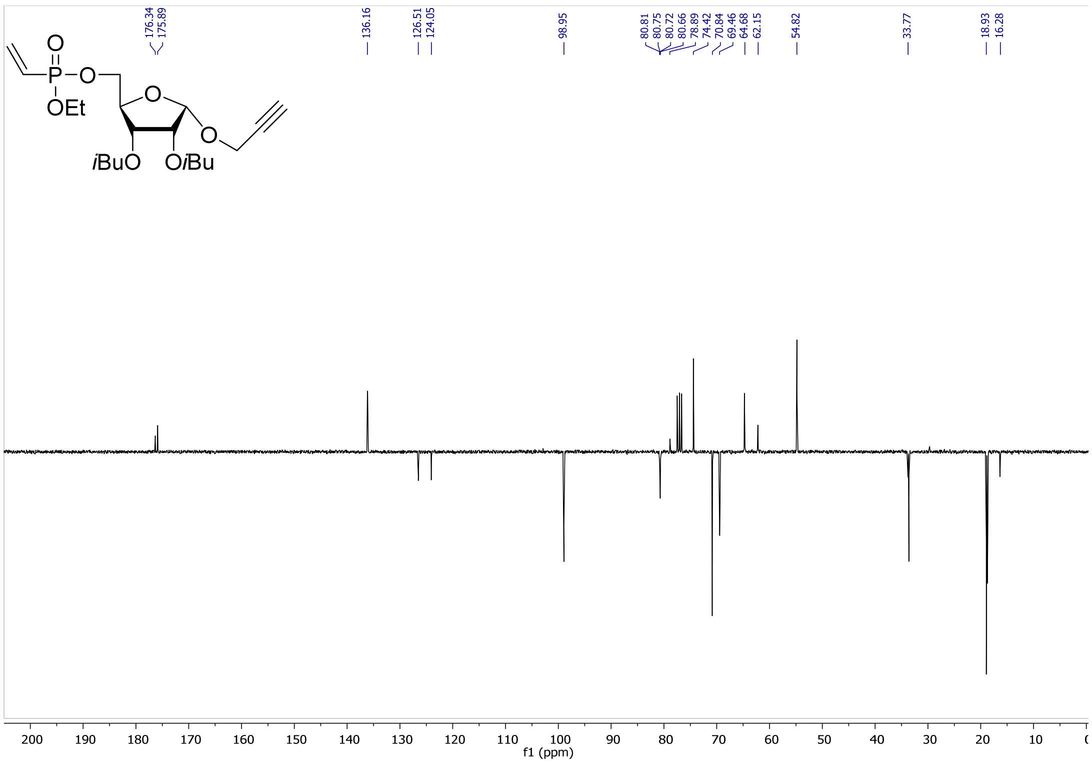

$^{31}P$ NMR (**11**)

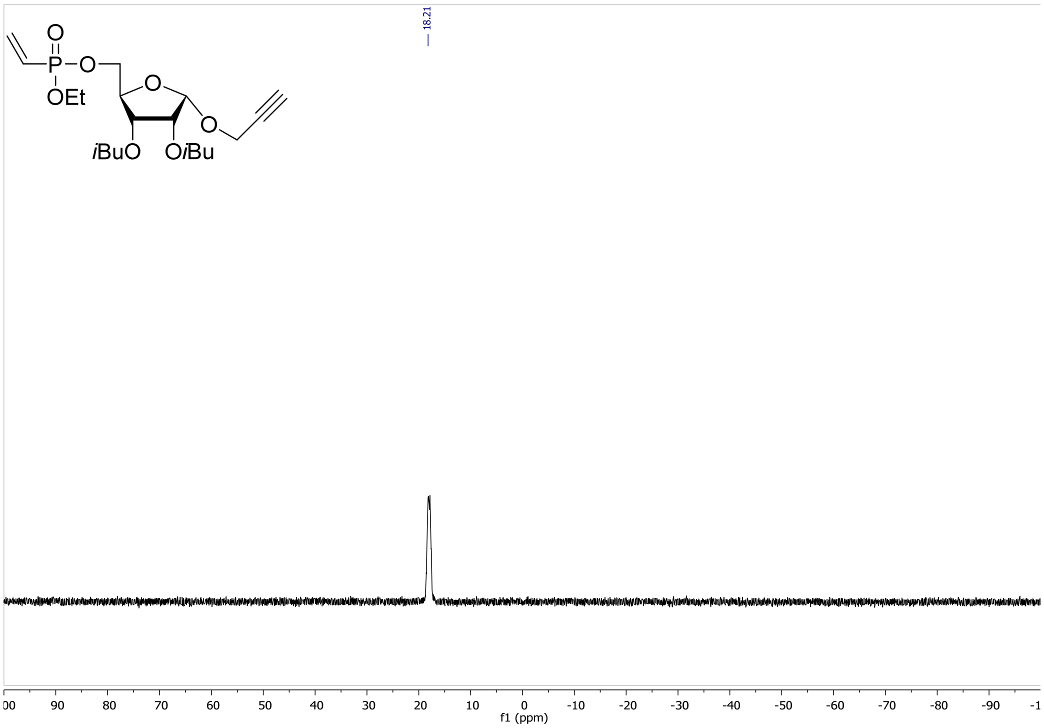

$^1$H NMR (**12**)

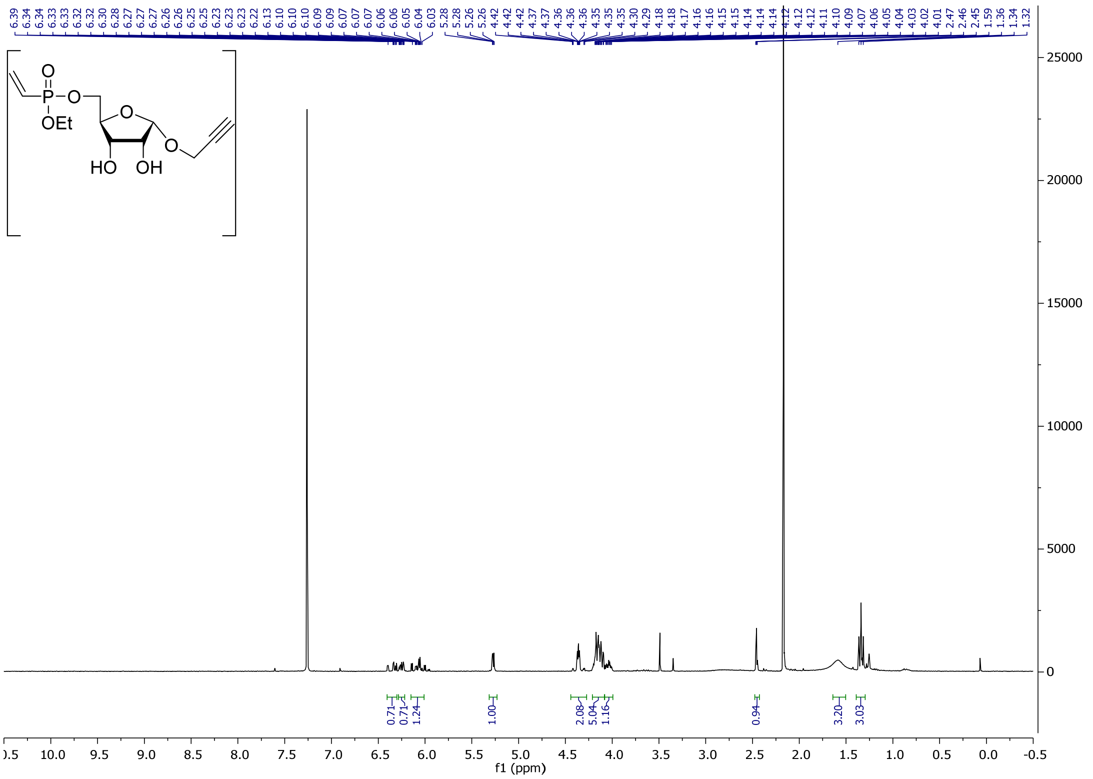

$^1$H NMR (**13**)

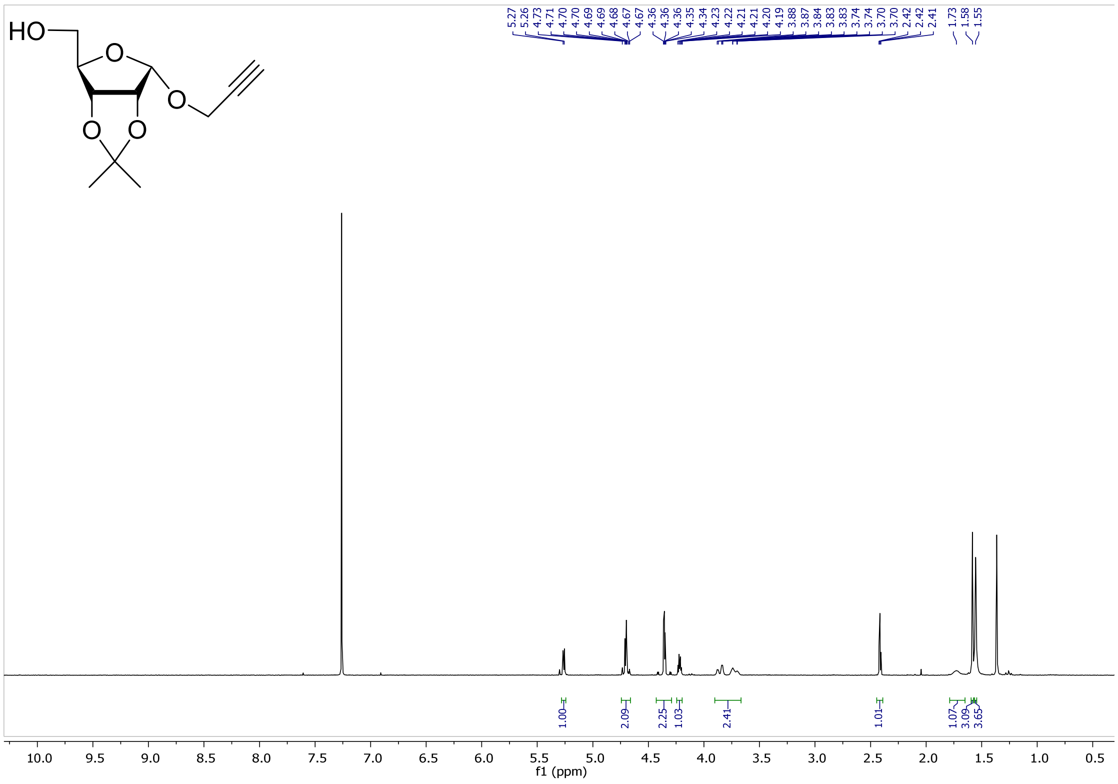

$^{13}C$ NMR (**13**)

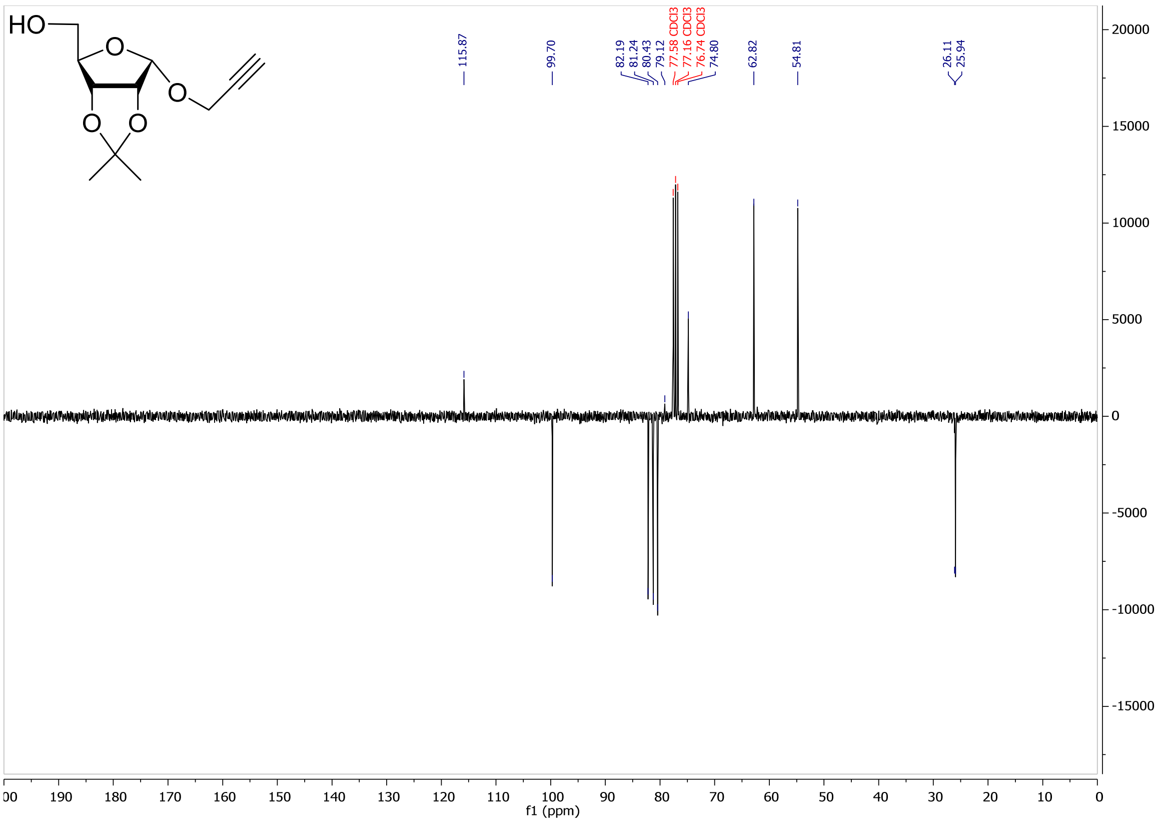

$^{1}H$ NMR (**14**)

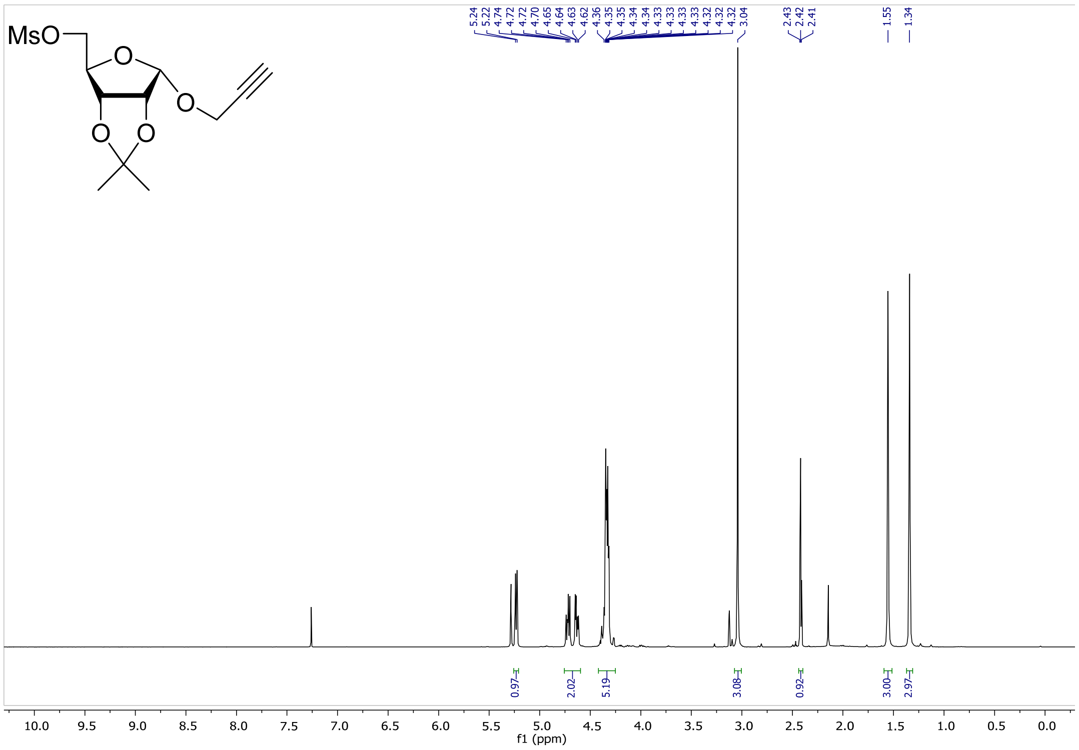

### $^{13}$C NMR (**14**)

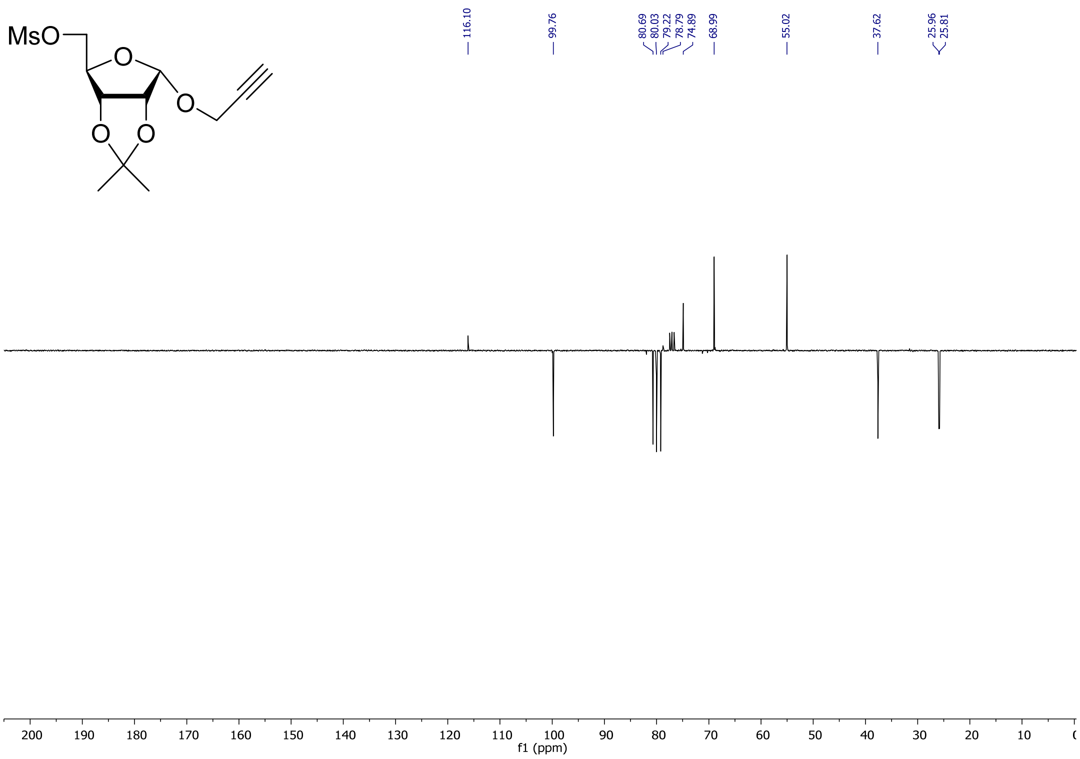

### $^{1}$H NMR (**15**)

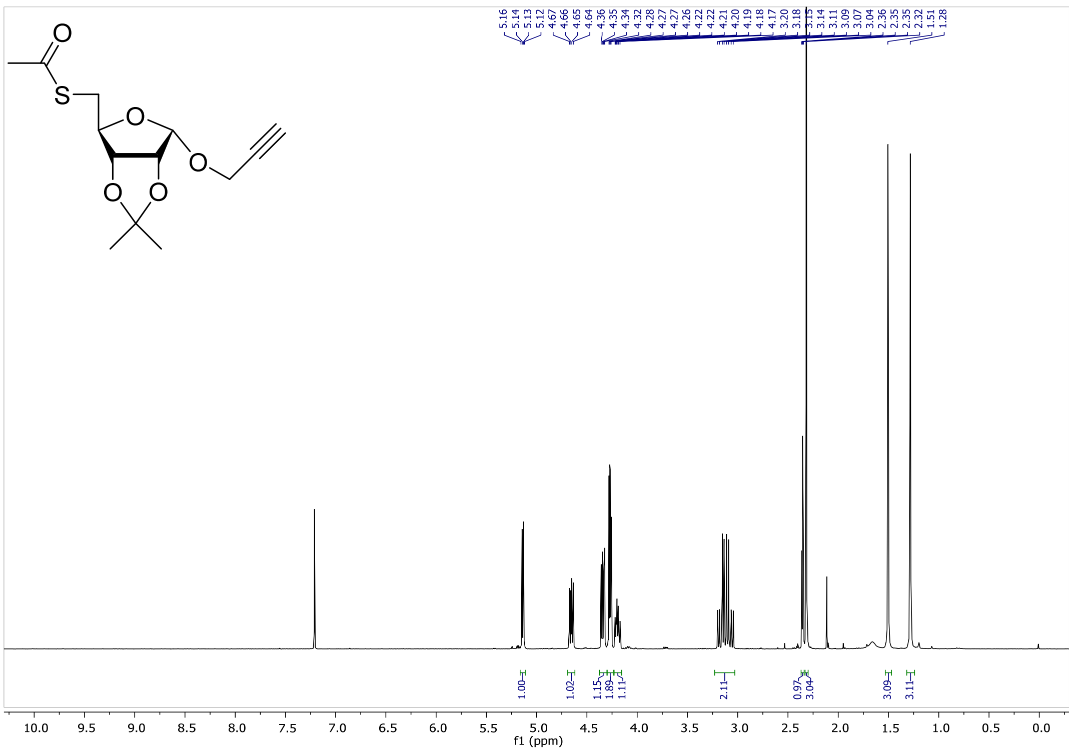

$^{13}C$ NMR (**15**)

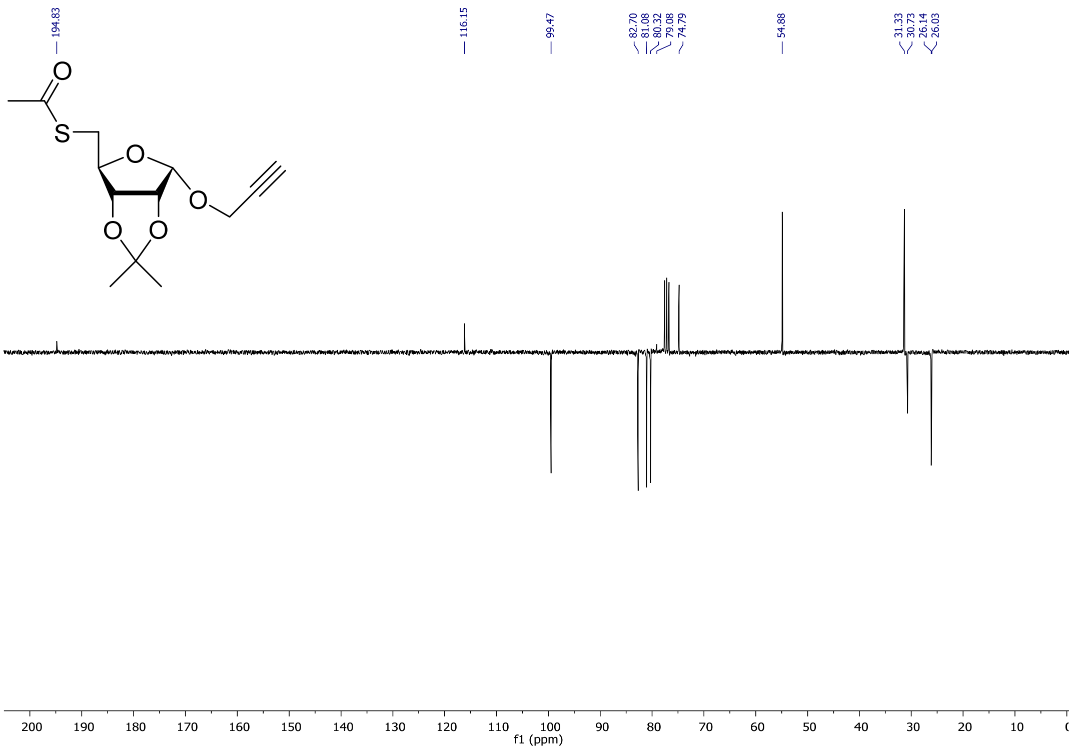

$^{1}H$ NMR (**16**)

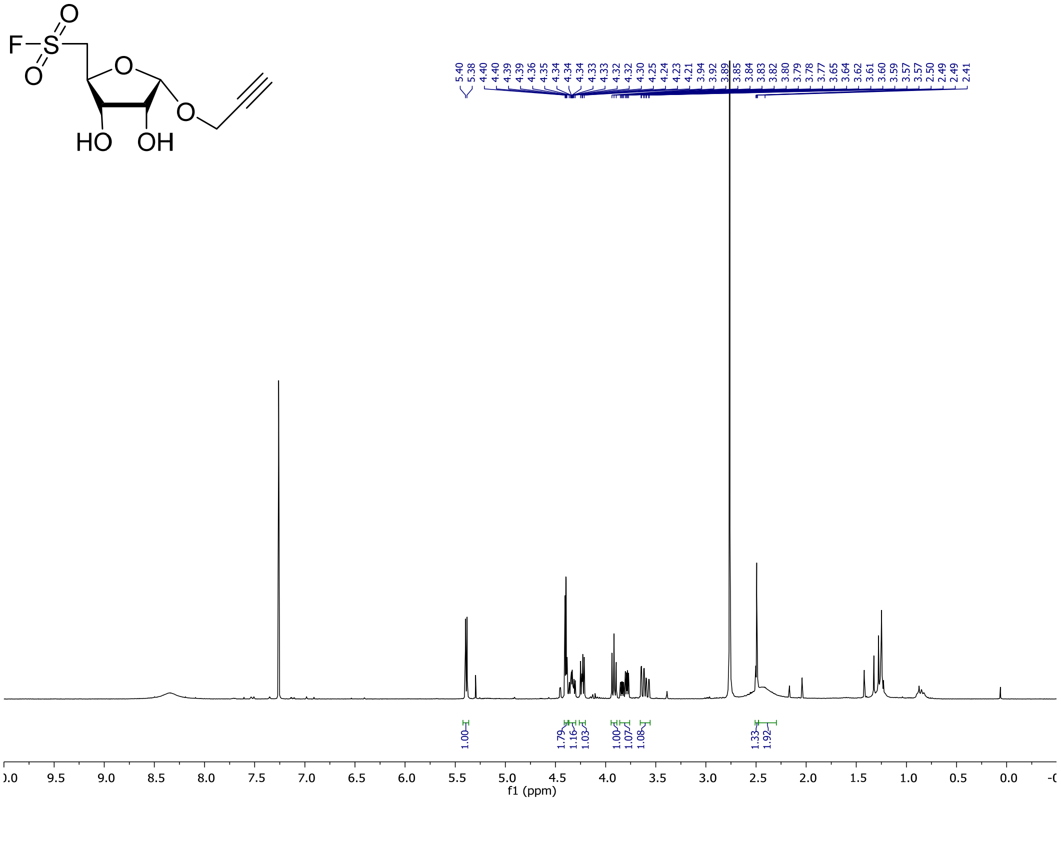

$^{13}$C NMR (**16**)

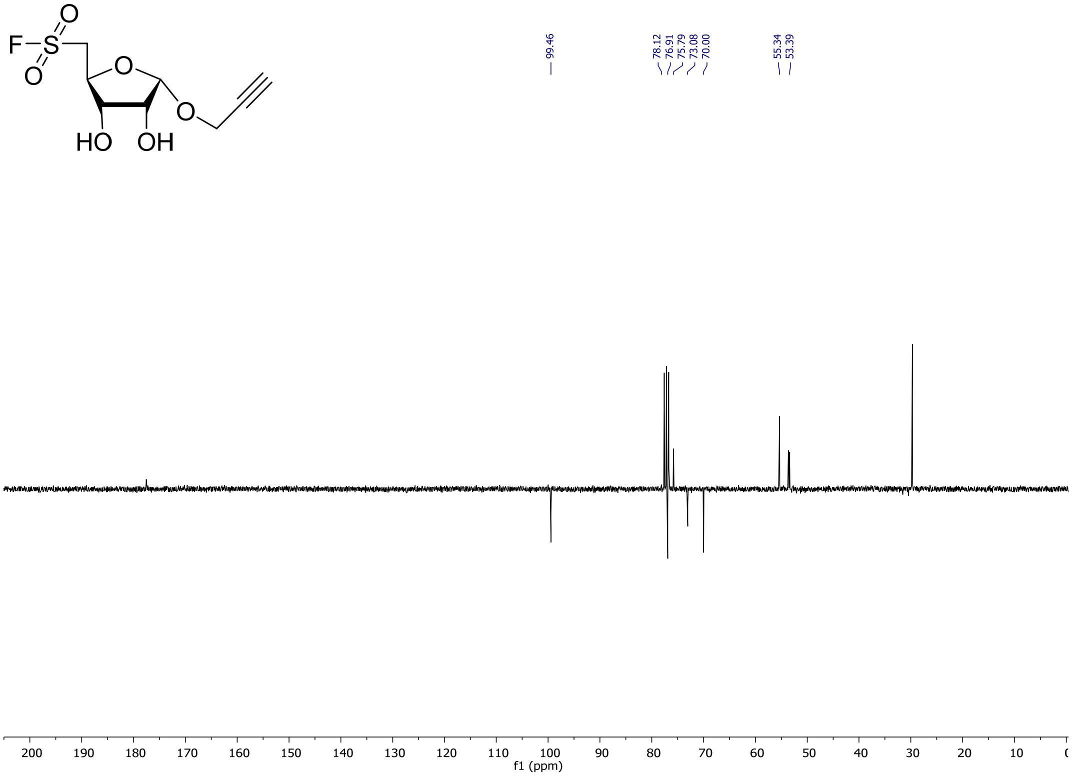

$^{19}$F NMR (**16**)

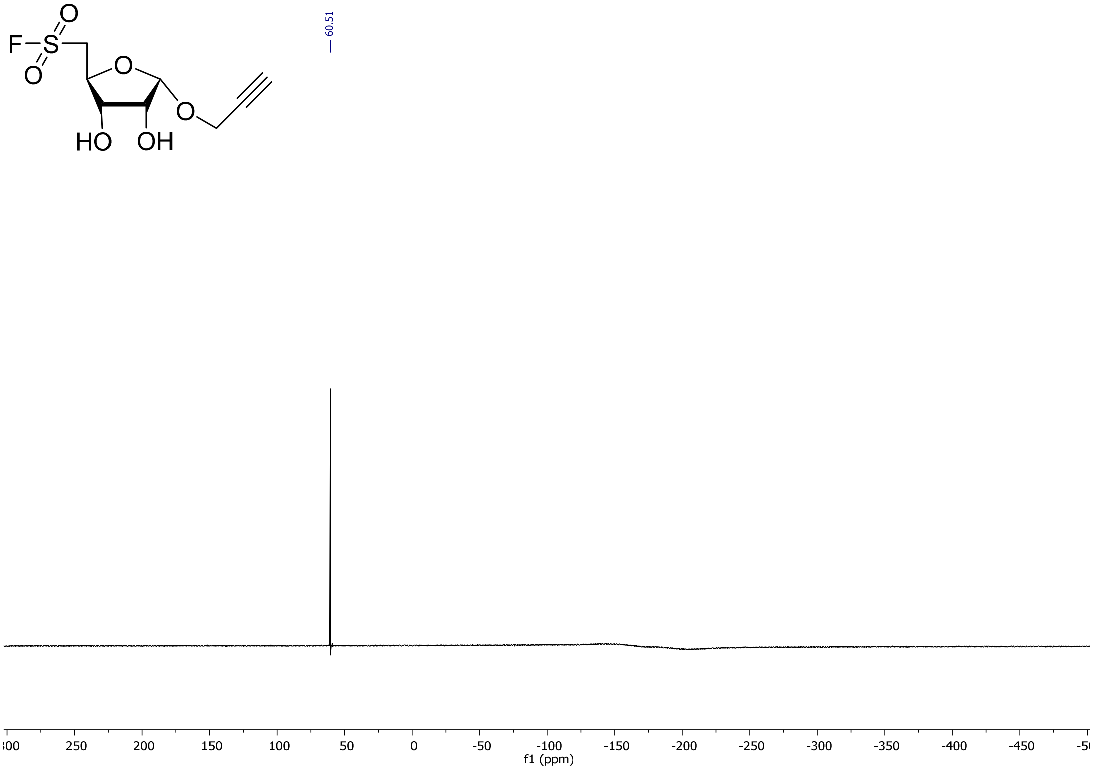

$^{1}$H NMR (**17**)

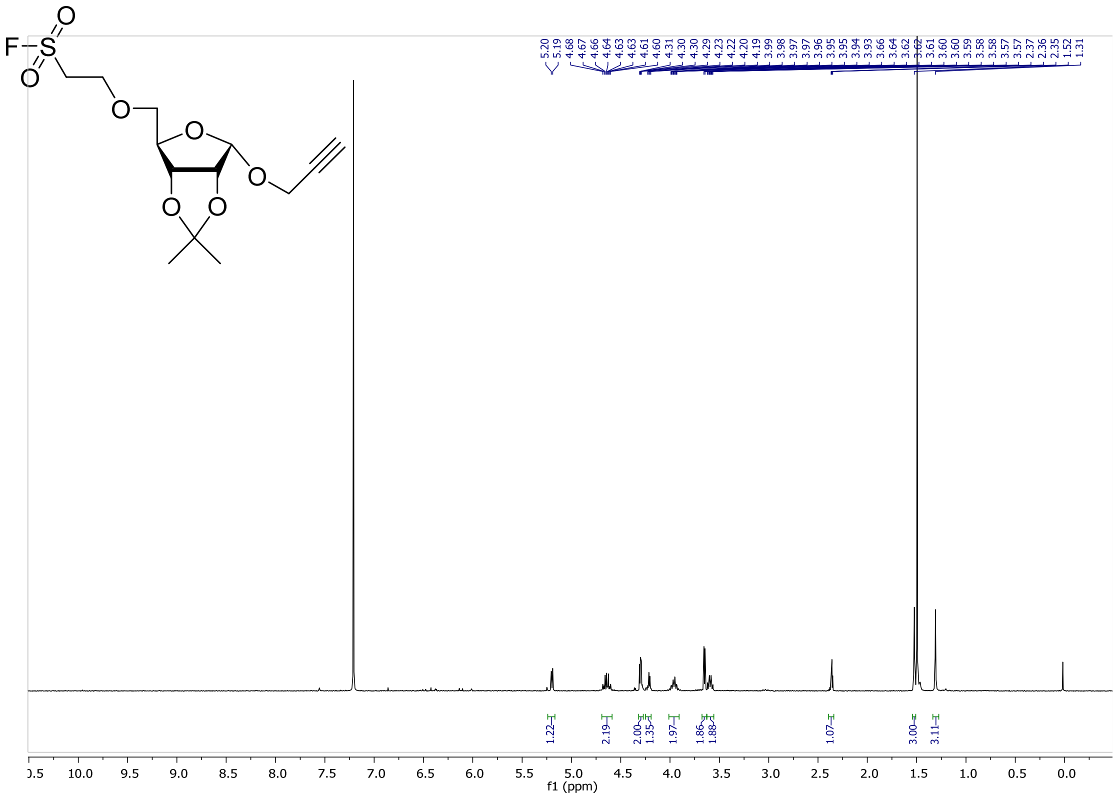

$^{13}$C NMR (**17**)

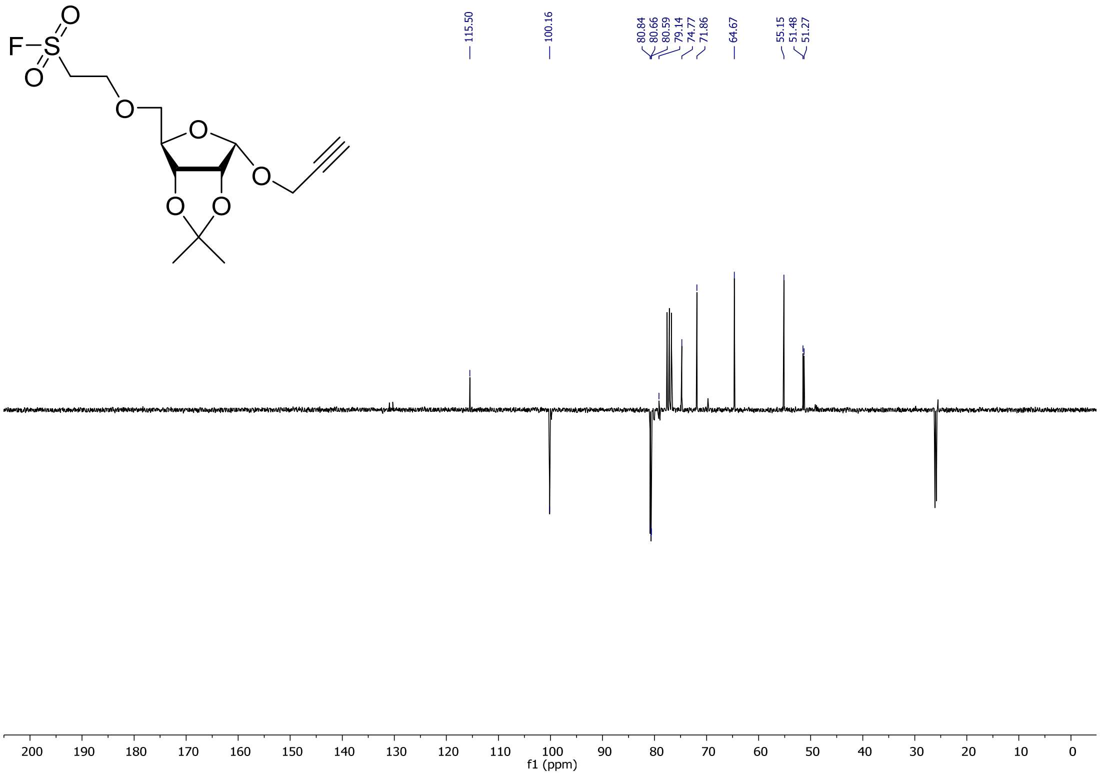

$^{19}$F NMR (**17**)

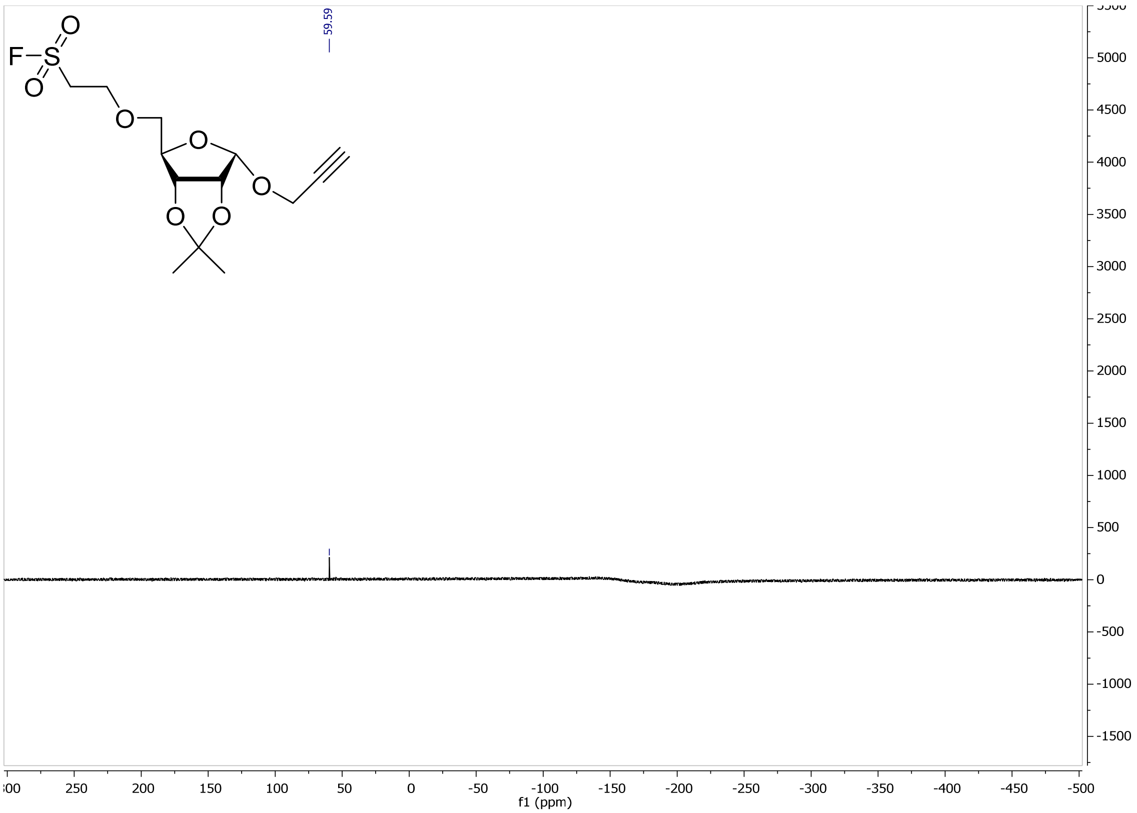

$^{1}$H NMR (**18**)

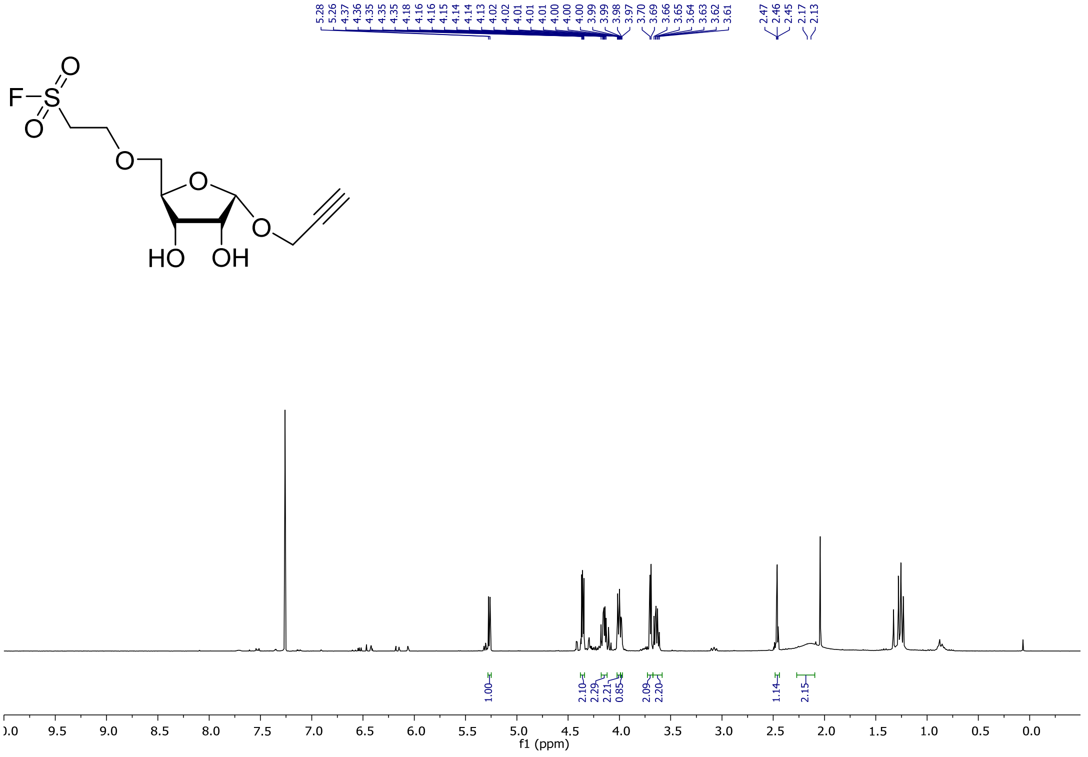

$^{19}$F NMR (**18**)

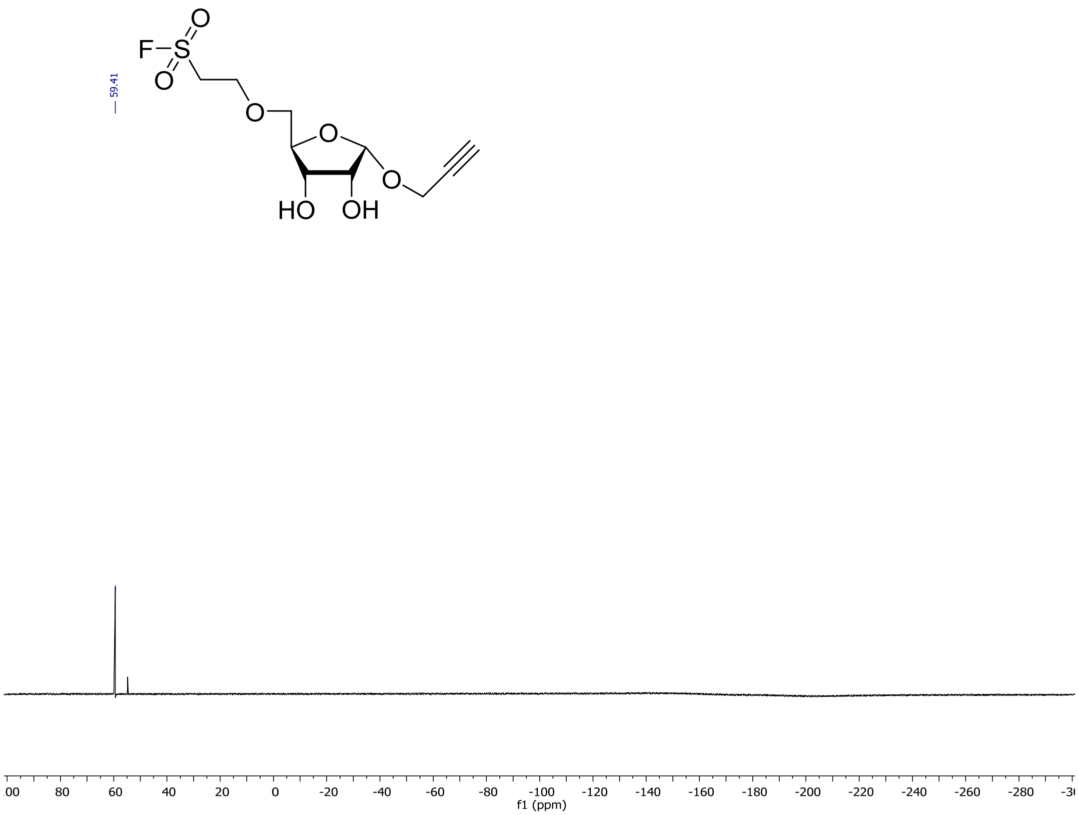

$^{1}$H NMR (**19**)

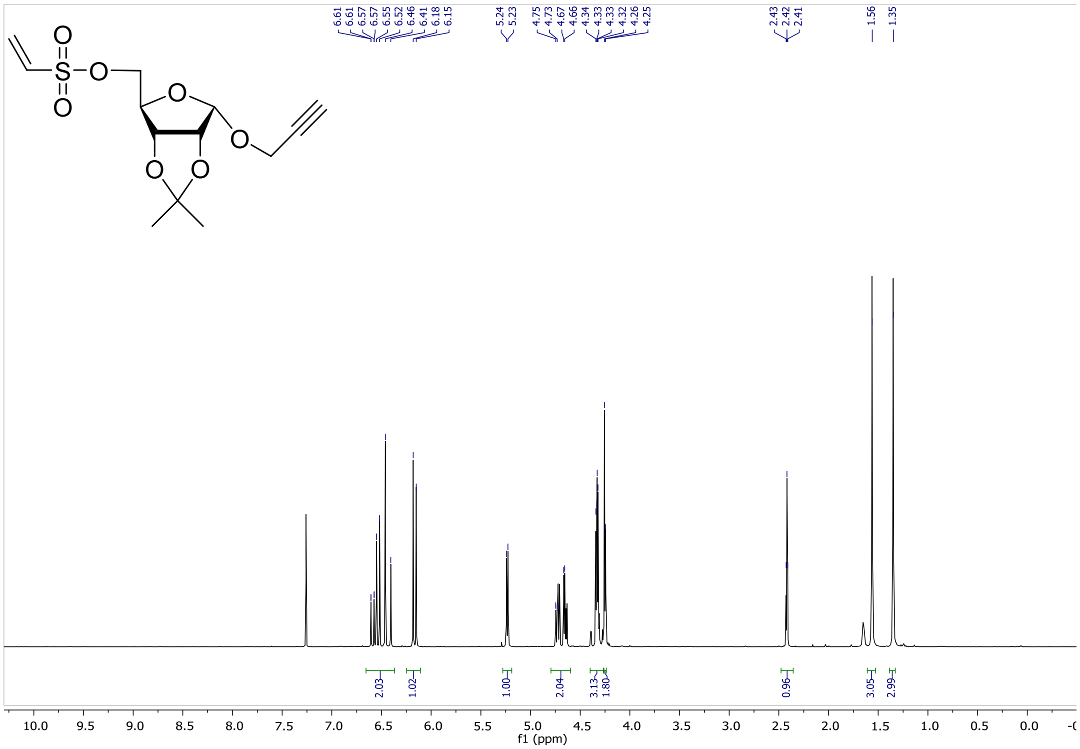

$^{13}C$ NMR (**19**)

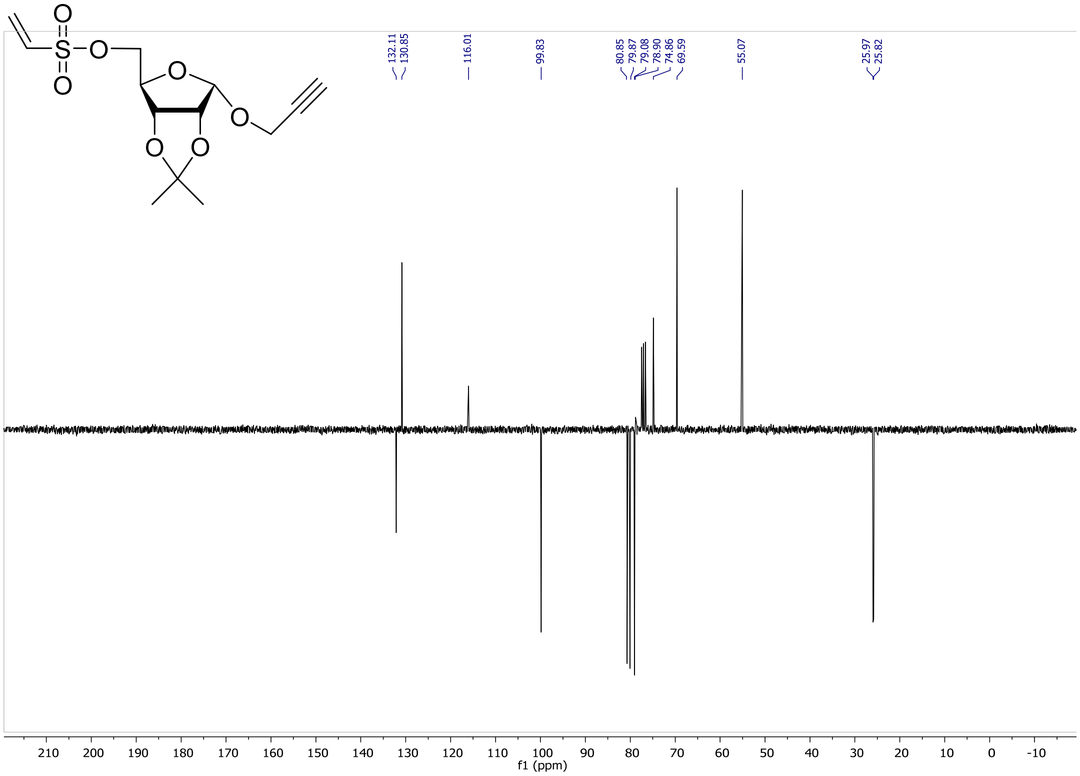

$^{1}H$ NMR (**20**)

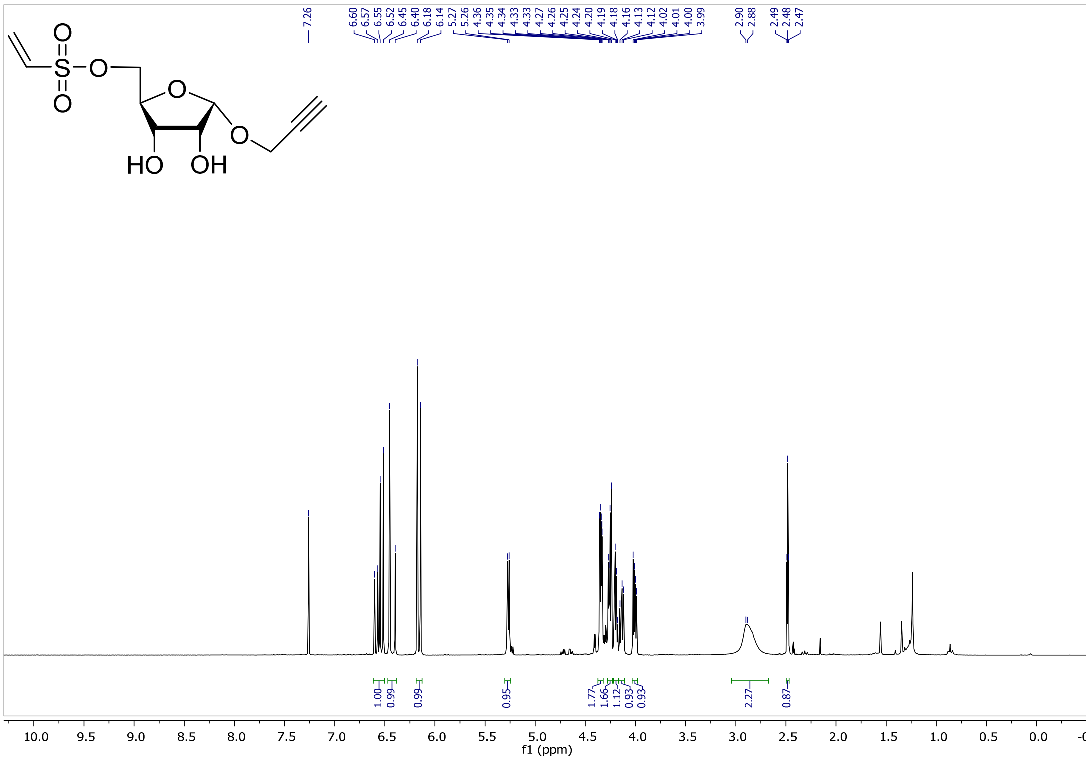

$^{13}$C NMR (**20**)

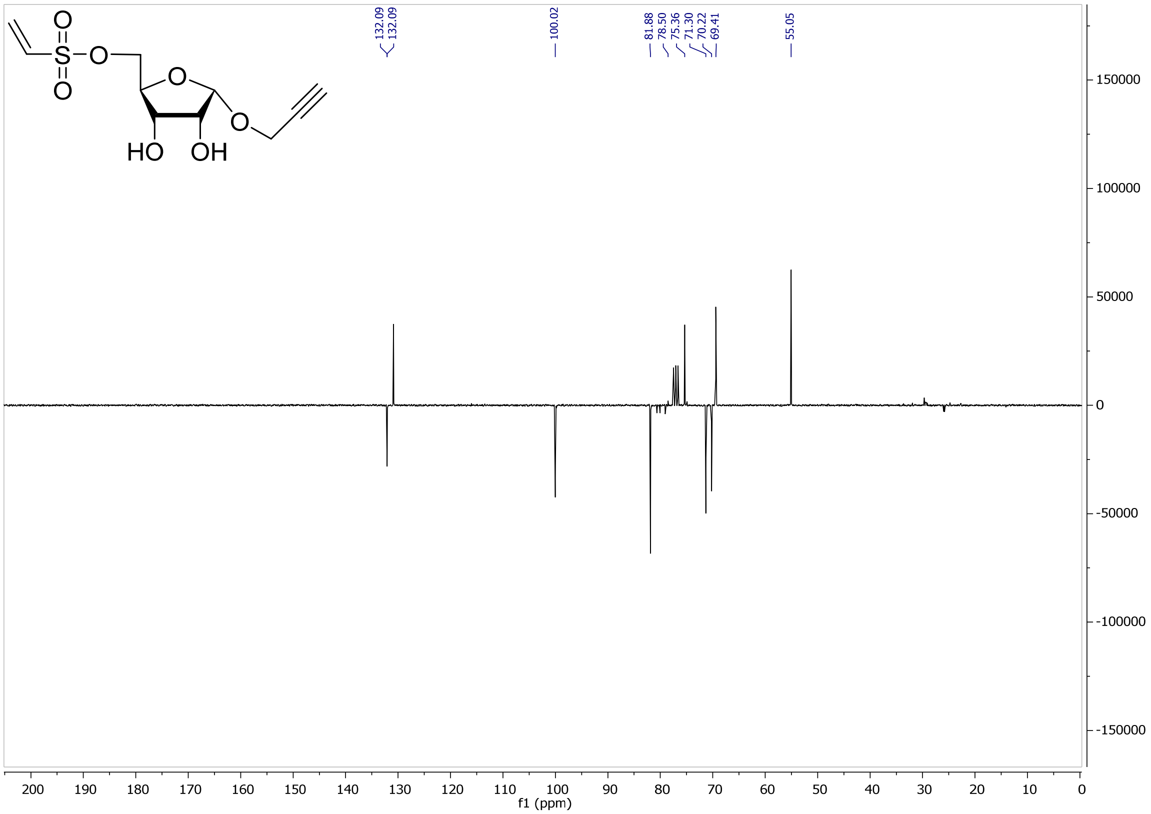

132.09
132.09
100.02
81.88
78.50
75.36
71.30
70.22
69.41
55.05
HO
OH
150000
100000
50000
0
-50000
-100000
-150000
200 190 180 170 160 150 140 130 120 110 100 90 80 70 60 50 40 30 20 10 0
f1 (ppm)
